# Drug reformulation for a neglected disease. The NANOHAT project to develop a safer more effective sleeping sickness drug

**DOI:** 10.1101/573329

**Authors:** Lisa Sanderson, Marcelo da Silva, Gayathri N. Sekhar, Rachel C. Brown, Hollie Burrell-Saward, Mehmet Fidanboylu, Bo Liu, Lea Ann Dailey, Cécile A. Dreiss, Chris Lorenz, Mark Christie, Shanta J. Persaud, Vanessa Yardley, Simon L Croft, Margarita Valero, Sarah A. Thomas

## Abstract

Human African trypanosomiasis (HAT or sleeping sickness) is caused by the parasite Trypanosoma brucei sspp. The disease has two stages, a haemolymphatic stage after the bite of an infected tsetse fly, followed by a central nervous system stage where the parasite penetrates the brain, causing death if untreated. Treatment is stage-specific, due to the blood-brain barrier, with less toxic drugs such as pentamidine used to treat stage 1. The objective of our research programme was to develop an intravenous formulation of pentamidine which increases CNS exposure by some 10-100 fold, leading to efficacy against a model of stage 2 HAT. This target candidate profile is in line with drugs for neglected diseases inititative recommendations. To do this, we evaluated the physicochemical and structural characteristics of formulations of pentamidine with Pluronic micelles (triblock-copolymers of polyethylene-oxide and polypropylene oxide), selected candidates for efficacy and toxicity evaluation *in vitro*, quantified pentamidine CNS delivery of a sub-set of formulations *in vitro* and *in vivo*, and progressed one pentamidine-Pluronic formulation for further evaluation using an *in vivo* single dose brain penetration study. Screening pentamidine against 40 CNS targets did not reveal any major neurotoxicity concerns, however, pentamidine had a high affinity for the imidazoline_2_ receptor. The reduction in insulin secretion in MIN6 β-cells by pentamidine maybe secondary to pentamidine-mediated activation of β-cell imidazoline receptors and impairment of cell viability. Pluronic F68 (0.01%w/v)-pentamidine formulation had a similar inhibitory effect on insulin secretion as pentamidine alone and an additive trypanocidal effect *in vitro*. However, all Pluronics tested (P85, P105 and F68) did not significantly enhance brain exposure of pentamidine. These results are relevant to further developing block-copolymers as nanocarriers, improving BBB drug penetration and understanding the side effects of pentamidine.

## Introduction

Human African trypanosomiasis (HAT or sleeping sickness) is a potentially fatal disease caused by the parasite *Trypanosoma brucei* sspp. Recent epidemiological studies in 30 of the 36 African countries listed as endemic for the disease indicate that, whilst the number of disease cases has been decreasing since 1990, there are still ~4,000 new infections/year, and ~15,000 cases worldwide [1][2]. Furthermore, there is a substantial unreported burden of HAT [3].

The disease has two stages – a haemolymphatic stage after the bite of an infected tsetse fly, followed by a central nervous system (CNS) stage when the parasite penetrates the brain, causing death if left untreated. The blood-brain barrier (BBB) makes the CNS stage difficult to treat because it prevents 99% of all known compounds from entering the brain, including most anti-HAT drugs[4][5][6][7]. Those that do enter the brain are toxic compounds, can have serious side effects, are complex to administer and/or are expensive. Pentamidine is a less toxic blood stage drug, which is known to treat early-late (transition) stage HAT[8], but cannot treat stage 2 disease as it does not sufficiently penetrate the BBB[7] and causes peripheral side effects (e.g. hypoglycaemia (incidence 5-40%) and diabetes mellitus (incidence: occasional but irreversible)[9] which preclude increasing the dose to overcome this limitation. Research has shown pentamidine has a limited ability to cross the BBB and reach the brain due to it physicochemical characteristics and its removal by the efflux transporters P-glycoprotein (Pgp) and multi-drug resistance associated protein (MRP) [7](Fig S1). Furthermore, transporters are considered essential in the mode of action of pentamidine against trypanosomes.

Poloxamers, with commercial trademark Pluronics^®^ (BASF) or Synperonics^®^ (CRODA), are triblock copolymers made of two poly(ethylene oxide) (PEO) blocks interspaced by a poly(propylene oxide) (PPO) block and follow the general basic formula: PEO_x_-PPO_y_-PEO_x_, where x and y are the size of PEO and PPO blocks, respectively (Table 1). In an aqueous environment and above the critical micelle concentration (CMC), the copolymers self-assemble into micelles, with the PEO chains forming a hydrophilic shell around a PPO hydrophobic core, within which lipophilic drugs can be solubilised, drug-free fraction decreased and circulation time increased [10]. A variety of Pluronic block copolymers differing in the lengths of the EO and PO blocks are available for formulation with pharmaceutical drugs. Importantly the size of the hydrophobic block affects micellization and drug solubilisation[11]. Furthermore, combining different Pluronics^®^ can enhance drug/micelle interactions and drug loading[12][13]. The PEO shell serves as a stabilizing layer between the hydrophobic core and the external medium, and prevents aggregation, plasma protein adsorption and opsonization and therefore recognition by the macrophages of the reticuloendothelial system [14]. Pluronic copolymers are also endowed with low cytotoxicity and weak immunogenicity in topical and systemic administration. Even though PEO–PPO–PEO materials are non-degradable, molecules with a molecular weight (MW) <7⍰kDa can be filtered by the kidney and cleared in urine[15] (Table 1). In addition, Pluronics are recognised pharmaceutical excipients listed in the US and British Pharmacopoeia so have an established safety profile.

**Table 1.** Pluronics used in this Study, with their Name, Block Composition, Hydrophilic-Lipophilic Balance (HLB) and General Formula. L, F, or P, Refers to Liquid, Flake, or Paste Physical Forms, respectively.

| Poloxamer | Pluronic | MW | Number of EO blocks | Number of PO blocks | HLB | Formula |
| --- | --- | --- | --- | --- | --- | --- |
| 235 | P85 | 4600 | 52.27 | 39.66 | 16 | EO<sub>26.13</sub>PO<sub>39.66</sub>EO<sub>26.13</sub> |
| 335 | P105 | 6500 | 73.86 | 56.03 | 15 | EO<sub>36.93</sub>PO<sub>56.03</sub>EO<sub>36.93</sub> |
| 188 | F68 | 8400 | 152.73 | 28.97 | 29 | EO<sub>76.37</sub>PO<sub>28.97</sub>EO<sub>76.37</sub> |
| 181 | L61 | 1950 | 4.55 | 31.03 | 3 | EO<sub>2</sub>PO<sub>30</sub>EO<sub>2</sub> |

Thus Pluronics have attracted a great deal of attention in pharmaceutical applications as drug solubilisers [14] or controlled drug-release agents[16][13][17]. Notably, Pluronic P85, P105, F68 and L61 have been shown to inhibit efflux transporters (including P-gp and MRP1-2) and have potential to enhance drug passage across the BBB [18] [19][20] [21] [22][23][24][16][25][26][27]. They have all been approved as cosmetic ingredients [15] with F68 having been utilized as a blood substitute component[28]. Transporter-targeting Pluronics^®^ (L61 and F127) have successfully completed a phase 2 clinical trial for the intravenous treatment of adenocarcinoma of the upper gastrointestinal tract [29][30]. Interestingly, F127-based amphotericin B-containing micelles have been shown to be highly effective in treating *Leishmania amazonensis*-infected BALB/c mice [31], demonstrating that Pluronics have potential beyond the traditional role of simple micellar vessels for drug encapsulation and longer circulation, but are also active agents with key biological functions [32].

In this Medical Research Council (MRC) developmental pathway funding scheme (DPFS) multi-disciplinary study our team developed a milestone driven progression strategy (Fig 1) in order to assess the potential of pentamidine-Pluronic formulations to effectively treat stage 2 disease, reduce the major known side effect of pentamidine on the pancreas and shorten the length of treatment required to treat stage 1 disease. It was anticipated that the benefits of this approach would be a combined pentamidine-Pluronic formulation which would provide a single therapeutic entity for safer, simpler and more cost-effective treatment of all HAT stages using an established drug with a known safety profile. Four Pluronics were selected for evaluation based on their block-copolymer architecture, established safety profile and known ability to inhibit Pgp. These were P85, P105, F68 and L61 (Table 1). An iterative approach was utilized as illustrated in Fig 1.

**Fig 1.**
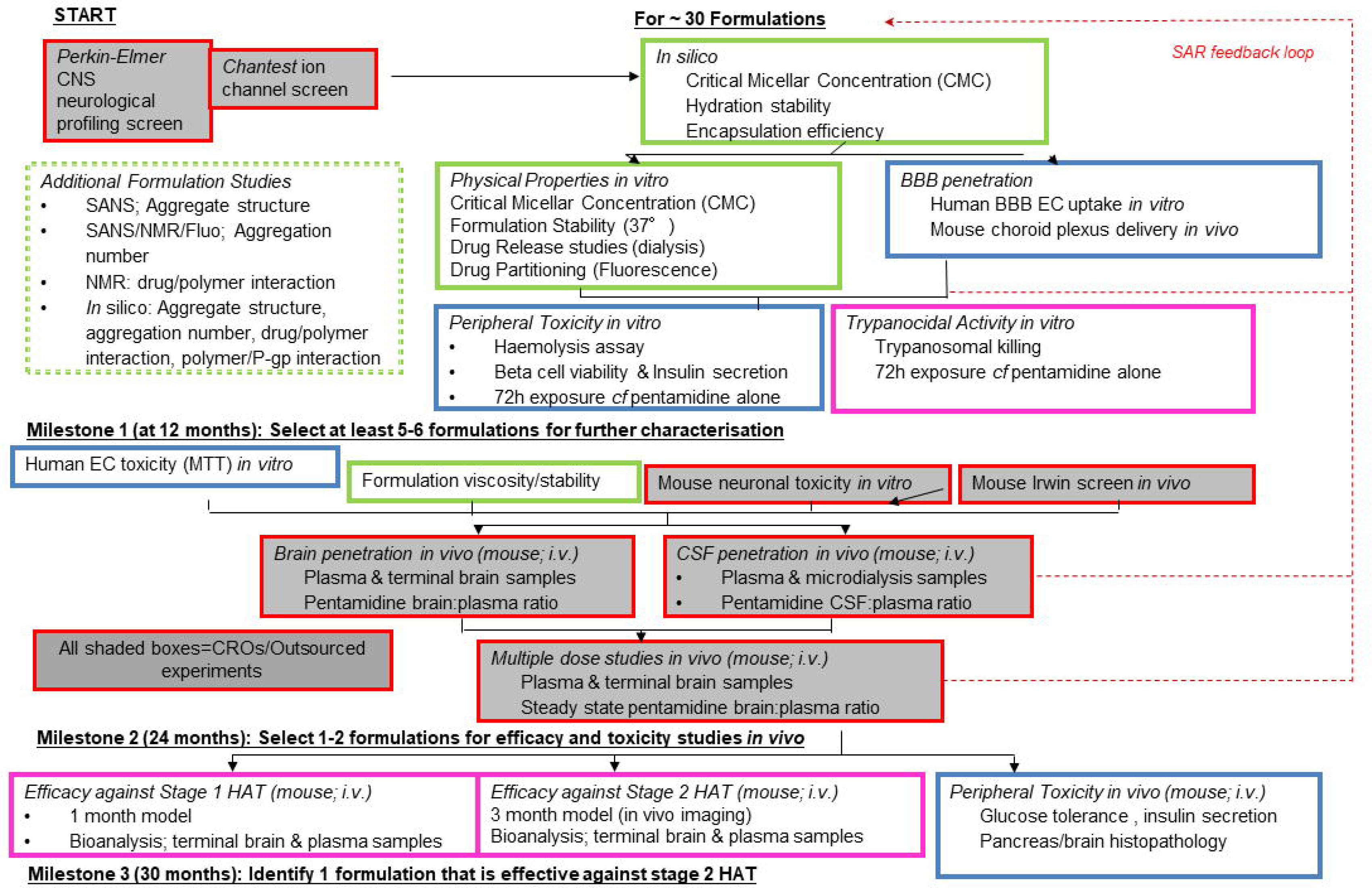
NANOHAT project progression strategy. We screened approximately 30 pentamidine/Pluronic^®^ formulations during this project using a rational, iterative approach. The three milestones were intended to ensure that the most appropriate formulations, on the basis of *in silico* and *in vitro* data, were taken forward to the *in vivo* pharmacokinetic studies and that the formulations with the greatest likelihood of success would be tested in the whole animal efficacy studies as outlined in the progression strategy.

## 2. Methods

### 2.1a Materials

Pentamidine (1,5-bis-4ρ-amidinophenoxypentane) isethionate salt (MW 592.68; 98% purity; catalogue number P0547) and Pluronic P105 (batch number BCBP8915V) were purchased from Sigma Aldrich (Poole, Dorset, UK). Pluronic P85 (mat 30085877 batch number: WPYE5378) was a kind donation from BASF plc (Cheshire, UK). Pluronic F68 (medical grade Catalogue number 2750016; batch numbers M7102 and MR29468) was purchased from MP Biomedicals, LLC (Illkirch Cedex, France). L61 was purchased from Aldrich (catalogue number 435422; batch number MKBH8737V).

### 2.1b Purity of excipients

All Pluronic formulations (F68, P105, P85 and L61) were tested for possible contamination due to the synthesis process and met specifications established by the US Pharmacopeia convention NF32 monograph for polaxamers [15], in that the EO, PO and p-dioxane in the Pluronic were below 1, 5 and 5 parts per million (ppm), respectively. This analysis was performed by an external specialist laboratory (Butterworth laboratories, Teddington, UK).

### 2.2 Evaluation of potential neurotoxicity of pentamidine

New toxicities may arise following pentamidine’s improved access to the CNS. The potential of pentamidine to cause neurotoxicity was evaluated by a brief review of the literature together with a neurological profiling screen and ion channel activity screens. The biological screens were performed by external specialist laboratories as described below.

#### 2.2a Neurological profiling screen

A CNS side effect panel was custom designed and binding assays performed by Perkin-Elmer Science Discovery Systems (Hanover MD 21076, USA). Testing was performed at a single concentration of 1 μM (100-times the trypanocidal concentration), with follow up concentration-response curves in any assay where there was greater than 70% inhibition to determine an inhibition constant (K_i_).

#### 2.2b Ion channel (hKir2.1) activity screens

The *in vitro* effects of pentamidine isethionate on cloned hKir2.1 potassium channels (encoded by the human KCNJ2 gene) responsible for the I_K1_, inwardly rectifying potassium current, were examined by ChantTest Corporation (Cleveland Ohio 44128, USA) to industry standards (Chantest FastPatch Assay; study number. 130827.DCC). Human epithelial kidney 293 (HEK293) cells (ATCC, Manassas VA USA) were stably transfected with the appropriate ion channel cDNA encoding the pore-forming channel unit. Cells were cultured in Dulbecco’s Modified Eagle Medium / Nutrient Mixture F-12 (D-MEM/F-12) supplemented with 10% fetal bovine serum, 100 U/mL penicillin G sodium, 100 μg/mL streptomycin sulphate and 500 μg/mL G418. Cultured cells were maintained in a tissue culture incubator set at 37°C in a humidified 95% air and 5% CO_2_ atmosphere. Pentamidine was dissolved in HEPES-buffered physiological saline containing 0.3% DMSO and sonicated (Model 2510/5510, Branson Ultrasonics, Danbury, CT) at room temperature for at least 20 minutes. A glass-lined 96 well compound plate was loaded with the appropriate amount of test (five different concentrations) and positive control (100μM BaCl_2_) solutions, and placed in the plate well of the QPatchHT (Sophion Bioscience A/S, Denmark). All experiments were performed at room temperature. Each cell acted as its own control. Vehicle was applied to naïve cells for a 5-10 minute exposure interval. The test solution applied for a minimum of three minutes via the QPatch robot pipetting system to naïve cells (n≥2, where n=the number of cells/concentration). Each solution exchange on the QPatch, performed in quadruplicate, consisted of a 5 μl exchange through the microfluidic flow channel, resulting in 100% replacement of the compound in the QPlate. Intracellular solution was loaded into the intracellular compartments of the QPlate planar electrode (130mM K-Asp, 5mM MgCl_2_, 5 mM EGTA, 4mM Tris-ATP and 10 mM HEPES). Cell suspension was pipetted into the extracellular compartments of the QPlate planar electrode.

Onset and steady state block of hKir2.1current was measured using a ramp protocol with fixed amplitudes (hyperpolarization: −110 mV, 200 ms duration, followed by a 1-second ramp from −110 mV to +50 mV) repeated at 10 s intervals from a holding potential of –70 mV. Current amplitude was measured at the end of the step to −110 mV. Leak current was calculated and subtracted from the total membrane current record.

### 2.3 Determination of the micellar aggregation properties of Pluronic

The CMC, micellar size and aggregation number were determined in different solvents, using a unique combination of light and neutron scattering and atomistic simulations. We also measured the partitioning of pentamidine isethionate in selected Pluronic and the *in vitro* release profile

#### 2.3a Preparation of solutions for physicochemical measurements

Unless stated, F68, P85, P105 or L61 were either dissolved in water (aqueous) or saline solution (0.9% w/v sodium chloride solution). Pluronic mixtures were also prepared either with a fixed mass ratio of 1:1 (F68-P105 or F68-P85) or in the case of L61, 0.01%. Samples were left to equilibrate for at least 3 hours prior to any measurement. Ultra-pure water (18.2 MΩ⋅cm - Millipore-filtered) was used throughout the experiments.

#### 2.3b: Phase behaviour

In this study, L61 alone and in mixtures with one or two other Pluronics in both water (aqueous) and saline mediums were visually assessed from 20°C to 50°C in 5°C steps, plus 37°C, to assess the impact of mixtures on L61 cloud point (24°C for a 1% solution) [33].

#### 2.3c CMC determination by fluorescence spectroscopy

The CMC determines thermodynamic stability of the micelles during dilution of the drug delivery system in body fluids[17][11]. Furthermore, CMC is an important parameter in view of the biological response modifying effects of Pluronic block copolymers since it is needed to determine the maximum achievable concentration of the polymer single chains (“unimers”) [20]. For measurement of the CMC, pyrene (Sigma catalogue number 82648; pyrene puriss p.a. for fluorescence, ≥99%) was used as a probe. A stock solution of pyrene in acetone (1.7×10^−2^ M) was initially prepared. A 35 μL aliquot of this solution was placed in a 100 mL volumetric flask and the solvent was evaporated to air. The residue was then dissolved in either ultra-pure water (18.2 MΩ⋅cm - Millipore-filtered) or 0.9% w/v sodium chloride solution, resulting in a final concentration of pyrene of 6×10^−6^ M. These solutions were then subsequently used as the solvent for the polymer solutions. Stock solutions of each Pluronic in water and saline solution were prepared. An appropriate aliquot of these solutions were dissolved in the pyrene/H_2_O or pyrene/saline solution. Solutions of different polymer concentration were obtained by diluting the stock polymer solution with the appropriate solvent. Mixed samples of two Pluronics were also prepared either with a fixed ratio of 1:1 or containing 0.01% L61. Samples were left to equilibrate for at least 3 hours prior to the experiment.

The fluorescence emission spectra were recorded on a Cary Eclipse fluorescence spectrophotometer (Varian, Oxford, UK) with λ_exc_ = 340 nm. For the CMC, fluorescence intensities at 373, 384, 393 nm and, when it appeared, also at the excimer band centred at 490 nm, were measured. For each polymer, the critical aggregation concentration value was determined by using the intensity of the best resolved peak. At least two repeats were performed for each sample. Measurements were performed at 20°C and 37°C.

#### 2.3d Stability testing

The purpose of stability testing is to check whether pentamidine becomes altered with time under the influence of a variety of environmental factors such as temperature, humidity and light (Climatic zone IV, 30°C and 65-75% relative humidity) [34].

In our initial 7 day assessment we also considered interaction of pentamidine with Pluronic as product-related factors may also influence its quality. A 5% or more change in initial content of pentamidine was considered significant. Pentamidine concentration at day 0, 10 and 7 was assessed by NMR.

A Bruker Advance 400 MHz spectrometer was used for recording the one-dimensional (1D) 1H NMR. Solutions of PTI, PTI/P85, PTI/P105 and PTI/F68 were prepared in D_2_O (≥99.85% in deuterated component). Data were collected at days 0, 1 and 7. Samples were stored in amber NMR tubes at 37°C.

#### 2.3e Partition coefficient determination

The partitioning coefficient, *P*, determines the fraction of drug incorporated into the micelle and provides thermodynamic characterization for the stability of the drug-micelle complex during dilution within the body fluids[17][11]. The partition coefficient of pentamidine in the micellar core and bulk solvent, as described by Kabanov and co-workers [11], was measured for F68, P105 and mixtures of P105 and F68 (1:1), in both saline and aqueous solutions and at 20°C and 37°C

The partition coefficient was obtained following the procedure described by Kabanov and co-workers [11], which is described briefly below. The partition coefficient is defined as:

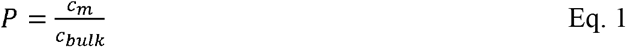

where *c*_*m*_ is the PTI concentration in the micelle core and *c*_*bulk*_ is the PTI concentration in the bulk solvent.

The partition coefficient *P* can be obtained from[11]:

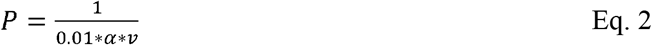

where, *v* (cm^3^/g), the partial molar specific volume, is defined as:

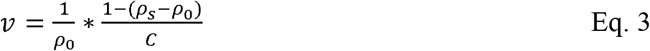

with ρ_0_ (g⋅cm^−3^) the solvent density, ρ (g⋅cm^−3^) the density of the solution and *C* the polymer concentration (g⋅mL^−1^).

α is defined as the angular coefficient of the plot 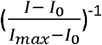 vs. (Pluronic concentration – cac)^−1^, where *I* is the fluorescence intensity of the solution, I_0_ is the intensity with no Pluronic present and *I*_*max*_ is the intensity at saturating concentration of the Pluronic, cac the critical aggregation concentration.

For the measurement of pentamidine isethionate salt (PTI) partition coefficient, a PTI 1×10^−6^ M in both water and saline solutions were initially prepared. These solutions were then subsequently used as the solvent for the polymer solutions and the preparation followed a similar method as for the CMC measurements. Samples were left to equilibrate for at least 3 hours prior to the experiment.

The fluorescence emission spectra were recorded on a Cary Eclipse fluorescence spectrophotometer (Varian, Oxford, UK) with λ_exc_ = **260nm**, for pentamidine. The fluorescence emission intensity at ca 340 nm was followed.

#### 2.3f Drug release

Solutions of Pluronic (1% F68 and 1% P105) with 10 mM PTI and PTI alone in water (2 mL) were loaded into 2 mL mini-dialysis tubes with 1 kDa molecular weight cut-off (GE Healthcare Bio-sciences Corp. USA). The tube was immersed in a 200 mL closed Duran flask which was placed in a water bath at 37°C for the duration of the experiment. Aliquots were collected from the immersion water (ultra-pure water (18.2 MΩ⋅cm - Millipore-filtered) in the flask every 30 min for the first 2 hours, every hour for the next 5 hours and then once more after 1 week. At the end of the experiment, an aliquot was collected from the dialysis cell. PTI concentrations were determined by UV spectroscopy (wavelength 260 mm).

The data was fitted to Ritger-Peppas model[35].

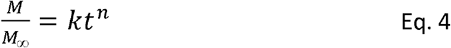

Where *M* and *M*_□∞_ are the cumulative amounts of drug released at time *t* and at infinite time, respectively; k, the reaction constant, *t* the time, *n*, the diffusional exponent describing the type of regime type: n=1, case II transport, n=0.5, Fickian diffusion, 0.5<n<1 non-Fickian diffusion.

#### 2.3g Dynamic light scattering (DLS)

Dynamic light-scattering (DLS) were performed with a photon correlation spectrometer Malvern Zetasizer Nano with a laser wavelength of 633nm. For obtaining the reduced scattered intensity, toluene was used as the standard and the increment in the refractive index, ∂n/∂c, was assumed to be independent on the temperature and taken as 0.133 ± 0.002 mL⋅g^−1^ [36]. The samples, of concentrations ranging between 1 to 5% w/v, were filtered prior to the measurements by 0.22 μm Millex syringe PVDF filters onto semi-micro glass cells. The temperature of the sample was controlled with 0.1°C accuracy by the built-in Peltier in the cell compartment. Size distributions were obtained for each sample from the analysis of the intensity autocorrelation function, which was performed with the Zetasizer software in the high resolution mode to distinguish overlapping distributions.

#### 2.3h Small-Angle Neutron Scattering (SANS)

The architecture of the nanocarriers was measured by SANS. The aggregation number (*N*_*agg*_) and radius micellar size, including volume of core and shell region, correlates directly with are relevant to properties such as drug loading encapsulation efficiency, stability, half-life and hence circulation time[37].

Small-angle neutron scattering experiments were performed on the LOQ instrument at ISIS pulsed neutron source (ISIS, Rutherford-Appleton Laboratory, STFC, Didcot, Oxford). LOQ uses incident wavelengths from 2.2 to 10.0 Å, sorted by time-of-flight, with a fixed sample-detector distance of 4.1 m, which provided a range of scattering vectors (q) from 0.009 to 0.29 Å^−1^.

The samples used in the SANS experiments were prepared in D_2_O to optimise the contrast with the protonated polymer. The samples were placed in clean disc-shaped quartz cells (Hellma) of 1 and 2 mm path length and the measurements were carried out at 25°C and 37°C. All scattering data were first normalised for sample transmission and then background-corrected using a quartz cell filled with D_2_O (this process also removes the inherent instrumental background arising from vacuum and windows) and finally corrected for the linearity and efficiency of the detector response using instrument-specific software package. The data were then converted to the differential scattering cross-sections (in absolute units of cm^−1^) using the standard procedures at ISIS [38,39].

The curves were fitted to a core-shell sphere model combined with a structure factor for hard spheres, implemented in the Sasview software [40]. The initial fitting assumptions were a dry PPO core, i.e., the scattering length density (SLD) for the core was set to the SLD of PPO (3.44×10^−7^ Å^−2^), the solvent as D_2_O (SLD of 6.38×10^−6^ Å^−2^) and the shell as dry PEO (SLD 6.4×10^−7^ Å^−2^). The SLD of core and solvent were kept fixed while the shell was left to vary as a high level of hydration of PEO is expected, as a result, the returned SLD should be an intermediate value between D_2_O and PEO. After obtaining a stable set of parameters, the SLD of the core was also let free to vary. Input values for the core radius and shell thickness were based on hydrodynamic radius values obtained by DLS. A term to compensate for polydispersity was included for Pluronic (around 0.20), as well as a structure factor (S(q)), corresponding to a hard sphere model, in order to account for intermicellar interactions.

Assuming that the shell SLD is a linear combination of the EO SLD and D_2_O SLD, i.e., the densities are additive, it is possible to calculate the solvent volume fraction in the shell using the following expression:

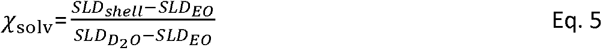

The amount of solvent molecules in the shell than can be obtained from the ratios of the shell volume by the solvent molecular volume weighted by the solvent volume fraction:

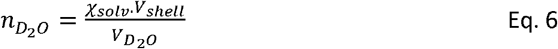

Furthermore, the total micelles volume is the sum of both Pluronics and D_2_O molecular volumes weighted by their respective volume fractions. Therefore, combining the total micelle volume (shell + core) minus the solvent contribution, it is possible to obtain the amount of Pluronic molecules present in the micelle and calculate the aggregation number of the micelles.

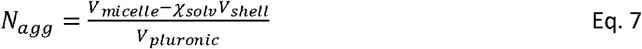

The molecular volume of the Pluronic was approximated from the Pluronic molar volume (v_pluronic_) in water.

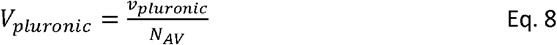

#### 2.3i Simulations of Pluronic self-assembly and pentamidine encapsulation

During the course of this project, we worked to develop a model of the Pluronic and pentamidine systems that would allow us to simulate the self-assembly of the polymers and the encapsulation of the drugs. In order to simulate the timescales and system sizes required to study these systems, we utilized a coarse-grain approach; dissipative particle dynamics (DPD)[41]. This method has been used to study Pluronic before and has been shown to represent expected phenomena well. So we used the simulation parameters from [42].

### 2.4 Evaluation of potential peripheral toxicity of pentamidine ± Pluronic

The toxicity of pentamidine in the presence of the Pluronic was explored using a variety of assays. These included haemolysis, insulin secretion and membrane integrity assays.

#### 2.4a Haemolytic effects on human blood samples

The proposed route of administration for our Pluronic formulations with pentamidine was intravenous, hence the propensity for Pluronic to lyse red blood cells was studied using a haemolytic assay.

##### 2.4ai Chemicals

Cyanmethaemoglobin (CMH) reagent, haemoglobin standard, Ca^2+^/Mg^2+^ free Dulbecco’s Phosphate Buffered Saline (DPBS), DMSO, and Triton X-100 were purchased from Sigma-Aldrich, Dorset, UK.

##### 2.4aii Research Donor Blood

Healthy volunteer blood samples were drawn under the guidelines of the Research Ethics Committee South East London REC 4 (10/H0807/99). Blood was collected in BD vacutainer tubes containing lithium heparin as anticoagulant. Blood samples from two healthy volunteers were collected and kept separate.

##### 2.4aiii Haemolysis Assay

The assay is based on the protocol detailed by [43]. In order to determine the total blood haemoglobin, the CMH method was used to map a standard curve based on the absorption wavelength at 540 nm. Nine calibration standards were produced by preparing a stock solution of 5 mg/ml (standard 1) using haemoglobin standards in CMH reagent and serially diluting it to produce further standards of concentrations 2.5, 1, 0.80, 0.40, 0.20, 0.10, 0.05, and 0.025 mg/ml. 10% Triton X-100 (v/v) was prepared using distilled water to be used as the positive control. A 0.05% solution of DMSO was prepared using distilled water and used as one of the negative controls. 0.9% saline was another negative control. Pluronic samples were prepared in 0.9% saline. The test concentrations of Pluronic (P85, P105 or F68) used were 0.01%, 0.1%, 0.25%, 1% or 5%.

A 5 ml vial of whole blood was centrifuged at 800 g for 15 minutes at room temperature. The supernatant was removed and the remainder was used to determine the plasma free haemoglobin (PFH) concentration.

200 μl × 2 of each of the calibration standards prepared were then transferred onto the 96-well plate and 200 μl × 4 of CMH reagent (control 1). Total blood haemoglobin (TBH) was then prepared by adding 20 μl of the whole blood with 5 ml of CMH reagent. 200 μl × 6 of TBH was transferred to the plate. 6 other wells were filled with 100 μl of plasma to which 100 μl of CMH reagent was added. After shaking it gently for a few minutes, the absorbance was read at 540 nm. Once the total haemoglobin concentration was adjusted to 10 mg/ml using Ca^2+^/Mg^2+^ free DPBS, 20 μl × 3 of the blank (control 2 0.9% saline), positive control, negative control, or Pluronic samples were added to Eppendorf tubes. 160 μl of Ca^2+^/Mg^2+^ free DPBS and 20 μl of whole blood were then added to each tube except for one. 20 μl of Ca^2+^/Mg^2+^ free DPBS was added instead. These served as control 3 without any blood enabling us to determine any interactions from Pluronic with the assay. These tubes were incubated for 3 hours ± 15 minutes in a temperature-controlled shaker (THERMOstar, BMG labtech, Offenburg, Germany) at 37 °C and 120 rpm. After the incubation, these were centrifuged at 800 × g for 15 minutes at room temperature. 100 μl of the test samples and the controls were transferred to the 96-well plate. 100 μl of CMH reagent was added to all these before measuring the absorbance spectrophotometrically at 540 nm. Using the calibration curve mapped earlier, the haemoglobin concentration in each of the wells was determined. The dilution factor of 18 was also considered when calculating the haemoglobin concentration. Haemoglobin concentration was converted to percentage haemolysis compared to the negative control 0.05% DMSO. A significant increase compared to control at the 5% level was taken as positively haemolytic.

#### 2.4b Membrane integrity/cytotoxicity

Method is described as part of 2.5b and 2.6.

#### 2.4c Insulin secretion and beta-cell viability

All tissue culture reagents were purchased from Sigma Aldrich (Poole, Dorset, UK).

Peripheral toxicity of pentamidine/Pluronic formulations to the endocrine pancreas was evaluated by quantifying β-cell viability and insulin secretion from the mouse MIN6 β-cell line [44].

MIN6 β-cells were maintained in culture at 37°C (95% air/5% CO_2_) in DMEM supplemented with 10% fetal bovine serum, 2mM L-glutamine and 100U.ml^−1^/0.1mg/ml^−1^ penicillin / streptomycin, with a change of medium every 3 days. Cell were trypsinised (0.1% trypsin, 0.02% EDTA) when approximately 70% confluent and seeded into 96 well plates at a density of 3×10^4^ cells/well. After a 24 hour culture period to allow cells to adhere, the wells were washed with PBS and cells were pre-incubated for 2 hours in DMEM supplemented with 2mM glucose after which the medium was replaced with DMEM supplemented with Pluronic, pentamidine and Pluronic/pentamidine solutions.

The following formulations were evaluated: F68/PTI, P85/PTI and P105/PTI with Pluronic concentrations of 0, 0.01, 0.025, 0.1 and 0.5% w/v and PTI concentrations of 0, 1, 10 and 100 μM (20 formulations in total, including controls, Pluronic only, PTI only and solvent only were used). The cells were incubated under each treatment condition for 24 hours and then evaluated for their capacity to secrete insulin in response to 30 minute stimulation in the presence of 10 μM forskolin and 100 μM IBMX. Insulin secretion was measured by RIA [45]. The effect of the formulations on β-cell viability was assessed by determining the access of trypan blue to the cell interior, indicative of a compromised plasma membrane[46].

### 2.5 Blood-brain barrier studies

#### 2.5a Radiochemicals

[^3^H(G)]pentamidine (specific activity, 31.9 Ci/mmol; concentration, 10.74 μg/ml; radiochemical purity, 99.4%; MW 342.64) was custom synthesized and [^14^C(U)]sucrose (specific activity, 536 mCi/mmol; concentration, 67.07 μg/ml; radiochemical purity, 98.7%) was purchased from Moravek Biochemicals, California, USA.

#### 2.5b *In vitro* permeability assays

Several *in vitro* BBB models were evaluated for this study including Caco2 (permeability format), hCMEC-D3 (accumulation format), bEnd-3 (accumulation format) and MDCK-MDR (accumulation format) cell lines, before selecting the MDR1-MDCK cells (permeability format) as the most appropriate tool to address our objectives. MDR1-MDCK cells originate from transfection of Madin-Darby canine kidney (MDCK) cells with the MDR1 gene, the gene encoding for the human efflux protein, P-glycoprotein (P-gp). Using MDR1-MDCK cells avoids the complexities of multiple transporters by focusing specifically on P-gp.

##### 2.5bi Preparation of formulation

1% (w/v) stock solutions of each Pluronic and 10 mM pentamidine isethionate were prepared in Hank’s Balanced Salt Solution (HBSS) containing 25 mM HEPES and 4.45 mM glucose, at pH 7.4. These were further diluted to give final concentrations of 0.01, 0.1 or 0.5% (w/v) Pluronic containing 10 μM pentamidine isethionate. Formulations were stored at room temperature for 2-4 days prior to use.

##### 2.5bii *In vitro* permeability assays

MDR1-MDCK cells (NIH, Rockville, MD, USA) were maintained and permeability assays were performed at both Cyprotex (Macclesfield, Cheshire, UK) and King’s College London. Analysis was by UPLC-MS/MS or liquid scintillation counting as appropriate.

Transmission electron microscopy confirmed appropriate cell morphology of a monolayer with microvilli on the apical membrane and Western blot confirmed expression of P-gp (data not shown). 3.4 × 10^5^ cells/cm^2^ were seeded on Multiscreen™ plates with 0.4 μ polycarbonate Isopore™ membranes (Millipore, MA, USA) in DMEM/High glucose (Sigma-Aldrich, UK, D6429) media containing 1% Non-Essential Amino Acids and 10% foetal calf serum (both from Sigma-Aldrich, UK). Plates were maintained at 37°C/5% CO_2_ for 4 days before use. On the day of the assay, DMEM was removed and both the apical and basolateral surfaces of the cell monolayer were washed twice with transport medium consisting of HBSS containing 25 mM HEPES and 4.45 mM glucose, (pH 7.40; 37°C). Plates were incubated for 40 minutes at 37°C/5% CO_2_ to stabilize physiological conditions. Transport buffer was removed from the apical or basolateral chamber and replaced with the formulation to be tested. Samples were taken from the apical and basolateral compartments after 1 hour of incubation at 37 °C/5% CO_2._ Samples, including the test formulation added to the apical chamber at t=0 were analysed at Cyprotex using UPLC-MS-MS method to quantify the pentamidine isethionate content or were analysed for radioactivity using a Tricarb 2900TR liquid scintillation counter.

##### 2.5biii UPLC-MS/MS

Quantification of pentamidine isethionate was carried out on an ABSciex API5500 QTrap triple quadrupole mass spectrometer coupled to an Agilent 1290 Infinity UPLC system. A 10μl sample from the apical or basolateral chamber was diluted with 80 μl buffer and 160 μl methanol and centrifuged for 20 minutes at 1,400 × g. After centrifugation 90 µl of this was removed and diluted with 10 µl of the internal deuterated standard to give a final concentration of 10 nM. This was injected onto an Acquity™ HSS T3 (1.8 µm) column 2.1 × 50 mm (Waters Ltd, Herts, UK), equilibrated at 70 °C. Separation was carried out by gradient elution using a mixture of 0.1% formic acid in H_2_O (solvent A) and 0.1% formic acid in acetonitrile (solvent B), at a flow rate of 600 µL/min. Upon injection (10 μl), the mobile phase was held at initial conditions of 100% solvent A for 0.05 minutes, the concentration of solvent B was then increased to 95% at 1.00 minute post-injection using a linear gradient and held for a further 0.40 minutes. The mobile phase was then returned to 100% solvent A at 1.41 minutes and held for a further 0.39 minutes. The eluent was analysed by MS/MS under positive ion electrospray mode and the multiple reaction monitoring transitions for pentamidine and deuterated pentamidine were 341.174 to 119.984 m/z and 345.212 to 120.028 m/z respectively. Calibration curves were prepared using pentamidine and the internal standard over a concentration range of 0.0078 to 12 µM. The column was washed with a weak wash: 9:1 H_2_O: MeCN and strong wash: 4:3:3 MeOH: isopropyl alcohol: acetone + 1% acetic acid between samples.

### 2.6 Sensitivity of MDCK-MDR cells to Pluronic

Permeability assay with 5 μCi (0.9 μM) [^14^C(U)]sucrose alone in the presence of varying concentrations of Pluronic was used to assess the effect of Pluronic on monolayer integrity. Sucrose is a paracellular permeability marker and therefore any effects of Pluronic will lead to increased diffusion of [^14^C(U)]sucrose from apical to basolateral chambers as a result of compromised BBB integrity. The assay was carried out for an hour in the apical to basolateral direction only and the Papp of radiolabelled sucrose determined at the end of the assay. Control for the assay was HBSS buffer alone.

### 2.7 In situ perfusions

The *in situ* brain/choroid plexus perfusion method for examination of the distribution of molecules into the brain and CSF is an established technique in the rat, guinea-pig and mouse [47][6][48]. It allows the passage of slowing moving molecules across the blood-brain and blood-CSF barriers to be examined and quantified in brain, capillary endothelial cells and choroid plexus tissue for perfusion periods up to 30 minutes.

#### 2.7a Preparation of formulation

All formulations were prepared on the day of experiment at a Pluronic concentration of 0.1, 1.0 or 5% (w/v) using artificial plasma as a diluent. The artificial plasma consisted of a modified Krebs-Henseleit mammalian Ringer solution containing; 117 mM NaCl, 4.7 mM KCl, 2.5 mM CaCl_2_, 1.2 mM MgSO_4_, 24.8 mM NaHCO_3_,1.2 mM KH_2_PO_4_, 39 g dextran, 1 g/L of bovine serum albumin and 10mM glucose. [^3^H(G)]pentamidine was added to give a final concentration of 157nM (equivalent to 5 μCi/ml). All formulations were stirred at room temperature for at least 1 hour to allow any chemical interactions and micelle formation to stabilize.

#### 2.7b Animal studies

Adult male BALB/c mice were purchased from Harlan UK Ltd (Oxon, UK). All animals were maintained under standard temperature/lighting conditions and given food and water *ad libitum*. Only mice above 23g in weight were used for experiments which were carried out within the framework of the Animals Scientific Procedures Act (1986) and Amendment Regulations 2012 and with consideration to the ARRIVE guidelines. The study was approved by the King’s College London Animal Welfare and Ethical Review Body.

#### 2.7c In situ perfusions

[^3^H(G)]pentamidine formulations were delivered to the brain using an in situ brain perfusion technique as previously described [6]. Briefly, mice were anaesthetized (mixture of 2 mg/Kg Domitor™/150 mg/Kg ketamine administered via the intraperitoneal route) and heparinized (100 U ip.). Oxygenated artificial plasma (described above) at 37°C was pumped via a 25 gauge cannula into the left ventricle of the heart, with the right atrium severed to prevent recirculation. Pumps were calibrated to deliver an overall flow rate of 5 ml/min from the cannula. [^3^H(G)]pentamidine formulations (maintained at room temperature) were fed into the flow line from a dual syringe infusion pump (Harvard Apparatus, UK), at a rate of 0.5 ml/min such that the formulation was diluted 1/10 immediately prior to entering the heart. 11 μM [^14^C(U)]sucrose in artificial plasma (equivalent to 5 μCi/ml) was simultaneously fed into the flow line from a second identical syringe using the same pump set at 0.5 ml/min (equivalent to 1.1 μM or 0.5 μCi/ml entering the heart from the cannula). The perfusion was terminated at 10 minutes or 30 minutes, and the brain was sectioned as previously described [49]. Samples taken were those known to be invaded by parasites during second stage sleeping sickness and/or those which control mechanisms that are disrupted by the disease such as the sleep/wake cycle[6]. All samples were solubilized with 0.5 ml Solvable™ (PerkinElmer Life and Analytical Sciences, Buckinghamshire, UK) for 48 hours. Scintillation fluid (3.5 ml Luma Safe, PerkinElmer Life and Analytical Sciences) was added and radioactivity (^3^H and ^14^C) was counted on a Packard Tri-Carb2900TR scintillation counter in dual-label mode.

#### 2.7d Capillary depletion

After the required samples were taken, the remaining brain tissue was homogenized and analyzed by the capillary depletion method described by Thomas & Segal [50]. This method uses dextran density gradient centrifugation to produce a vascular endothelial cell-enriched pellet and a brain parenchyma-containing supernatant from homogenized brain tissue. Thus the partitioning of a drug between the endothelial cells and the post-vascular brain parenchyma can be examined [51]. Briefly, the brain tissue was weighed and homogenized in a Dounce homogenizer with 3 ml/g capillary depletion buffer [51] and 4 ml/g 26% dextran. The homogenate was separated by centrifugation at 5,400g and 4°C for 15 minutes. Homogenate, pellet and supernatant samples were solubilized and counted for radioactivity using the method described above.

#### 2.7e Expression of results

The radioactivity (either ^3^H or ^14^C) present in tissue samples (dpm/g) was expressed as a percentage of that measured in the artificial plasma (dpm/ml) and was termed R_TISSUE_%, as previously described [49]. Where stated, measurements for [^3^H(G)]pentamidine were corrected for the contribution of drug present in the vascular space by subtraction of the R % for [^14^C(U)]sucrose from the R_TISSUE_% of [^3^H(G)]pentamidine and these corrected values were termed R_CORR TISSUE_%.

### 2.8 Pharmacokinetic brain distribution experiments

#### 2.8a *in vivo* pharmacokinetic experiments with [^3^H(G)]pentamidine

Formulations containing 0.025% F68 with 8 μM [^3^H(G)]pentamidine, 0.5% F68 with 8 μM [^3^H(G)]pentamidine and 8 μM [^3^H(G)]pentamidine alone were prepared in 0.9% sterile saline and allowed to equilibrate at room temperature for at least 1 hour before use. A 200 μl bolus of the formulation to be tested (equivalent to 15 μCi [^3^H(G)]pentamidine) was administered to mice via the tail vein. At 2 hours post-injection, mice were exsanguinated via the right atrium of the heart into a heparinised syringe then perfused for 2.5 minutes with [^14^C(U)]sucrose (1.1 μM, 0.5 μCi/ml) via the left ventricle, (all mice were anaesthetised with Domitor™/ketamine and heparinised 20 minutes prior to exsanguination). Whole blood samples were immediately centrifuged for 15 minutes at 5,400 × g to remove red blood cells and the resulting plasma was placed on ice. A CSF sample was taken from the cisterna magna, the IVth ventricle choroid plexus and pituitary gland were collected and the brain was sectioned into right brain and left brain (both comprising frontal cortex and caudate putamen), cerebellum and midbrain (including pons and hypothalamus). The remaining brain (including occipital cortex and hippocampus) was used for capillary depletion analysis and all brain, circumventricular organs (CVO) and plasma samples were solubilized and subjected to dual label (^3^H/^14^C) scintillation counting as previously described

#### 2.8b *in vivo* pharmacokinetic experiments with pentamidine isethionate

Adult female CD1 mice (20-25g) were injected intravenously with pentamidine isethionate (4 mg/kg in 0.9% physiological saline) in the absence and presence of concomitant dosing with F68 (initial plasma concentration, calculated by estimating plasma volume at 10% of body weight) at 0.025%. Each group had an n = 3. Blood (<10 μl) was collected using a heparinized syringe at 1, 30, 120, 600 minutes post-injection and plasma prepared. Both blood and plasma samples were snap frozen on dry ice and stored at −80°C before analysis. After the last blood sample, the mice were perfused with sterile 0.9% physiological saline (via the hepatic portal vein), the brains removed, weighed and snap frozen. Analysis of samples was by a validated weak cation exchange solid phase extraction (WCX-SPE) approach performed by a specialist contract research organization (Cyprotex). Briefly samples were diluted with water, WCX-SPE sorbent was primed with MeOH and then water (to ensure phase was fully ionised). Samples were then loaded onto sorbent and washed with pH7 buffer and MeOH. Pentamidine was then washed off sorbent by eluting with a combination of MeOH/H_2_O + 5% v/v formic acid. If necessary samples were then evaporated to dryness and reconstituted in injection solvent. Samples were analysed by UPLC-MS/MS as described above. LLOQ in plasma samples was 2 ng/ml and in brain samples was 80 ng/ml.

Additional experiments revealed that intravenous administration of 10mg/kg pentamidine isethionate plus or minus 0.5% F68 was toxic to the mice and the experiment was terminated.

##### Data analysis

All data are presented as means ±S.E.M and statistical analysis was carried out using Sigma Stat software, version 12.0 (SPSS Science Software UK Ltd, Birmingham, UK).

### 2.9 Trypanocidal activity *in vitro*

*In vitro* activity of drug formulations against *Trypanosoma brucei* blood stream form trypomastigotes was determined *in vitro* using Alamar Blue (resazurin: Bio-Source, Camarillo, CA) as described by [52]. Prior to determination of the trypanocidal activity of Pluronic-pentamidine combinations, the IC_50_ values of the Pluronic alone was established. Each Pluronic was tested in a 3-fold serial dilution in triplicate and in three separate experiments (n=3). The diluent was HMI-9 media (Invitrogen, UK). Blood stream form *T. b. brucei* (strain S427) trypomastigotes, cultured in modified HMI-9 media supplemented with 10% v/v heat-inactivated foetal calf serum, (hi-FCS, Gibco, Life Technologies, UK), were incubated (37°C; 5% CO_2_) at a density of 2 × 10^4^/ml in the presence of pentamidine alone or pentamidine-Pluronic formulations for 66h. Resazurin (20 µl 0.49mM in PBS) solution was then added to each well and incubation continued for 6 hours. After incubation, samples were removed and fluorescence was measured using excitation 530nm and emission 590nm on a Spectramax M3 plate reader (Molecular Devices, USA). IC_50_ values were determined (where appropriate) using GraphPad Prism.

## 3. Results

### 3.1 Evaluation of potential neurotoxicity of pentamidine

#### 3.1a Literature review

We conducted a brief review of the literature to assess the potential neurotoxicity of pentamidine. Information was considered relevant to the NanoHAT project if it described an activity that could be detected in a simple profiling screen, rather than secondary readouts (e.g. hERG-mediated, downstream effects on cardiomyocyte [Ca^2+^]i). Table 2 lists the known pharmacology and approximate affinities of the interaction that have been reported for this compound.

**Table 2.**
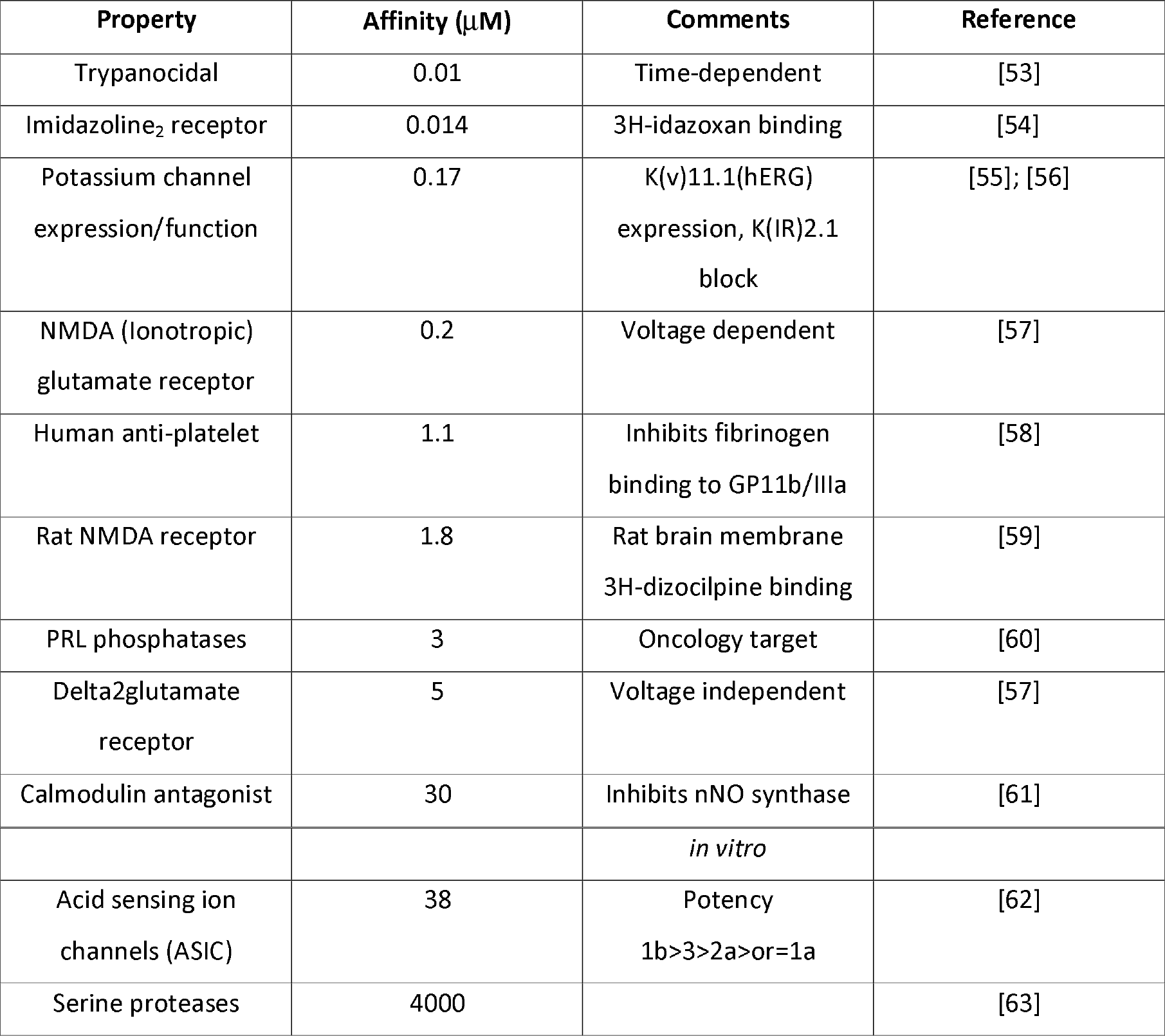
Reported Pharmacology of Pentamidine *in vitro*.

As the trypanocidal activity of pentamidine occurs at around 10 nM *in vitro*[53], we considered that any affinity greater than 1 µM (i.e. more than 100-fold greater than the trypanocidal concentration) was unlikely to be relevant.

There are 3 major target families for which pentamidine has significant affinity (<20 fold above trypanocidal range) that were of concern: the imidazoline_2_ receptor (responsible for effects on central blood pressure control and pancreatic beta cells); inward rectifying (IR) potassium channels particularly blockade of Kir2.1 (this is more likely cardiac than CNS-relevant) and NMDA glutamate receptors.

#### 3.1b A neurological profiling screen

A wide ligand profiling screen was carried out against 40 CNS targets (Perkin Elmer customised CNS screening; listed in Table S1), testing at a single concentration of 10 µM (1000-times the trypanocidal concentration), with follow up concentration-response curves in any assay where there was greater than 70% inhibition. Pentamidine was inactive at 29 out of 40 CNS targets (including 5 glutamate receptor binding sites) at 10 µM, and was re-tested against the remaining targets at a range of concentrations to generate an inhibitory constant, K_i_. These results, together with the calculated relative selectivity values compared with trypanocidal affinity, are listed in Table 3

**Table 3.** K_i_ Values for Pentamidine Determined for Selected CNS targets.

| Target | $K_i$ (μM) | Relative to trypanocidal activity |
| --- | --- | --- |
| Trypanocidal Activity | 0.01 | 1.0 |
| Imidazoline 1<sub>2</sub> | 0.001 | 0.1 |
| Monoamine oxidase B | 0.181 | 18 |
| Monoamine oxidase A | 0.217 | 22 |
| Adrenergic alpha1 | 0.273 | 27 |
| Muscarinic (central) | 0.281 | 28 |
| Histamine H<sub>2</sub> | 7.21 | 721 |
| Opioid | 1.41 | 141 |
| DA transporter | 2.11 | 211 |
| Adrenergicalpha2 | 10 | 1000 Estimate from single-point screen |
| Adrenergic $\beta$ | 10 | 1000 Estimate from single-point screen |
| 5HT transporter | 10 | 1000 Estimate from single-point screen |

Selectivity screening of pentamidine identified 5 targets (imidazoline I_2_ receptor; monoamine oxidase A and B; adrenergic α_1_ receptor; muscarinic receptor) for which it has significant affinity, and which should be monitored as we progressed through the screening cascade. In particular, pentamidine’s high affinity for the imidazoline receptor may explain the cardiovascular adverse events associated with this drug. The project team considered that remaining targets were of minor concern, as the adverse events of drugs targeting the adrenergic monoamine oxidase and muscarinic systems are reasonably well described. The relatively low affinity of pentamidine for the remaining targets (histamine H_2_ receptor; opioid receptor; adrenergic α_2_, β receptors; 5HT transporter) indicated that the drug was unlikely to have significant effects until plasma/brain levels exceeded ~ 100-fold the trypanocidal concentration.

#### 3.1c Ion channel screen

We carried out ion channel screening at Chantest in order to investigate the potential potassium (K(IR)2.1) blocking liability reported by de Boer et al., (2010) (Table 2). Pentamidine isethionate salt was evaluated at 0.001, 0.01, 0.1, 1 and 10 µM (Table S2). The IC_50_ value for pentamidine isethionate salt could not be calculated as the highest tested concentration resulted in hKir2.1 inhibition less than 50% (i.e. 12.3±1.3%). The IC_50_ is estimated to be greater than 10 µM. The positive control (100 μM barium) confirms the sensitivity of the test system to ion channel inhibition.

### 3.2 Formulation Development

As this was a milestone driven project an iterative, dynamic approach was utilized to select the lead formulation to take forward as quickly as possible in the screening cascade (Fig 1), hence not all Pluronic formulations were assessed with each of the methods.

#### 3.2a. Phase Behaviour

L61 phase diagrams were evaluated by visual inspection from 20°C to 50°C for L61 alone and in mixtures with P105 and/or F68 in water and saline solutions. L61 presents a cloud point around 24°C [64]and F68 does not improve its solubility, while P105 does to some extent (Tables S3 and S4).

#### 3.2b. Critical micelle concentration (CMC) by fluorescence spectroscopy

CMC were measured for individual Pluronic and mixtures of F68, P85, P105 and L61 at 20°C and 37°C, both in aqueous and saline (0.9 wt%) solutions, using the intensity of pyrene fluorescence emissions (Table 4; Fig S2). Mixtures of two Pluronics in both aqueous (aq) and saline (sal) mediums were prepared in either a fixed mass ratio of 1:1 or with the addition of 0.01% w/v L61 and the CMC determined. All CMC curves show two inflection points, a feature widely reported in the literature; the first corresponds to the onset of aggregation and was chosen as the CMC (Fig S2; Table 4), giving the following values in saline solution at 37°C (g/L): P85_sal_=0.042±0.018; F68_sal_=0.048±0.012 and P105_sal_=0.069±0.020. Overall, these CMC values are fairly similar and do not allow a prioritisation based on CMC alone. The CMC of F68 and P85 mixtures (1:1 mass ratio) is about double the CMC, when expressed in total Pluronic mass, of the individual polymers suggesting the absence of mixed micelles in these mixtures. Small amounts of L61 (0.01%w/v) does not affect the CMC of F68 or P85 or P105 under the conditions tested

#### 3.2c Stability of the formulations

Pentamidine stability in solution was followed by NMR. Pentamidine and pentamidine/Pluronic solutions prepared in D_2_O were kept in amber NMR tubes at 37°C. Spectra were measured at days 0, 1 and 7. As a control, pentamidine in D_2_O was left at 4°C and measured at day 0 and 7. NMR data showed no significant change on peak position or peak intensity when compared to day 0 measurements or to control samples, confirming no thermal degradation of pentamidine after 7 days at 37°C

#### 3.3d Partition

Partition of PTI in the micelles was measured by fluorescence spectroscopy for P105 and F68. Pentamidine has a slightly larger partition coefficient in F68 than in P105 (Table 5). Measurements in mixtures (F68/L61, P105/L61 and F68/P105, 1:1 mass ratio in all cases) do not significantly change the partition coefficient (data not shown).

**Table 4.**
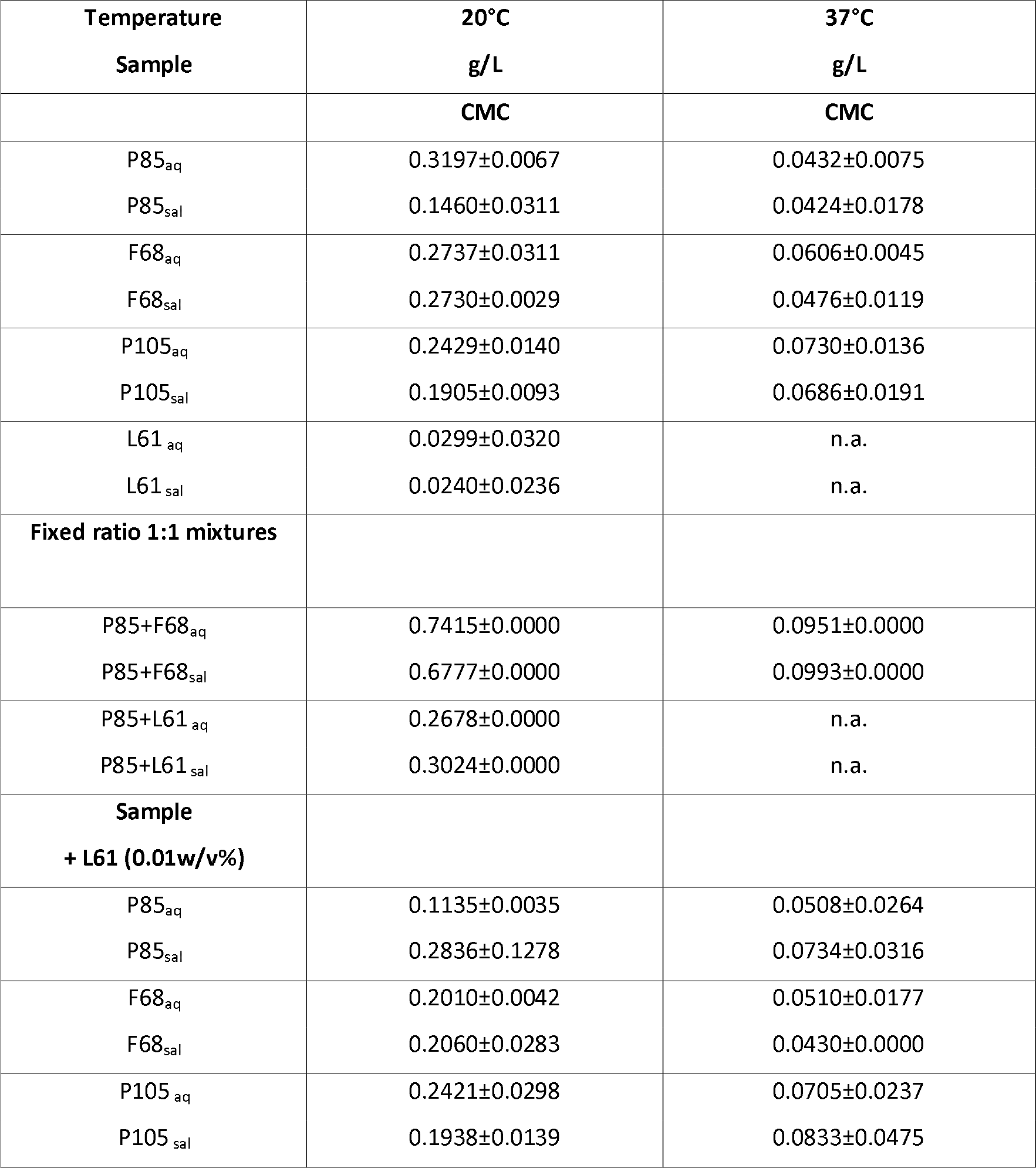
CMC Values of Pluronic Dissolved in Pure Water (aq) or Saline (sal) at 20°C and 37°C Determined Using Pyrene Fluorescence Intensity. Values Mean ± S.D. Saline (0.9 wt%).

| Temperature<br>Sample | 20°C<br>g/L | 37°C<br>g/L |
| --- | --- | --- |
|  | CMC | CMC |
| P85<sub>aq</sub> | 0.3197±0.0067 | 0.0432±0.0075 |
| P85<sub>sal</sub> | 0.1460±0.0311 | 0.0424±0.0178 |
| F68<sub>aq</sub> | 0.2737±0.0311 | 0.0606±0.0045 |
| F68<sub>sal</sub> | 0.2730±0.0029 | 0.0476±0.0119 |
| P105<sub>aq</sub> | 0.2429±0.0140 | 0.0730±0.0136 |
| P105<sub>sal</sub> | 0.1905±0.0093 | 0.0686±0.0191 |
| L61<sub>aq</sub> | 0.0299±0.0320 | n.a. |
| L61<sub>sal</sub> | 0.0240±0.0236 | n.a. |
| **Fixed ratio 1:1 mixtures** |  |  |
| P85+F68<sub>aq</sub> | 0.7415±0.0000 | 0.0951±0.0000 |
| P85+F68<sub>sal</sub> | 0.6777±0.0000 | 0.0993±0.0000 |
| P85+L61<sub>aq</sub> | 0.2678±0.0000 | n.a. |
| P85+L61<sub>sal</sub> | 0.3024±0.0000 | n.a. |
| **Sample**<br>**+ L61 (0.01w/v%)** |  |  |
| P85<sub>aq</sub> | 0.1135±0.0035 | 0.0508±0.0264 |
| P85<sub>sal</sub> | 0.2836±0.1278 | 0.0734±0.0316 |
| F68<sub>aq</sub> | 0.2010±0.0042 | 0.0510±0.0177 |
| F68<sub>sal</sub> | 0.2060±0.0283 | 0.0430±0.0000 |
| P105<sub>aq</sub> | 0.2421±0.0298 | 0.0705±0.0237 |
| P105<sub>sal</sub> | 0.1938±0.0139 | 0.0833±0.0475 |

**Table 5.** The fraction of pentamidine incorporated into the Pluronic micelle expressed as a partitioning coefficient, P. The Pluronic was dissolved in pure water (aqueous) or saline (saline) at 20°C and 37°C. (Also see Fig S3).

| Pluronic | Solvent | Temperature<br>°C | Log P |
| --- | --- | --- | --- |
| P105 | saline | 20 | 1.06 |
|  |  | 37 | 1.15 |
| P105 | water | 20 | 0.99 |
|  |  | 37 | 1.09 |
| F68 | saline | 20 | 1.67 |
|  |  | 37 | 1.47 |
| F68 | water | 20 | 1.67 |
|  |  | 37 | 1.46 |

The values of Log P obtained in saline and aqueous solutions are rather similar, suggesting that pentamidine partition is not sensitive to the saline levels used here.

The effect of temperature is quite weak (Table 5), and does not follow the same trend with the two Pluronic studied: values of LogP for P105 are lower at 20°C than at 37°C (but still very close); instead, for F68 the partition of PTI decreases slightly at higher temperature.

At biologically relevant concentrations, 0.5 wt% Pluronics and 1.0×10^−6^ M PTI, extrapolation of the Log P data suggest that ca. 0.1 PTI molecules would be incorporated in one P105 micelle, and 0.01 PTI molecules in one F68 micelle. At the concentrations used for SANS (5 wt% Pluronic and 1 wt% PTI), extrapolating these numbers give 166 PTI molecules in the micellar core per P105 micelle and 15 for F68 micelle.

The relative low values of log P for PTI/Pluronic system (for comparison log P for pyrene/Pluronics is ca. 2.5 and 3.5 for F68 and P105, respectively [11]), is not surprising given the high water solubility of pentamidine, and helps to explain the drug release profile for PTI / Pluronics systems discussed next. Overall, this means that Pluronic have a limited capacity to interact with pentamidine and prolong its circulation.

#### e. Drug release

Solutions of 10 mM pentamidine or 10 mM pentamidine plus 1 % F68 or 1% P105 were loaded in dialysis cells and the amount of pentamidine eluting from the cells into water at 37°C were measured over time (Fig S4). Both reaction type and reaction constant, for PTI alone and PTI/Pluronic were in a similar range. ca. 0.5 (Fickian diffusion) for reaction type and ca 0.3 for reaction constant. No significant difference was observed between PTI/Pluronics and PTI/water systems. Thus in the conditions tested, pentamidine release seems to be dominated by diffusion and Pluronic micelles were not a barrier for drug release.

#### f. Aggregation number and Micellar size

Pluronic micelles can be reasonably described as a compact core formed by a dry PPO block surrounded by a highly hydrated shell formed by the two PEO blocks[65][66]. The core-shell model was thus used to provide a more detailed characterisation of the morphology of the Pluronic micelles in D_2_O and how it is affected by the presence of PTI, using input values for the core radius and shell thickness were based on hydrodynamic radius values obtained by DLS (Table 6). A term to compensate for polydispersity was included for both Pluronics, as well as a structure factor (S(q)), corresponding to a hard sphere model, in order to account for intermicellar interactions. A summary of the main parameters obtained from the analysis of the data (Fig S5) is present in Table 7.

**Table 6.**
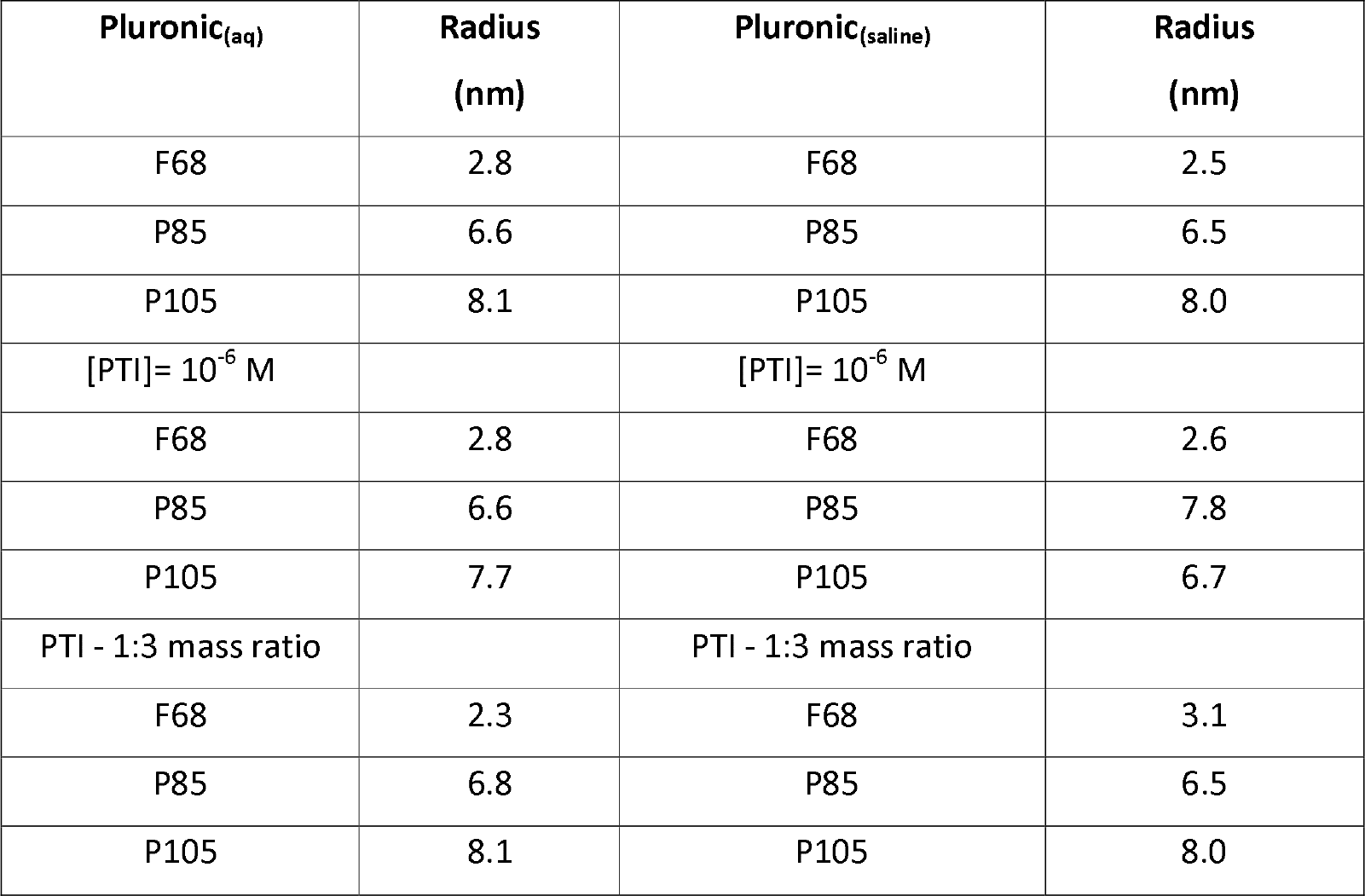
Stokes Radii of P105, P85 and F68 Micelles Obtained from DLS (1% w/w, 37°C).

| Pluronic<sub>(aq)</sub> | Radius<br>(nm) | Pluronic<sub>(saline)</sub> | Radius<br>(nm) |
| --- | --- | --- | --- |
| F68 | 2.8 | F68 | 2.5 |
| P85 | 6.6 | P85 | 6.5 |
| P105 | 8.1 | P105 | 8.0 |
| [PTI]= 10<sup>-6</sup> M |  | [PTI]= 10<sup>-6</sup> M |  |
| F68 | 2.8 | F68 | 2.6 |
| P85 | 6.6 | P85 | 7.8 |
| P105 | 7.7 | P105 | 6.7 |
| PTI - 1:3 mass ratio |  | PTI - 1:3 mass ratio |  |
| F68 | 2.3 | F68 | 3.1 |
| P85 | 6.8 | P85 | 6.5 |
| P105 | 8.1 | P105 | 8.0 |

**Table 7.** Geometric parameters from model-fitting of the SANS Pluronic data at 37°C, including core and shell micellar sizes, fraction of solvent in the corona (χ_solv_) and aggregation number (N_agg_). (Also see Fig S5).

| Sample | Core radius<br>(Å) | Shell<br>thickness<br>(Å) | Total radius<br>(Å) | $\chi_{solv}$ | $N_{agg}$ |
| --- | --- | --- | --- | --- | --- |
| F68 5% | 15.4 | 36.5 | 52.0 | 0.99 | 2.37 |
| F68 5%/ PTI 1% | 15.1 | 34.7 | 49.8 | 0.98 | 2.25 |
| F68 5%/ PTI 3% | 15.5 | 33.9 | 49.4 | 0.99 | 2.38 |
| P85 5% | 42.9 | 31.4 | 74.3 | 0.95 | 35.4 |
| P85 5%/ PTI 1% | 41.5 | 30.5 | 72.0 | 0.95 | 32.4 |
| P85 5%/ PTI 3% | 41.0 | 30.5 | 71.5 | 0.99 | 31.6 |

A direct comparison of F68 and P85 micelles in D_2_O shows that both have similar shell thickness, with F68 showing values slightly larger, 36.5 Å vs 31.4 Å, respectively. It is worth noting that F68 EO blocks have on average 76.4 EO units while P85 blocks are only 26.1 units long. The core of F68 micelles are significantly smaller than P85 micelles, 15.4 vs 42.9 Å. In terms of PO content, the F68 PO block is 29 units long while P85 is 40 units long. Overall, P85 micelles are larger than F68 micelles, 74.3 vs 52.0 Å, respectively.

The simulation work agrees well with these experimental results. The average aggregation number per micelle (N_agg_) and the average number of micelles (N_mic_) were calculated once the systems had equilibrated have been measured. Fig 2 shows plots of N_agg_ and N_mic_ as a function of Pluronic concentration for both the F68 and P105 Pluronics. We carried out simulations over a range of Pluronic concentrations that span the CAC and the CMC values observed experimentally to validate the models (at least qualitatively). From Fig 2, one can see that in both systems, once we have passed the CAC the number of micelles remains more or less constant but they continue to grow in size as the concentration increases until we reach the CMC at which point the size of the micelles more or less plateaus. Also, when comparing the behavior of the F68 and P105 Pluronics, we found that the P105 Pluronics form larger aggregates when near the CMC as compared to that for the F68 Pluronics, and therefore fewer micelles. Note, we have also simulated mixtures of F68 and L61 Pluronics, and the results of those systems are presented in Fig S6.

**Fig 2.**
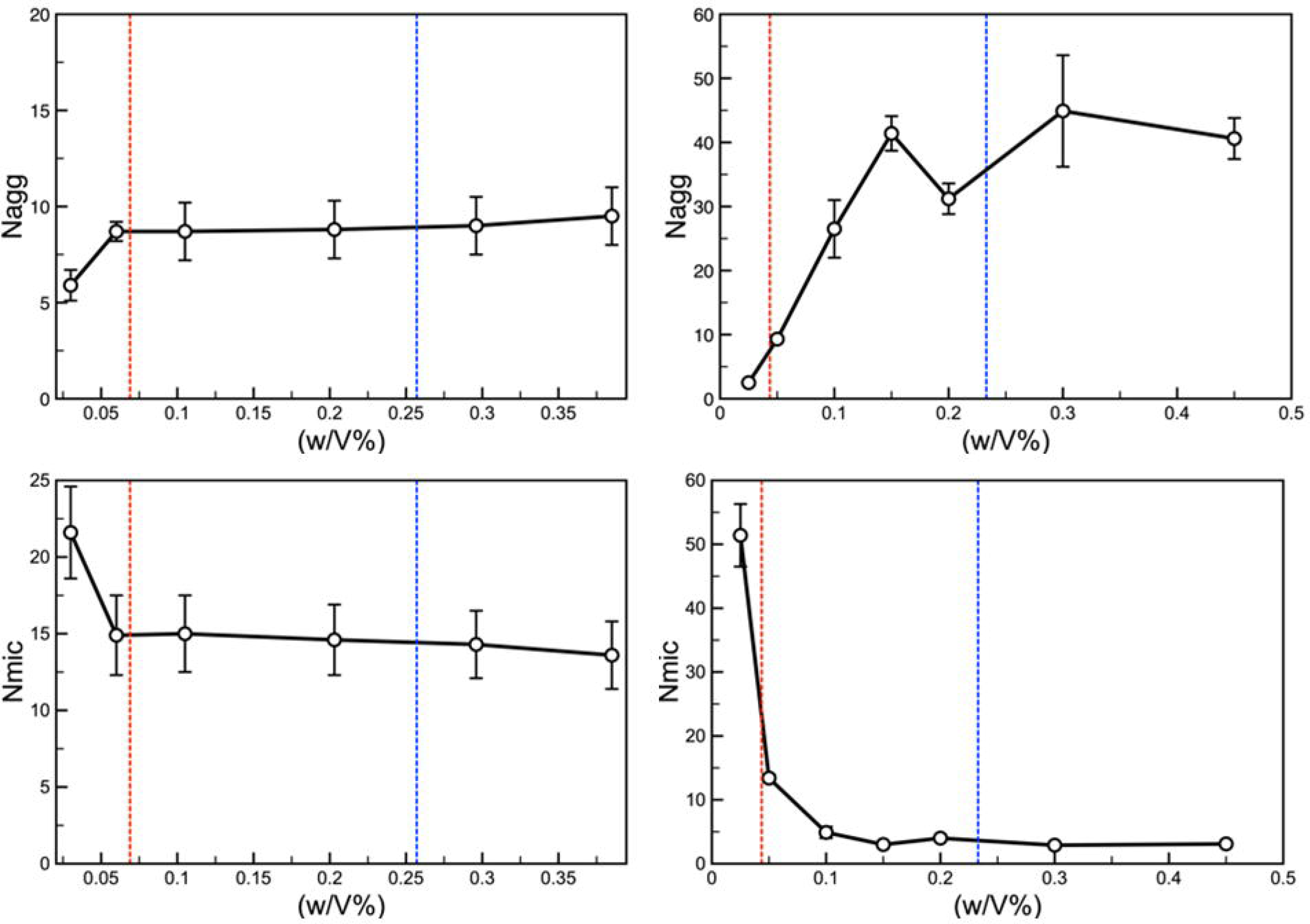
The average number of Pluronic molecules found in a micelle (N_agg_) and the number of micelles in our system (after they have equilibrated) (N_mic_) as a function of the concentration of the Pluronics in the system for both the F68 (left) and P105 (right) Pluronics. Additionally, we have compared the findings from the simulations with the identified values (dashed lines) of the CAC and CMC from the experimental systems.

In the presence of 1% PTI, a small reduction in size was observed for both Pluronics, ca. 2 Å in both cases. The increase to 3% PTI does not cause further changes.

The coronas were highly hydrated, as reported for these polymers [67][68]. F68 micelles were more hydrated than P85: for each EO unit in the shell, there were are 17 D_2_O molecules in a F68 micelle but only 3.4 in a P85 micelle.

The addition of pentamidine leads to a subtle, but perceptible, reduction of the number of water molecules in the F68 micelle shell. For P85, no measurable changes were observed.

### 3.3 Peripheral Toxicity

Pluronic concentrations used in the biological assays were based on the CMC measurements. Peripheral toxicity of the individual polymers was assessed. L61 was not studied at this stage due to its limited solubility.

#### a. Haemolysis assay

All the Pluronics (P85, F68 and P105) at each of the tested concentrations (0.01%, 0.1%, 0.25%, 1% or 5%) and both the negative controls (0.05% DMSO or 0.9% saline) did not cause any haemolysis (0%) of the human cells. In contrast the positive control, 10% Triton X-100, caused haemolysis (***p<0.001 compared to the negative controls).

#### b. Permeability assay (MDCK-MDR) to assess membrane integrity

[^14^C(U)]sucrose is an inert, polar molecule which normally does not cross cell membranes, but may cross between cells through the paracellular cleft. Significant differences in [^14^C(U)]sucrose Papp values existed in the presence of all tested concentrations of P85, and 0.5% and 0.01% P105 compared to [^14^C(U)]sucrose P_app_ in the absence of Pluronic (Fig S7) indicating loss of monolayer integrity. No tested concentration of F68 significantly affected the radiolabelled sucrose P_app_ values.

#### c. Effect of Pluronic on insulin secretion and beta-cell viability

Exposure of MIN6 β-cells to 1 and 100μM pentamidine for 24 hours caused a concentration-dependent inhibition of acute insulin secretion in response to the cyclic AMP elevating agents forskolin and IBMX (Fig 3). Surprisingly, P85 and 105 were significantly more effective than pentamidine in inhibiting insulin secretion in response to forskolin/IBMX, such that insulin release was fully inhibited by these Pluronics in the absence of pentamidine at all concentrations tested (0.01-0.5% w/v) (Fig 3A-D). Low concentrations of F68 (0.01 and 0.025% w/v) generated similar inhibitory effects on insulin secretion as unformulated pentamidine (Fig 3A-B) and increased toxicity was observed with higher concentrations of F68 (Fig 3C-D).

**Fig 3.**
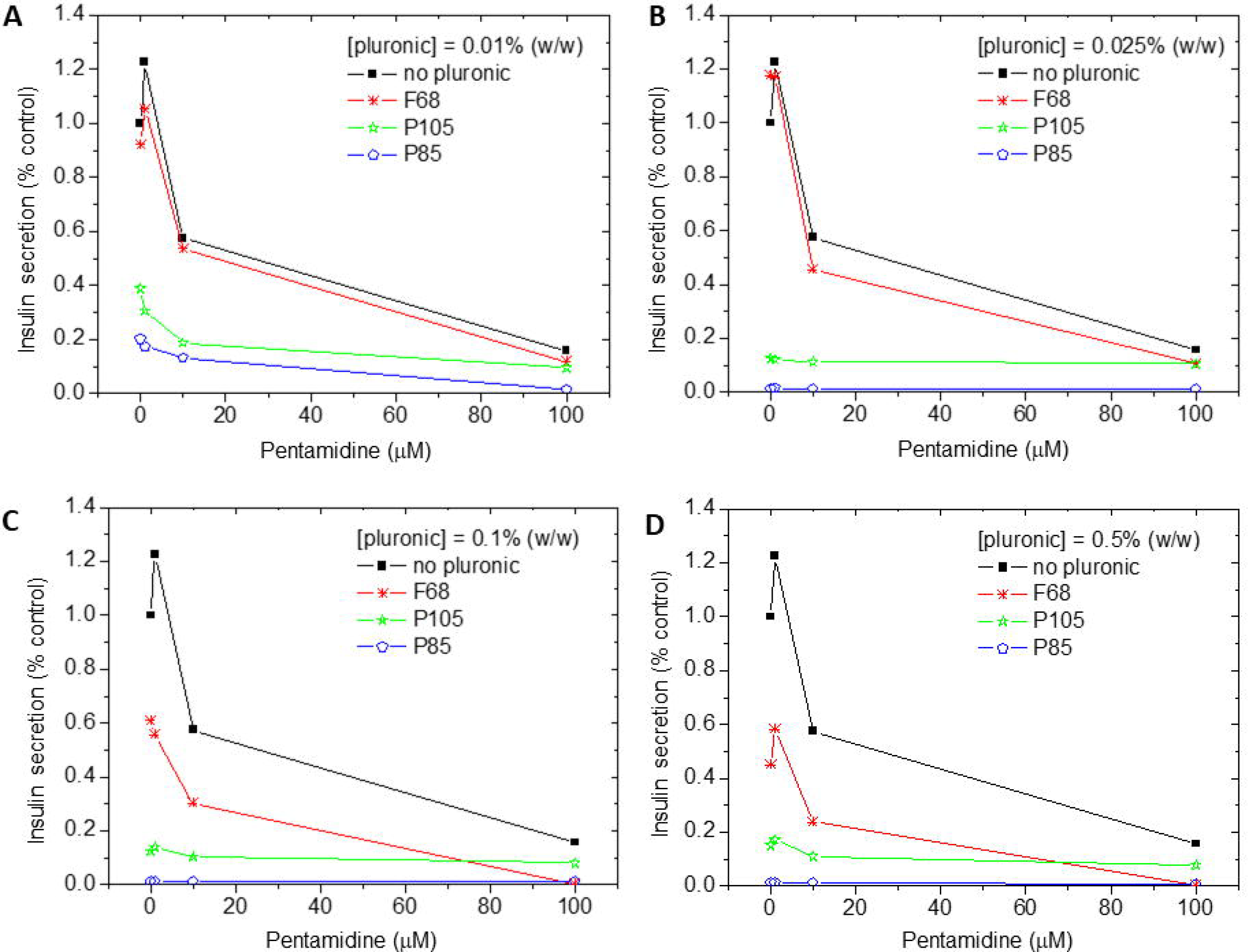
The effect of Pluronic and pentamidine in DMEM on insulin secretion from MIN6 β-cells expressed as a percentage of control. Control values were obtained from DMEM supplemented with 2mM glucose. (A-D) P85 and P105 induced a strong suppression of insulin secretion from MIN6 β-cells even at low concentrations. (C-D) F68 only induced insulin secretion suppression at concentrations ≥0.1% w/v.

Trypan blue staining indicated that the MIN6 β-cells were able to tolerate pentamidine concentrations of 1 and 10 μM, but 100 μM pentamidine, which induced maximal inhibition of insulin secretion, was accompanied by a large number of cells taking up Trypan blue (Fig S8 and S9). These micrographs are indicative of the suppression of insulin secretion by pentamidine being associated with marked reductions in β-cell viability, but the plasma membrane was largely intact as there was no leakage of insulin, a 5.5 kDa peptide, from the cell interior. The combination of 100 μM pentamidine with 0.5% w/v F68, which caused maximal suppression of insulin release (Fig 3), led to the highest proportion of cells that showed Trypan blue staining.

### 3.4 Trypanocidal activity *In vitro*

The *in vitro* activity of Pluronic drug formulations alone against *T. b. brucei* blood stream form trypomastigotes was determined showing low trypanocidal activity of F68 compared to high activity of P85 and P105 (Table 8).

**Table 8.**
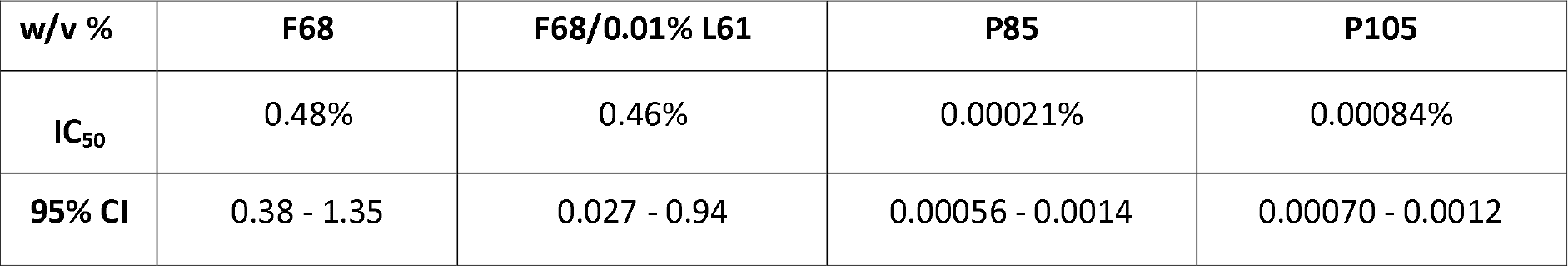
The Inhibitory Concentration (IC_50_) required to reduce number of bsf trypomastigotes by 50%. Pluronic were tested at 12 serial dilutions in triplicate and repeated in 3 separate experiments (n=3) to produce IC_50_ values.

In further studies the anti-trypanosomal activity of combinations of F68 and pentamidine were assessed (Table 9). The IC_50_ (± 95% CI) values of pentamidine were 2.11 × 10^−5^ ± (1.79 × 10^−5^ – 2.50 × 10^−5^) μM alone, 6.36 × 10^−6^ (± 4.43 × 10^−6^ – 9.12 × 10^−6^) μM with 0.01% F68 and 3.25 × 10^−6^ ± (3.13 × 10 ^−7^ – 3.38 × 10^−5^) μM with 0.001% F68.

**Table 9.** The % of bsf trypomastigotes inhibited by pentamidine/pluronic combinations. The combination formulation was tested in triplicate and repeated in 3 separate experiments (n=3).

|  | Pentamidine (µM) |  |  |  |  |  |
| --- | --- | --- | --- | --- | --- | --- |
| | 1 | 0.3 | 0.000152 | $5.1 \times 10^{-5}$ | $1.7 \times 10^{-5}$ | $5.7 \times 10^{-6}$ |
| F68 (w/v %) |  |  |  |  |  |  |
| 0.5% | 99.5% | 98.6% | 98.6% | 98.3% | 98.3% | 99.2% |
| 0.1% | 98.5% | 97.7% | 97.1% | 97.1% | 97.3% | 97.7% |
| 0.025% | 98.3% | 97.5% | 97.0% | 96.9% | 97.0% | 90.6% |
| 0.01% | 98.4% | 97.6% | 96.4% | 95.1% | 82.8% | 3.4% |
| 0.001% | 98.3% | 97.4% | 96.4% | 91.9% | 73.1% | 1.8% |
| 0% | 98.3% | 97.4% | 92.7% | 65.3% | 35.0% | 4.1% |

To determine if the addition of Pluronic to pentamidine had an additive effect on the trypanocidal activity of pentamidine, it was decided that work should focus on F68 rather than the other Pluronics, as both P85 and P105 caused an inhibitory effect on insulin secretion. Although IC_50_ values could only be determined for two combinations, in part due to the high starting concentration of pentamidine used, a limited interaction between Pluronic F68 and pentamidine was observed at the lowest F68 concentrations (Table 9 boxes shaded in red), suggesting that the addition of Pluronic had an additive effect on the trypanocidal activity.

### 3.5 Blood-brain barrier: *In vitro* permeability assays

We examined the ability of different pentamidine-Pluronic formulations to cross the BBB using the MDR1-MDCK cell line. Two analytical methods were applied: one detected pentamidine isethionate using UPLC-MS/MS (Table S5) and the other detected radiolabelled pentamidine using liquid scintillation counting (Table 10).

**Table 10.** The Effect of P85, F68 and P105 on the Apparent Permeability of [^3^H(G)]pentamidine (9 nM) MDR1-MDCK Cell Monolayers in the Apical to Basolateral Direction and the Basolateral to Apical Direction. The percentage recovery of pentamidine is also shown. All the data has been corrected for extracellular space by subtracting [^14^C(U)]sucrose (5.5 μM) P_app_ values which ranged from 0.89 to 2.00 × 10^−6^ cm/s. Each value represents three replicates for each n and n =3. n.d. = not determined as integrity of the barrier compromised.

| [<sup>3</sup>H(G)]Pentamidine<br>(9 nM) | Pluronic Concentration<br>(%) | P<sub>app</sub> A2B<br>(10<sup>-6</sup> cm/s) | P<sub>app</sub> B2A<br>(10<sup>-6</sup> cm/s) | A2B<br>(%) | B2A<br>(%) |
| --- | --- | --- | --- | --- | --- |
|  |  | Mean±SEM | Mean±SEM | Mass<br>balance | Mass<br>balance |
|  | 0 | 0.678±0.025 | 0.776±0.062 | 84 | 85 |
|  | 0.01% P85 | 0.310±0.142 | 0.431±0.161 | 86 | 87 |
|  | 0.1% P85 | 0.561±0.0172 | 0.227±0.081 | 89 | 89 |
|  | 0.5% P85 | n.d. | n.d. | 90 | 90 |
|  | 0.01% P105 | 0.577±0.0710 | 0.818±0.086 | 86 | 89 |
|  | 0.1% P105 | 0.898±0.161 | 0.776±0.054 | 89 | 88 |
|  | 0.5% P105 | n.d. | n.d. | 91 | 91 |
|  | 0.01% F68 | 0.200±0.115 | 0.106±0.061 | 95 | 83 |
|  | 0.1% F68 | 0.221±0.067 | 0.033±0.019 | 98 | 87 |
|  | 0.5% F68 | 0 | 0 | 98 | 84 |

The ability of pentamidine isethionate to cross the MDR1-MDCK cell monolayers in the apical (luminal) to basolateral (abluminal) direction was limited as the concentration of pentamidine isethionate in the basolateral chamber were below the limits of UPLC-MS/MS detection, even when the pentamidine isethionate concentration was increased to 20 μM (data not shown). The lower limit of quantification (LLQ) for UPLC-MS/MS method was ≤ 7.8 nM which is equivalent to <0.039% of the dose in the donor (apical) chamber. These results would suggest that pentamidine is a ‘CNS negative’ drug. The mass balance (% recovery) results suggest that up to 25% may have been trapped either in the endothelial cells or by non-specific binding to the plastic plate or the polycarbonate membrane and this may contribute to the reduced recovery. It is important to note that the two amine groups of pentamidine are ionized at physiological pH (pKa is 11.4) and are likely to react with static charges on the surface of plastics. Interestingly the presence of Pluronic increased the mass balance results by 10-20%. In contrast the movement of pentamidine isethionate (20μM) across the monolayer in the basal to apical direction was measurable, the calculated P_app_ being 0.418 × 10^−6^ cms^−1^. Taken together with the absence of a detectable movement of pentamidine isethionate in the apical to basolateral direction these data would indicate the presence of an efflux mechanism for this molecule, likely MDR1. The presence of Pluronics (F68, P105 or P85 at concentrations of 0.01%, 0.1% or 0.5%) did not affect the distribution of pentamidine isethionate across the monolayer in either direction.

MDR1-MDCK permeability assay experiments were also performed using radiolabelled pentamidine and liquid scintillation counting. The rate of transport of [^3^H(G)]pentamidine (9 nM) across MDR1-MDCK monolayer was examined and in contrast to the results achieved with the pentamidine isethionate was measurable in both directions (Table S5). In these experiments the highest concentrations of P85 (0.5%) and P105 (0.5%) affected the integrity of the cell monolayer and the Papp for [^14^C(U)]sucrose was increased to 4.8 and 4.3 × 10^−6^ cm/s respectively. The [^14^C(U)]sucrose P_app_ for all other experiments was 1.61±0.15 × 10^−6^ cm/s. The presence of the Pluronics (F68, P105 or P85) at concentrations of 0.01% and 0.1% did not significantly increase the distribution of [^3^H(G)]pentamidine across the MDR1-MDCK monolayer measured over 60 minutes confirming the results obtained using pentamidine isethionate and the UPLC-MS/MS detection method.

In conclusion, our target formulation characteristics of at least a 2-fold increase in pentamidine / pentamidine isethionate movement across the monolayer, compared with unformulated pentamidine, was not observed using these *in vitro* models of BBB permeability.

### 3.6: Blood-brain barrier In situ brain perfusion

#### 3.6a PLURONIC P85

Co-formulation of 15.7 nM [^3^H(G)]pentamidine with Pluronic P85 did not significantly increase [^3^H(G)]pentamidine accumulation in any of the brain regions examined (Table S6).

In fact, overall there was a decrease in the [^14^C(U)]sucrose-corrected uptake of [^3^H(G)]pentamidine with P85 at 0.01% (p<0.001) and at 0.1% (p<0.01) (Two-Way ANOVA), but these decreases were not statistically significant when individual brain regions were examined using Bonferroni’s pairwise comparisons.

Table S6 shows the results of capillary depletion analysis of the brain tissue after 10 minutes of perfusion in the presence or absence of the Pluronic, P85. [^3^H(G)]pentamidine accumulated in the capillary endothelial cells (pellet) of control mice while less than 2% of the plasma concentration crossed the basolateral membrane to reach the parenchyma (supernatant). These results are in good agreement with our previously published data[49]. There appeared to be an overall reduction in this accumulation of [^3^H(G)]pentamidine into the endothelial cells and consequently a reduction in the parenchyma when [^3^H(G)]pentamidine was co-formulated with 0.01% and 0.1% P85, but the reduction did not attain statistical significance (Two-Way ANOVA with Bonferroni’s pairwise comparisons).

Whilst there appeared to be an overall inhibitory effect of Pluronic on the transport of [^3^H(G)]pentamidine across the BBB, there was a significant, 3-fold increase in the uptake of [^3^H(G)]pentamidine into the pituitary gland after 10 minutes of perfusion with 0.1% and 0.5% P85 (p<0.05 at both concentrations; Two-way ANOVA with Bonferroni’s pairwise comparisons). This enhanced uptake of pentamidine appeared to be associated with an approximate 2-fold increase in accumulation of [^14^C(U)]sucrose from 20.5±3.9% ([^14^C(U)]sucrose alone) to 35.0±5.5% (+0.01%P85), 43.0±4.0% (+0.1% P85) and 34.3±7.9% (+0.5% P85). A similar effect was observed in the choroid plexus sampled from a few individual mice that were perfused with pentamidine co-formulated with P85 at concentrations above the CMC. This resulted in a 2-fold increase in the mean uptake of [^3^H(G)]pentamidine which was not statistically significant. P85 did not affect accumulation of [^3^H(G)]pentamidine or [^14^C(U)]sucrose by the pineal gland.

#### 3.6b PLURONIC P105

An overall decrease in the [^14^C(U)]sucrose-corrected uptake of [^3^H(G)]pentamidine into brain parenchyma was observed when 15.7nM [^3^H(G)]pentamidine was co-formulated with 0.1% (p<0.001) and 0.5% (p<0.001) P105, as shown in Table S7, but (like P85) these data did not reach statistical significance in any of the individual regions sampled (Two-Way ANOVA with Bonferroni’s pairwise comparisons).

In contrast, there was a 33% increase in the [^14^C(U)]sucrose-corrected uptake of [^3^H(G)]pentamidine into the endothelial cell pellet when it was co-formulated with 0.1% P105 (p=0.027; Two-way ANOVA with Bonferroni’s pairwise comparisons). This increase was apparent in only 3 out of 6 mice, and was associated with penetration of the brain tissue by the vascular space marker [^14^C(U)]sucrose, perhaps indicating an increase in the permeability of the apical/luminal endothelial cell membrane.

No significant differences were observed in the uptake of either the vascular space marker [^14^C(U)]sucrose or [^3^H(G)]pentamidine into the pineal gland, choroid plexus or pituitary gland after 10 minutes of perfusion with [^3^H(G)]pentamidine co-formulated with P105 as shown in Table S7 (p>0.05 for each Pluronic concentration and circumventricular organ; Two-way ANOVA with Bonferroni’s pairwise comparisons).

#### 3.6c PLURONIC F68

##### 10 minute perfusions

Co-formulation of [^3^H(G)]pentamidine with F68 resulted in an overall decrease in accumulation of [^3^H(G)]pentamidine into brain parenchyma after 10 minutes of perfusion (p=0.002 for 0.1% and p=0.03 for 0.5% respectively; Two-way ANOVA with Bonferroni’s pairwise comparisons) (Table S8). A decrease in vascular space as measured by accumulation of [^14^C(U)]sucrose was also measured when 0.01 or 0.1% F68 (but not 0.5%) was present in the artificial plasma (p=0.042 for 0.01% and p=0.004 for 0.1% respectively; Two-way ANOVA with Bonferroni’s pairwise comparisons) (Table S9).

F68 did appear to increase accumulation of [^3^H(G)]pentamidine into the endothelial cell pellet at concentrations of 0.01% and 0.1%, but these results did not attain significance. This increase in [^3^H(G)]pentamidine, did not appear to be associated with a concomitant increase in uptake of [^14^C(U)]sucrose (p>0.05) and might have been due, at least in part, to a small decrease in the amount of drug crossing the basolateral membrane to enter the brain parenchyma, as indicated by a marginal reduction of [^3^H(G)]pentamidine in the supernatant (Table S8).

Co-formulation of [^3^H(G)]pentamidine with 0.5% F68 resulted in a 2-fold increase in uptake into the pituitary gland after 10 minutes of perfusion (p=0.017; 1-way ANOVA with Bonferroni’s pairwise comparisons). A similar, but not statistically significant increase was observed in uptake of [^14^C(U)]sucrose into this organ over the same time period.

##### 30 minute perfusion

Accumulation of [^14^C(U)]sucrose measured in brain parenchyma, as a percentage of concentration in the artificial plasma (R_TISSUE/PLASMA_%), ranged from 1.3% in the hippocampus to 4.3% in the pons after 30 minutes of perfusion. These values are almost identical to our previously published data for BALB/c male mice (1.6 and 4.5% respectively)[7]. Accumulation of [^3^H(G)]pentamidine, when corrected for vascular space ranged from 6.9% in the hippocampus to 15% and 10.9% in the more highly vascularized regions of the hypothalamus and pons respectively. These values were slightly higher than our previously published data (4.3% for hippocampus, 7.6% for hypothalamus and 8.2% for pons) and might reflect changes in expression of transporters due to differences in environment/diet or selective pressures during breeding.

Formulation of 15 nM [^3^H(G)]pentamidine with 0.01% or 0.1% F68 did not affect [^14^C(U)]sucrose brain space (p=0.139 and 0.460 respectively; 2-way ANOVA with Bonferroni’s post-tests). No significant differences were observed in [^3^H(G)]pentamidine accumulation at these concentrations (p=0.120 and 1.000 respectively; 2-way ANOVA with Bonferroni’s post-tests). Similarly, F68 had no significant effect on [^14^C(U)]sucrose or [^3^H(G)]pentamidine accumulation in the capillary depletion samples after 30 minutes of perfusion (p>0.05 for each concentration tested for each isotope; 2-Way ANOVA) nor in the circumventricular organs (p>0.05 for each concentration tested for each isotope; 2-Way ANOVA).

There was an approximate 2-fold increase in accumulation of both [^3^H(G)]pentamidine and the vascular space marker [^14^C(U)]sucrose in the brain parenchyma of mice that were perfused with formulations containing 0.5% F68, (p=0.003 and p <0. 001 respectively; 2-way ANOVA with Bonferroni’s post-tests), as shown in Table S10A and S10B. Visible signs of damage to the BBB including permeation and staining with Evans blue (MW 961), were also observed in some mice. The results from the capillary depletion analysis after 30 minutes of perfusion would also appear to reflect damage to both the apical and basolateral endothelial cell membranes, with a tendency for increased permeation of [^14^C(U)]sucrose into the brain parenchyma, as demonstrated by a small, though not statistically significant rise in this isotope being detected in the supernatant (Table S10).

Co-formulation of [^3^H(G)]pentamidine and [^14^C(U)]sucrose with 0.5% F68 resulted in an increase into the pituitary gland and the choroid plexus when the perfusion time was extended to 30 minutes, although these results were not statistically significant.

### 3.7 *In vivo* pharmacokinetic experiments with pentamidine isethionate or [^3^H(G)]pentamidine

F68 at the 0.025% does not change the accumulation of pentamidine isethionate in the plasma, brain parenchyma or blood in the mouse up to 10 hours post-dosing (Fig 4). There might be a late-onset increase in brain concentrations in the pentamidine alone group, but as the standard deviations for this group at this time-point are large this is unlikely to be significant.

**Fig 4.**
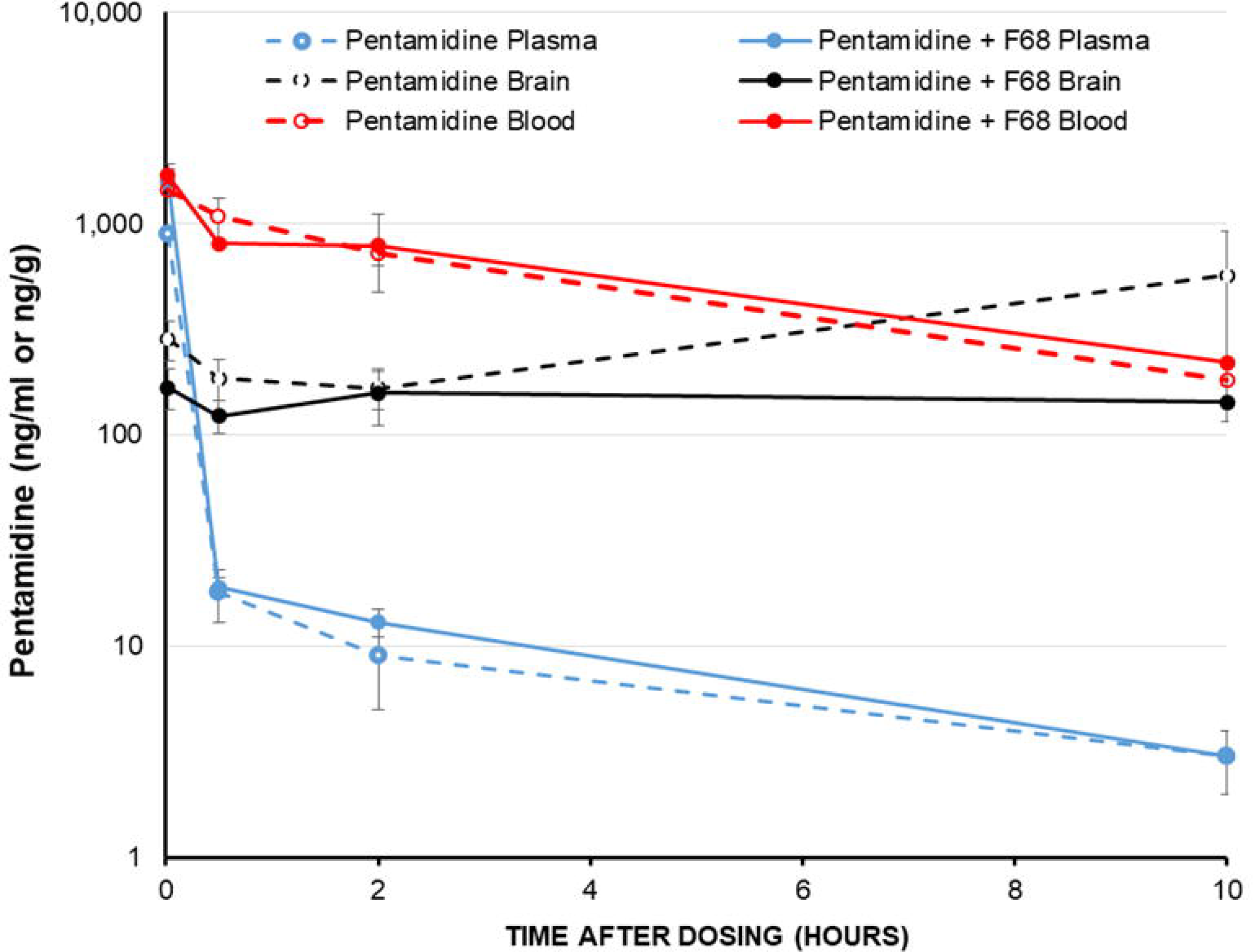
The effect of Pluronic F68 on pentamidine concentrations in mouse plasma, blood and brain after an intravenous dose. Each point represents an n of 3. 4mg/kg pentamidine ± 0.025% F68 i.v. Values ± SD.

Table 11 shows the mean plasma and CSF (corrected for blood/sucrose contamination) concentrations for [^3^H(G)]pentamidine and/or its metabolites, measured at 2 hours after intra-venous injection. No significant differences were observed when [^3^H(G)]pentamidine was co-formulated with either 0.025% or 0.5% F68 (p >0.05 for plasma and CSF; One-way ANOVA). Similarly, no significant differences were observed in uptake of [^3^H(G)]pentamidine or the vascular space marker [^14^C(U)]sucrose, into the brain parenchyma, capillary depletion samples or the circumventricular organs when [^3^H(G)]pentamidine was injected in the presence or absence of F68 (p>0.05; 2-way ANOVA with Bonferroni’s pairwise comparisons) as shown in Table 12.

**Table 11.** Concentration of [^3^H(G)]pentamidine/metabolites in the plasma and CSF at 2 hours post-injection.

| Group | mean concentration in plasma | mean concentration in CSF |
| --- | --- | --- |
| | ng/ml (± SEM) | pg/ml (± SEM) |
| control | 0.343 (± 0.061) | 2.669 (± 0.765) |
| 0.025% F68 | 0.345 (± 0.013) | 1.948 (± 0.826) |
| 0.5% F68 | 0.356 (± 0.026) | 3.592 (± 1.932) |

**Table 12.** Uptake of [^3^H(G)]pentamidine into brain tissue (corrected for vascular space) at 2 hours post-injection. Data is presented as the tissue/plasma ratio (a) and converted into concentrations in ng/g of tissue (b) A limitation of measuring pentamidine by scintillation counting is that any metabolites produced during the 2 hours that have retained the radiolabel, will be counted as [^3^H(G)]pentamidine. These metabolites may have different transport characteristics and may or may not be active against trypanosomes.

| **(a)** | **R<sub>TISSUE/PLASMA</sub>% (mean±SEM)** |  |  |
| --- | --- | --- | --- |
| **Region** | **Control**<br>**(15.7 nM**<br>**pentamidine)**<br>**(n=6)** | **0.025% F68 +**<br>**(15.7 nM**<br>**pentamidine)**<br>**(n=6)** | **0.5% F68 +**<br>**(15.7 nM**<br>**pentamidine)**<br>**(n=5)** |
| **Right brain** | 115.52 (± 12.46) | 120.29 (± 17.14) | 87.36 (± 20.36) |
| **Left brain** | 152.29 (± 33.48) | 111.85 (± 19.15) | 106.10 (± 12.92) |
| **Cerebellum** | 204.02 (± 35.28) | 208.87 (± 28.81) | 172.48 (± 30.34) |
| **Midbrain** | 181.18 (± 45.30) | 254.02 (± 35.48) | 180.00 (± 32.83) |
| **Homogenate** | 249.41 (± 35.59) | 184.18 (± 35.22) | 293.81 (± 122.95) |
| **Supernatant** | 123.35 (± 28.45) | 99.72 (± 9.02) | 98.66 (± 9.47) |
| **Pellet** | 479.72 (± 72.50) | 310.63 (± 38.62) | 536.52 (± 212.72) |
| **Choroid plexus** | 24666.66 (± 4928) | 19628.89 (± 4672) | 20463.70 (± 1827) |
| **Pituitary gland** | 15053.41 (± 3598) | 11285.42 (± 2008) | 15061.87 (± 5321) |

| **(b)** | **Mean concentration**<br>**(ng/g or ng/ml for the supernatant ±SEM)** |  |  |
| --- | --- | --- | --- |
| **Region** | **Control**<br>**(15.7 nM**<br>**pentamidine)**<br>**(n=6)** | **0.025% F68**<br>**(15.7 nM**<br>**pentamidine)**<br>**(n=6)** | **0.5% F68 +**<br>**(15.7 nM**<br>**pentamidine)**<br>**(n=5)** |
| **Right brain** | 0.363 (± 0.035) | 0.417 (± 0.061) | 0.302 (± 0.058) |
| **Left brain** | 0.472 (± 0.084) | 0.383 (± 0.063) | 0.375 (± 0.048) |
| **Cerebellum** | 0.607 (± 0.032) | 0.719 (± 0.097) | 0.591 (± 0.084) |
| **Midbrain** | 0.494 (± 0.075) | 0.866 (± 0.115) | 0.614 (± 0.072) |
| **Homogenate** | 0.820 (± 0.183) | 0.643 (± 0.132) | 0.988 (± 0.375) |
| **Supernatant** | 0.363 (± 0.037) | 0.345 (± 0.035) | 0.351 (± 0.043) |
| **Pellet** | 1.482 (± 0.151) | 1.067 (± 0.125) | 1.827 (± 0.662) |
| **Choroid plexus** | 74.68 (± 11.48) | 84.04 (± 5.78) | 72.20 (± 7.60) |
| **Pituitary gland** | 43.76 (± 3.82) | 37.58 (± 6.54) | 68.13 (± 15.05) |

## Discussion

In this study we generated pentamidine/Pluronic^^®^^ formulations and prioritised 18 formulations using a rational, iterative approach (Fig 1). The milestones were intended to ensure that the most appropriate formulations, on the basis of *in silico* and *in vitro* data, were taken forward to the *in vivo* pharmacokinetic studies and that the formulations with the greatest likelihood of success would be assessed for toxicity issues *in vivo* and tested in animal efficacy models of stage 1 and stage 2 HAT. An ideal formulation for injection should be equipped with characteristics that improved the stability and safety profile of pentamidine, enhanced therapeutic effect, and accelerated the absorbance of drugs. Since increasing the concentration of pentamidine in the brain may cause an intractable neurotoxicity and serious adverse events our starting point was a customised, wide ligand profiling screen carried out against 40 CNS targets (Tables 2 and S1). Five targets (imidazoline I_2_ receptor; monoamine oxidase A and B; adrenergic α_1_ receptor; muscarinic receptor) were identified to have significant affinity for pentamidine (Table 3). All but one of these (imidazoline I_2_ receptor) had a 20-1000 fold lower affinity than the relative trypanocidal activity and did not generate major concern[59]. The activity against the imidazoline I_2_ receptor may explain the cardiovascular adverse events with this drug. We were unable to reproduce the result of De Boer et al., 2010[55] in a recombinant human system indicating that pentamidine was without effect (at up to 10 μM) on the hKir2.1 potassium channel-induced inward rectifying current (Tables 2 and S2). Thus progression could continue through the screening cascade.

For the Pluronics tested in this study (P85, P105, F68 and L61), phase behaviour [69][33] and cloud points [70] are well established. P85, P105 and F68 are soluble in water and saline solutions at both 24°C and 37°C. L61 has a very low cloud point at 24°C. Pure L61 therefore has limitations as a formulation for drug delivery. Our phase diagrams revealed that F68, which is highly hydrated, is unable to improve the solubility of highly hydrophobic L61 to a great extent, so it was not possible to pursue a 1:1 mixture of L61:F68 in the assays (Table S3 and S4).

Using molecular dynamics (MD) simulations and physical techniques, we elucidated the structural properties of Pluronic P85, P105, F68 and L61 micelles, and were able to extract fundamental parameters required for biological evaluation of the formulations. For example, the CMC were measured for F68, P85 and P105 at 20°C and 37°C both in aqueous as well as saline (0.9 wt%) solutions. Several values for the CMC of Pluronics can be found in the literature [71][72][73][11][74]. These values tend to vary widely, showing as much as one order of magnitude differences for the same Pluronic[75]. This has been attributed to several reasons: difference in molecular weight distribution between batches [76][74], presence of impurities such as diblocks[76][77] and differences inherent to the technique employed[78]. In addition, for some Pluronic systems, two critical concentrations are detected, both in surface tension and spectroscopic experiments [76][72]. This behaviour has been ascribed to formation of premicellar aggregates occurring before full micelle formation[79][80][71][72][81]. In this work, which used the intensity of pyrene fluorescence emission, two critical concentrations were also detected (Fig S2). The CMC values presented here (Table 4) are taken from the first break point. The CMC values achieved for F68, P85 and P106 were fairly similar and did not allow a prioritisation of a specific formulation based on CMC alone. The concentrations of Pluronic (0.001 to 0.025%) used in the biological assays were based on the CMC values and were selected on the basis that they would be likely to consist of mainly unimers (0.001-0.025%); a mixture of unimers and micelles (0.1%) and mostly micelles (0.5%) respectively.

F68 micelles have a relatively small radius of 52.0 Å (Table 7). This attribute will increase stability, half-life and therefore circulation time of this Pluronic, since small micelles evade detection and destruction by the reticuloendothelial system. However, this small volume may also correlate to low drug loading (Table 5; Fig S3). In addition, the fact that pentamidine release from both F68 and P105 micelles is by diffusion would indicate that these Pluronics are unlikely to significantly prolong the circulation time of pentamidine (Fig S4).

Haemolysis of human red blood cells was not observed in the presence of 0.5%, 0.1%, 0.025%, 0.01%, and 0.001% P85, P105 or F68, the results being comparable to the negative control (0.05% DMSO). This suggests that an intravenous formulation containing P85, P105, or F68 would not lead to haemolysis at the tested concentrations, supporting the safety profile of Pluronic polymers for medical use[15][82]. In agreement, no differences were reported in the terminal haematological values (including haemoglobin, packed cell volume, number of erythrocytes, total number of leukocytes) and blood-chemical values (including urea, total protein, alkaline phosphatase) obtained from rats who had received once daily intravenous doses of F68 (doses ranging from 10-1000 mg/kg body weight) or from rats who had been administered physiological saline for one month [83]. No morphological abnormalities were detected in the rats which received the 0-50 mg/kg daily dose of F68, however, rats which received the higher doses had detectable alterations i.e. the presence of foam cells in the lungs (dose was 500-1000 mg/kg) and focal cortical degenerative changes in the kidneys (dose was 100-1000 mg/kg).

Pentamidine caused a concentration-dependent inhibition of insulin secretion from MIN6 β-cells suggesting that this is one mechanism through which it could induce diabetes[9]. Pentamidine is known to be an agonist at imidazoline receptors [84], but it is unlikely that this explains its inhibitory effects on insulin secretion since β-cell imidazoline receptors are coupled to increased insulin release[85]. However, the imidazoline ligand idazoxan is reported to cause a concentration-dependent inhibition of β-cell viability[86], similar to the effects observed here with pentamidine, so it is possible that the reduction in insulin secretion is secondary to pentamidine-mediated activation of β-cell imidazoline receptors and impairment of cell viability. Pentamidine-induced diabetes is not thought to be reversible [9], and so testing for a marker of pancreatic off target adverse effects occurred early in the screening cascade. Importantly, a number of Pluronic formulations (P85, P105) were shown to increase the peripheral toxicity of pentamidine as measured by decreases in insulin secretion. In a human tissue cell model (HEK-293), P105 has previously been shown to cause dose dependent changes in cell viability[16]. However, a lead Pluronic (F68) was identified which demonstrated equivalent toxicity to unformulated pentamidine, on β-cell viability and insulin secretion. Supporting this formulation selection our studies also revealed that P85 and P105 at 0.01% and 0.5% concentrations caused loss of MDCK-MDR monolayer integrity, whereas F68 at concentrations up to 0.5% had no effect (Fig S7). A correlation between HLB and cytotoxicity has previously been observed with low cytotoxicity being guaranteed when the HLB of the polymer is ≥10 (Table S1)[28].

Importantly, all formulations tested did not prevent pentamidine killing Trypanosoma brucei blood stream form trypomastigotes. In fact, pure P85 and P105 were highly trypanocidal and F68-pentamidine formulations had a slight synergistic effect.

*In vitro* BBB studies indicated that there was an efflux process for pentamidine as also demonstrated in P-gp knockout mice studies [7]. However, we were unable to demonstrate an increase in pentamidine movement across the barrier in either direction, compared with unformulated pentamidine in any of our *in vitro* systems.

Further studies utilizing the *in situ* brain perfusion technique confirmed that the Pluronics (P85, P105 or F68) did not increase pentamidine delivery to the brain, including the choroid plexus, after either 10 or 30 minutes exposure. Our studies using in situ brain perfusions over 10 minutes in mice have shown that the P85, P105 and F68 formulations have a tendency to actually prevent uptake of pentamidine into brain tissue and/or vascular endothelial cells, which constitute an intact BBB. This may be related to interactions of the Pluronics with influx transporters for pentamidine (e.g. OCT1), although our *in vitro* BBB studies did not indicate that the pentamidine permeability was affected by the presence of F68, P85 and P105 (0.01% and 0.1%) in either direction. Importantly, a similar P85 induced reduction in BBB permeability was observed by other workers, [87] who noted a reduction in the rate of uptake into brain tissue of P85-leptin conjugates during the first 90 minutes after iv injection compared with native leptin. Despite this initial inhibition of P85-leptin influx, a greater overall concentration of the conjugate was measured in brain tissue after 4 hours, an observation that the authors ascribed to improved pharmacokinetic properties. Digoxin delivery to the brain has previously been determined 1 - 10 hr post-injection in mice and found to be significantly enhanced when Pluronic 85 is present [88].

Sucrose does not cross phospholipid membranes and was used in the brain perfusion experiments as a vascular space marker. An increase in [^14^C(U)]sucrose would indicate that the integrity of the membrane or the tight junctions between cells had been compromised. Conversely, a decrease would suggest that the proportionate volume of tissue occupied by blood vessels had been reduced. It is therefore interesting that F68 has previously been shown to interact with the mechanisms that control vasoconstriction and vasodilation[89][90] and could lead to the observed reduction in vascular space.

Interestingly, the *in vivo* mouse pharmacokinetic study revealed that the concentrations of pentamidine in brain parenchyma in this species seem high compared with data from human (using CSF rather than brain parenchyma) which indicated that less than 1% of the plasma pentamidine concentration is detected in CSF[91]. Furthermore, assessment of this lead formulation in an *in vivo* pharmacokinetic study confrmed that F68 did not increase pentamidine delivery to the brain under the conditions studied. This is not linked to partitioning of pentamidine inside the micelles as this is low, hence the use of Pluronic micelles to protect this drug after administration and extend its circulation time is probably limited. Although it may be related to the fact that F68 is hydrophilic and prefers to remain in the plasma than be distributed to organs [17].

Whilst there are limitations to all assay systems, the package of data generated by the team provided a compelling and robust data set. The screening cascade has successfully identified Pluronic-pentamidine formulations that harbour trypanocidal activity and do not increase the safety concerns centrally or peripherally (over unformulated pentamidine). However, the data suggested that we would not be able to significantly enhance brain exposure of pentamidine using the Pluronic (F68, P85 or P105) within a reasonable time frame and existing budget. We therefore drew the study to a close at milestone 2 (Fig 1). Importantly a significant body of high-quality data has been generated as part of this project which may be highly relevant to other teams looking to understand block-copolymer architecture, further develop block-copolymers as nanocarriers, improve BBB penetration of drugs or to those looking to understand toxicity of pentamidine.

## Author Contributions for uploading during submission

**LS: PDRA** *in vivo* BBB (PK and In situ brain perfusion technique) and in vitro BBB studies. Data analysis. Result writing.

**MAS**: PDRA insulin secretion and cell viability studies, physical chemistry, data analysis result writing.

**GNS**: PhD Student *in vitro* BBB studies, haemolysis assay, membrane integrity assay. Data analysis

**RCB**: *in vitro* BBB studies.

**HBS**: PhD Student *in vitro* anti-trypanosomal assays.

**MF**: PhD Student contributed to the initial concept and data sets with P85.

**BL**: Trained MF, MAS and LS in insulin secretion assay and helped obtain preliminary data

**LAD**: Haemolysis assay and with SAT obtained funding from BBSRC Centre of Integrative Bioscience for the use of this assay in this study.

**SAT**: Introduction, method, result and discussion writing, paper collation, co-ordination. CNS drug delivery expert. Obtained MRC DPFS funding, concept, project management and experimental design, data analysis and interpretation.

**CD, CL, MV, SP, MC, SC, SAT**: obtaining MRC DPFS funding, concept, project management and experimental design, data analysis and interpretation.

## Abbreviations

aq: aqueous
BBB: blood-brain barrier
CAC: critical aggregation concentration
CMC: critical micellar concentration
CVO: circumventricular organs
CMH: Cyanmethaemoglobin
DPFS: developmental pathway funding scheme
DPD: dissipative particle dynamics
DNDi: Drugs for Neglected Diseases initiative
DLS: dynamic light-scattering
HAT: human African trypanosomiasis
HLB: hydrophilic-lipophilic balance
iv: intravenous
IR: inward rectifying
MW: molecular weight
MRP: multi-drug resistance associated protein
MRC: Medical Research Council
na: not available
ppm: parts per million
PTI: pentamidine isethionate
Pgp: P-glycoprotein
PFH: plasma free haemoglobin
PEO: poly(ethylene oxide)
PPO: poly(propylene oxide)
sal: saline
SLD: scattering length density

## Acknowledgements

This research was funded by a Medical Research Council (MRC) DPFS award (MR/K015451/1) to Dr S.A. Thomas (PI), Dr C.A. Dreiss, Dr C. Lorenz, Professor S. Persaud, Dr M. Christie (all at King’s College London), Dr V. Yardley and Professor S. Croft (both at LSHTM). This award was jointly funded by the UK MRC and the UK Department for International Development (DFID) under the MRC/DFID Concordat agreement and is also part of the EDCTP2 programme supported by the European Union. This research was also supported by the MRC (PhD studentship MR/K500811/1) and a multi-user equipment grant from the Wellcome Trust which provided a Perkin-Elmer Tricarb 2900TR liquid scintillation counter (080268). The BBSRC Centre of Integrative Biomedicine provided funding for the haemolysis assay.

We are grateful for the project management support of Dr Gayle Chapman (Biomedical Catalyst Ltd), UK. We would like to acknowledge the support of Dr Surbi Gupta (King’s IP & Licensing team) who was involved in the IP assessment of the project.

We would like to acknowledge the support of Drugs for Neglected Diseases initiative (DNDi) a collaborative, patients’ needs-driven, non-profit drug research and development organization that is developing new treatments for Neglected Diseases.

We would like to thank Miss Raha Ahmadkhanbeigi (King’s College London) for help with the haemolysis assay. We would like to thank Ms Ana-Charlott Berglar (King’s College London) for help with acquisition of the preliminary data. We would like to thank ISIS, Rutherford-Appleton Laboratory (Science and Technology Facilities Council, Didcot, Oxford) for the award of neutron beam time on the LOQ instrument to perform the SANS experiments.

## Supporting information captions

**Fig S1. Pentamidine is returned to the blood from the capillary endothelial cell by P-gp and MRP. Pluronic P85 inhibits-mediated efflux (e.g. P-gp and MRP transport) by two mechanisms; the first through membrane fluidisation and the second through transient ATP depletion.** These effects are believed to be mediated by unimers (single polymer chains) [21][19]. Inhibition of efflux should facilitate the accumulation of pentamidine in the human cerebral capillary endothelium and the murine choroid plexus epithelium, leading to higher concentrations of pentamidine.

**Fig S2. Pyrene fluorescence intensity dependence on pluronic concentration for F68, P85 and P105.** The CMC was determined using 18 different concentrations (range 0.0001 to 1 w/v%) of pure P85, P105 and F68. The value at each concentration is the mean of two samples, each prepared from a separate preparation of the stock solution. As expected the curves show two inflection points. The first was taken as the CMC.

**Fig S3. Typical partition data for PTI fluorescence as a function of F68 and P105 concentration.**

**Fig S4. Drug release from dialysis cells measured over time. The experiments were conducted in water at 37°C for concentrations as close as possible to *in vitro* conditions, within experimental limitations, namely, 1% w/v of Pluronics and 10mM PTI.** No significant differences between the Pluronics were observed and drug release is diffusion controlled (Fickian diffusion) under the experimental conditions. Pluronics micelles are not a barrier to drug release.

**Fig S5: SANS Pluronic data at 37°C.** A) P85 5% B) F68 5% C) P85 5% / PTI 1 % D) F68 5% / PTI 1 % E) P85 5% / PTI 3 % F) F68 5% / PTI 3 %.

**Fig S6. The average number of Pluronic molecules found in a micelle (N_agg_) and the number of micelles in our system (after they have equilibrated) (N_mic_) as a function of the concentration of the F68 Pluronic in a system that contains F68 and 0.01 w/v% of L61 Pluronic.** In the both plots, the black curve represents the results when considering both the L61 and F68 polymers in the mixture, and the blue dashed curve represents the data from the pure F68 simulated systems. In the top curve, the red curve represents the number of F68 in a micelle which contains both F68 and L61, and the green curve represents the number of L61 in a micelle. The results show that as we increase the concentration of F68, and therefore make the system more and more like the pure F68 system, the number of polymer molecules in a micelle and the number of micelles converge to that observed in the pure F68 system, as expected. Interestingly, it seems that from our simulations that L61 causes the aggregation of F68 to become slightly enhanced as the number of F68 in the average micelle is always larger than that found in the pure F68 micelles, which naturally results in their being fewer micelles.

**Fig S7. Apical to basolateral permeability of [^14^C]sucrose in the presence of P85, P105, and F68 concentrations measured over 60 minutes.** Significant differences compared to control (no pluronic) was observed in the presence of P85 and P105 (***p<0.001, **p<0.01). All data are expressed as mean ± S.E.M, n= 3 wells. Data were analysed using one-way ANOVA with SigmaPlot 13.0.

**Fig S8. Effects of exposure of MIN6 β-cells to 0 (control), 1 or 100 μM pentamidine for 3 and 24 hours. Trypan blue uptake.** Blue staining demonstrates cells of compromised viability, highlighting the toxicity of 100 μM pentamidine to these cells after 3 hours exposure.

**Fig S9. Effects of exposure of MIN6 β-cells to 0, 1, 10 or 100 μM pentamidine and 0, 0.01, 0.025, 0.1 or 0.5% w/v% F68 for 24 hours.** Trypan blue uptake. Blue staining demonstrates cells of compromised viability, highlighting the toxicity of 100 μM pentamidine and 0.5% F68 to these cells.

**Table S1. Single point CNS screening of pentamidine at a concentration of 1.0E M (PE study no. 13-9625).** Values are expressed as the percent inhibition of specific binding and represent the average of replicate tubes. Bolder values represent inhibition of 50% or greater.

**Table S2. Inhibition of hKir2.1 potassium channel activity with pentamidine isethionate.**

**Evaluated by the QPatch HT an automatic parallel patch clamp system.** The duration of exposure to each test concentration was 3 minutes.

**Table S3. A visual evaluation of the phase separation of Pluronics dispersions in pure water. Transparent is fully transparent. Opaque completely blocks light.** Slight indicates for slightly translucent (faintly white tint in the solution), and medium indicates obvious translucence.

**Table S4. A visual evaluation of the phase separation of pluronics dispersions in saline. Transparent is fully transparent.** Opaque completely blocks light. Slight indicates for slightly translucent (faintly white tint in the solution), and medium indicates obvious translucence

**Table S5. The effect of P85, F68 and P105 on the apparent permeability of pentamidine isethionate across MDR1-MDCK cell monolayers in the basolateral to apical direction.** The apical to basolateral movement of pentamidine isethionate was below the limits of detection. The percentage recovery of pentamidine isethionate is also shown. Lucifer yellow permeation was below 0.5 × 10^−6^ cm/s in all experiments confirming the integrity of the monolayer. Transcellular marker (propranolol) and Pgp and BCRP substrate (prazosin) apparent permeability values are also shown.

**Table S6. The effect of Pluronic P85 on the accumulation of [^3^H(G)]pentamidine (15.7 nM) into brain tissues after 10 minutes of in situ perfusion.** All values have been corrected for vascular space by subtraction of the R_TISSUE_ % for [^14^C(U)]sucrose from the R_TISSUE_ % for [^3^H(G)]pentamidine. All values mean ± SEM.

**Table S7. The effect of Pluronic P105 on the accumulation of [^3^H(G)]pentamidine (15.7 nM) into brain parenchyma after 10 minutes of in situ perfusion.** All values have been corrected for vascular space by subtraction of the R_TISSUE_% for [^14^C(U)] sucrose from the R_TISSUE_% for [^3^H(G)]pentamidine.

**Table S8. Accumulation of H-pentamidine (15.7 nM) after 10 minutes perfusion with or without pluronic F68 (not corrected for vascular space; Control A and 0.01% and 0.1% F68 experiments were carried out using MP Biomedicals dextran.** Control B and 0.5% F68 experiments were carried out using VWR dextran).

**Table S9. Accumulation of **[^14^C]sucrose** after 10 minutes perfusion with or without Pluronic F68; Control A and 0.01% and 0.1% F68 experiments were carried out using MP Biomedicals dextran.** Control B and 0.5% F68 experiments were carried out using VWR dextran).

**Table S10A. Accumulation of [^3^H]pentamidine after 30 minutes perfusion with or without pluronic F68.** (Not corrected for vascular space).

**Table S10B. Accumulation of [^14^C]sucrose (B) after 30 minutes perfusion with or without pluronic F68.** (Not corrected for vascular space).

## References

1. Simarro PP, Diarra A, Postigo JAR, Franco JR, Jannin JG. The human african trypanosomiasis control and surveillance programme of the world health organization 2000-2009: The way forward. PLoS Negl Trop Dis. 2011;5.

2. WHO. No Title. In: http://www.who.int/trypanosomiasis_african/country/en/. 2014.

3. Franco JR, Simarro PP, Diarra A, Jannin JG. Epidemiology of human African trypanosomiasis. Clin Epidemiol. 2014;6: 257–75. doi:10.2147/CLEP.S39728

4. Sanderson L, Dogruel M, Rodgers J, Bradley B, Thomas SA. The blood-brain barrier significantly limits eflornithine entry into Trypanosoma brucei brucei infected mouse brain. J Neurochem. 2008;107: 1136–1146.

5. Jeganathan S, Sanderson L, Dogruel M, Rodgers J, Croft S, Thomas SA. The distribution of nifurtimox across the healthy and trypanosome-infected murine blood-brain and blood-cerebrospinal fluid barriers. J Pharmacol Exp Ther. 2011;336: 506–515.

6. Sanderson L, Khan A, Thomas S. Distribution of suramin, an antitrypanosomal drug, across the blood-brain and blood-cerebrospinal fluid interfaces in wild-type and P-glycoprotein transporter-deficient mice. Antimicrob Agents Chemother. 2007;51: 3136–3146.

7. Sanderson L, Dogruel M, Rodgers J, Koning HP De, Thomas SA. Pentamidine movement across the murine blood-brain and blood-cerebrospinal fluid barriers: Effect of trypanosome infection, combination therapy, P-glycoprotein, and multidrug resistance-associated protein. J Pharmacol Exp Ther. 2009;329: 967–971.

8. Doua F, Miezan TW, Singaro JRS, Yapo FB, Baltz T. The efficacy of pentamidine in the treatment of early-late stage Trypanosoma brucei gambiense trypanosomiasis. Am J Trop Med Hyg. 1996;55: 586–588.

9. Burri C. Chemotherapy against human African trypanosomiasis: is there a road to success? Parasitology. 2010;137: 1987–94. doi:10.1017/S0031182010001137

10. Batrakova E V, Li S, Li Y, Alakhov VY, Elmquist WF, Kabanov A V. Distribution kinetics of a micelle-forming block copolymer Pluronic P85. J Control Release. 2004;100: 389–97. doi:10.1016/j.jconrel.2004.09.002

11. Kozlov MY, Melik-Nubarov NS, Batrakova E V., Kabanov A V. Relationship between Pluronic Block Copolymer Structure, Critical Micellization Concentration and Partitioning Coefficients of Low Molecular Mass Solutes. Macromolecules. American Chemical Society; 2000;33: 3305–3313. doi:10.1021/ma991634x

12. Cagel M, Tesan FC, Bernabeu E, Salgueiro MJ, Zubillaga MB, Moretton MA, et al. Polymeric mixed micelles as nanomedicines: Achievements and perspectives. Eur J Pharm Biopharm. 2017;113: 211–228. doi:10.1016/j.ejpb.2016.12.019

13. Oh KT, Bronich TK, Kabanov A V. Micellar formulations for drug delivery based on mixtures of hydrophobic and hydrophilic Pluronic^®^ block copolymers. J Control Release. 2004;94: 411–422. doi:10.1016/j.jconrel.2003.10.018

14. Chiappetta D.A. and Sosnik A. Poly(ethylene oxide)–poly(propylene oxide) block copolymer micelles as drug delivery agents: Improved hydrosolubility, stability and bioavailability of drugs. Eur J Pharm Biopharm. 2007;66: 303.

15. Singh-Joy SD, McLain VC. Safety assessment of poloxamers 101, 105, 108, 122, 123, 124, 181, 182, 183, 184, 185, 188, 212, 215, 217, 231, 234, 235, 237, 238, 282, 284, 288, 331, 333, 334, 335, 338, 401, 402, 403, and 407, poloxamer 105 benzoate, and poloxamer 182 dibenzoate as use. Int J Toxicol. 2008;27 Suppl 2: 93–128. doi:10.1080/10915810802244595

16. Fang X, Chen, Sha X, Zhang, Zhong, Fan, et al. Pluronic mixed micelles overcoming methotrexate multidrug resistance: in vitro and in vivo evaluation. Int J Nanomedicine. 2013; 1463. doi:10.2147/IJN.S42368

17. Alakhov VY, Kabanov A V. Block copolymeric biotransport carriers as versatile vehicles for drug delivery. Expert Opin Investig Drugs. 1998;7: 1453–1473. doi:10.1517/13543784.7.9.1453

18. Shaik N, Pan G, Elmquist WF. Interactions of pluronic block copolymers on P-gp efflux activity: Experience with HIV-1 protease inhibitors. J Pharm Sci. 2008;97: 5421–5433. doi:10.1002/jps.21372

19. Batrakova E V., Han H-Y, Alakhov VY, Miller DW, Kabanov A V. Effects of Pluronic Block Copolymers on Drug Absorption in Caco-2 Cell Monolayers. Pharm Res. Kluwer Academic Publishers-Plenum Publishers; 1998;15: 850–855. doi:10.1023/A:1011964213024

20. Batrakova E, Lee S, Li S, Venne A, Alakhov V, Kabanov A. Fundamental Relationships Between the Composition of Pluronic Block Copolymers and Their Hypersensitization Effect in MDR Cancer Cells. Pharm Res. 1999;16: 1373–1379. doi:10.1023/A:1018942823676

21. Shaik N, Giri N, Elmquist WF. Investigation of the micellar effect of pluronic P85 on P-glycoprotein inhibition: cell accumulation and equilibrium dialysis studies. J Pharm Sci. 2009;98: 4170–90. doi:10.1002/jps.21723

22. Miller DW, Batrakova E V., Kabanov A V. Inhibition of Multidrug Resistance-Associated Protein (MRP) Functional Activity with Pluronic Block Copolymers. Pharm Res. Kluwer Academic Publishers-Plenum Publishers; 1999;16: 396–401. doi:10.1023/A:1018873702411

23. Rapoport N, Marin AP, Timoshin AA. Effect of a Polymeric Surfactant on Electron Transport in HL-60 Cells. Arch Biochem Biophys. 2000;384: 100–108. doi:10.1006/abbi.2000.2104

24. Batrakova E V. Optimal Structure Requirements for Pluronic Block Copolymers in Modifying P-glycoprotein Drug Efflux Transporter Activity in Bovine Brain Microvessel Endothelial Cells. J Pharmacol Exp Ther. 2003;304: 845–854. doi:10.1124/jpet.102.043307

25. Wang Y, Hao J, Li Y, Zhang Z, Sha X, Han L, et al. Poly(caprolactone)-modified Pluronic P105 micelles for reversal of paclitaxcel-resistance in SKOV-3 tumors. Biomaterials. 2012;33: 4741–51. doi:10.1016/j.biomaterials.2012.03.013

26. Venne A, Li S, Mandeville R, Kabanov A, Alakhov V. Hypersensitizing effect of pluronic L61 on cytotoxic activity, transport, and subcellular distribution of doxorubicin in multiple drug-resistant cells. Cancer Res. 1996;56: 3626–3629.

27. Huang J, Si L, Jiang L, Fan Z, Qiu J, Li G. Effect of pluronic F68 block copolymer on P-glycoprotein transport and CYP3A4 metabolism. Int J Pharm. 2008;356: 351–3. doi:10.1016/j.ijpharm.2007.12.028

28. Budkina OA, Demina T V., Dorodnykh TY, Melik-Nubarov NS, Grozdova ID. Cytotoxicity of nonionic amphiphilic copolymers. Polym Sci Ser A. 2012;54: 707–717. doi:10.1134/S0965545X12080020

29. Valle JW, Armstrong A, Newman C, Alakhov V, Pietrzynski G, Brewer J, et al. A phase 2 study of SP1049C, doxorubicin in P-glycoprotein-targeting pluronics, in patients with advanced adenocarcinoma of the esophagus and gastroesophageal junction. Invest New Drugs. 2011;29: 1029–1037.

30. Alakhova DY, Zhao Y, Li S, Kabanov A V. Effect of Doxorubicin/Pluronic SP1049C on Tumorigenicity, Aggressiveness, DNA Methylation and Stem Cell Markers in Murine Leukemia. PLoS One. 2013;8. doi:10.1371/journal.pone.0072238

31. Mendonça DVC, Lage LMR, Lage DP, Chávez-Fumagalli MA, Ludolf F, Roatt BM, et al. Poloxamer 407 (Pluronic^®^ F127)-based polymeric micelles for amphotericin B: In vitro biological activity, toxicity and in vivo therapeutic efficacy against murine tegumentary leishmaniasis. Exp Parasitol. 2016;169: 34–42. doi:10.1016/j.exppara.2016.07.005

32. Alakhova DY, Kabanov A V. Pluronics and MDR reversal: an update. Mol Pharm. 2014;11: 2566–78. doi:10.1021/mp500298q

33. Zhang K, Khan A. Phase Behavior of Poly(ethylene oxide)-Poly(propylene oxide)-Poly(ethylene oxide) Triblock Copolymers in Water. Macromolecules. 1995;28: 3807–3812. doi:10.1021/ma00115a009

34. World Health Organization. Stability testing of active pharmaceutical ingredients and finished pharmaceutical products. WHO Tech Rep Ser. 2009;953: 87–130. Available: http://www.ich.org/fileadmin/Public_Web_Site/ICH_Products/Guidelines/Quality/Q1F/Stability_Guideline_WHO.pdf

35. Fu Y, Kao WJ. Drug release kinetics and transport mechanisms of non-degradable and degradable polymeric delivery systems. Expert Opin Drug Deliv. 2010;7: 429–444. doi:10.1517/17425241003602259

36. Gaitano GG, Brown W, Tardajos G. Inclusion Complexes between Cyclodextrins and Triblock Copolymers in Aqueous Solution: A Dynamic and Static Light-Scattering Study. J Phys Chem B. 1997;101: 710–719. doi:10.1021/jp961996w

37. Chiappetta DA, Sosnik A. Poly(ethylene oxide)–poly(propylene oxide) block copolymer micelles as drug delivery agents: Improved hydrosolubility, stability and bioavailability of drugs. Eur J Pharm Biopharm. 2007;66: 303–317. doi:10.1016/j.ejpb.2007.03.022

38. Heenan, R K; King, S.M., Osborn, R., Stanley HB. COLETTE Users Guide, Rutherford Appleton Laboratory Report. Rutherford Applet Lab Rep. 1989; 89–128.

39. King, S.M., Heenan RK. Using COLETTE. Rutherford Applet Lab Rep. 1995;95.

40. SasView [Internet]. [cited 23 Nov 2014]. Available: http://www.sasview.org/

41. Warren PB. Dissipative particle dynamics. Curr Opin Colloid Interface Sci. 1998;3: 620–624. doi:10.1016/S1359-0294(98)80089-7

42. Chen S, Guo C, Hu G-H, Liu H-Z, Liang X-F, Wang J, et al. Dissipative particle dynamics simulation of gold nanoparticles stabilization by PEO–PPO–PEO block copolymer micelles. Colloid Polym Sci. 2007;285: 1543–1552. doi:10.1007/s00396-007-1721-x

43. Neun BW, Dobrovolskaia MA. Method for analysis of nanoparticle hemolytic properties in vitro. Methods Mol Biol. 2011;697: 215–24. doi:10.1007/978-1-60327-198-1_23

44. Persaud SJ, Roderigo-Milne HM, Squires PE, Sugden D, Wheeler-Jones CPD, Marsh PJ, et al. A key role for beta-cell cytosolic phospholipase A(2) in the maintenance of insulin stores but not in the initiation of insulin secretion. Diabetes. 2002;51: 98–104. Available: http://www.ncbi.nlm.nih.gov/pubmed/11756328

45. Jones PM, Salmon DM, Howell SL. Protein phosphorylation in electrically permeabilized islets of Langerhans. Effects of Ca2+, cyclic AMP, a phorbol ester and noradrenaline. Biochem J. 1988;254: 397–403. Available: http://www.ncbi.nlm.nih.gov/pubmed/2845950

46. Persaud, S.J., Al-Majed, H., Raman, A. JPM. Gymnema sylvestre stimulates insulin release in vitro by increased membrane permeability. J Endocrinol. 1999;163: 207–212. doi:10.1677/joe.0.1630207

47. Thomas née Williams SA, Segal MB. Identification of a saturable uptake system for deoxyribonucleosides at the blood-brain and blood-cerebrospinal fluid barriers. Brain Res. 1996;741: 230–239. doi:10.1016/S0006-8993(96)00930-4

48. Williams SA, Abbruscato TJ, Hruby VJ, Davis TP. Passage of a delta-opioid receptor selective enkephalin, [D-penicillamine2,5] enkephalin, across the blood-brain and the blood-cerebrospinal fluid barriers. J Neurochem. 1996;66: 1289–99. Available: http://www.ncbi.nlm.nih.gov/pubmed/8769896

49. Sanderson L, Khan A, Thomas S. Distribution of suramin, an antitrypanosomal drug, across the blood-brain and blood-cerebrospinal fluid interfaces in wild-type and P-glycoprotein transporter-deficient mice. Antimicrob Agents Chemother. 2007;51: 3136–3146.

50. Thomas née Williams SA, Segal MB. Identification of a saturable uptake system for deoxyribonucleosides at the blood-brain and blood-cerebrospinal fluid barriers. Brain Res. 1996;741: 230–239. doi:10.1016/S0006-8993(96)00930-4

51. Triguero D, Buciak J, Pardridge WM. Capillary depletion method for quantification of blood-brain barrier transport of circulating peptides and plasma proteins. J Neurochem. 1990;54: 1882–8.

52. Räz B, Iten M, Grether-Bühler Y, Kaminsky R, Brun R. The Alamar Blue assay to determine drug sensitivity of African trypanosomes (T.b. rhodesiense and T.b. gambiense) in vitro. Acta Trop. 1997;68: 139–47. Available: http://www.ncbi.nlm.nih.gov/pubmed/9386789

53. Miézan TW, Bronner U, Doua F, Cattand P, Rombo L. Long-term exposure of Trypanosoma brucei gambiense to pentamidine in vitro. Trans R Soc Trop Med Hyg. 88: 332–3. Available: http://www.ncbi.nlm.nih.gov/pubmed/7974681

54. Wood DH, Hall JE, Rose BG, Tidwell RR. 1,5-Bis(4-amidinophenoxy)pentane (pentamidine) is a potent inhibitor of []idazoxan binding to imidazoline I2 binding sites. Eur J Pharmacol. 1998;353: 97–103. doi:10.1016/S0014-2999(98)00386-0

55. De Boer T, Nalos L, Stary A, Kok B, Houtman M, Antoons G, et al. The anti-protozoal drug pentamidine blocks KIR2.x-mediated inward rectifier current by entering the cytoplasmic pore region of the channel. Br J Pharmacol. 2010;159: 1532–1541. doi:10.1111/j.1476-5381.2010.00658.x

56. Nalos L, de Boer TP, Houtman MJC, Rook MB, Vos MA, van der Heyden MAG. Inhibition of lysosomal degradation rescues pentamidine-mediated decreases of K(IR)2.1 ion channel expression but not that of K(v)11.1. Eur J Pharmacol. 2011;652: 96–103. doi:10.1016/j.ejphar.2010.10.093

57. Williams K, Dattilo M, Sabado TN, Kashiwagi K, Igarashi K. Pharmacology of delta2 glutamate receptors: effects of pentamidine and protons. J Pharmacol Exp Ther. 2003;305: 740–8. doi:10.1124/jpet.102.045799

58. Cox D, Motoyama Y, Seki J, Aoki T, Dohi M YK. Pentamidine: a non-peptide GPIIb/IIIa antagonist--in vitro studies on platelets from humans and other species. Thromb Haemost. 1992;68: 731–736.

59. Reynolds IJ, Aizenman E. Pentamidine is an N-methyl-D-aspartate receptor antagonist and is neuroprotective in vitro. J Neurosci. 1992;12: 970–5. Available: http://www.ncbi.nlm.nih.gov/pubmed/1532027

60. Pathak MK, Dhawan D, Lindner DJ, Borden EC, Farver C, Yi T. Pentamidine is an inhibitor of PRL phosphatases with anticancer activity. Mol Cancer Ther. 2002;1: 1255–64. Available: http://www.ncbi.nlm.nih.gov/pubmed/12516958

61. Kitamura Y, Arima T, Imaizumi R, Sato T, Nomura Y. Inhibition of constitutive nitric oxide synthase in the brain by pentamidine, a calmodulin antagonist. Eur J Pharmacol. 1995;289: 299–304. Available: http://www.ncbi.nlm.nih.gov/pubmed/7542607

62. Chen X, Qiu L, Li M, D??rrnagel S, Orser BA, Xiong ZG, et al. Diarylamidines: High potency inhibitors of acid-sensing ion channels. Neuropharmacology. 2010;58: 1045–1053. doi:10.1016/j.neuropharm.2010.01.011

63. Perilo CS, Pereira MT, Santoro MM, Nagem RAP. Structural binding evidence of the trypanocidal drugs berenil and pentacarinate active principles to a serine protease model. Int J Biol Macromol. 2010;46: 502–11. doi:10.1016/j.ijbiomac.2010.03.006

64. Cimecioglu AL, Weiss RA. Aromatic Polyamides of 3,8-Diamino-6-phenylphenanthridine and Their Molecular Complexes with Sulfonated Polystyrene Ionomers. Macromolecules. 1995;28: 6343–6346. doi:10.1021/ma00122a047

65. Mortensen K, Brown W. Poly(ethylene oxide)-poly(propylene oxide)-poly(ethylene oxide) triblock copolymers in aqueous solution. The influence of relative block size. Macromolecules. 1993;26: 4128–4135. doi:10.1021/ma00068a010

66. Hammouda B. SANS from Pluronic P85 in d-water. Eur Polym J. 2010;46: 2275–2281. doi:10.1016/j.eurpolymj.2010.10.012

67. Valero M, Castiglione F, Mele A, da Silva MA, Grillo I, González-Gaitano G, et al. Competitive and Synergistic Interactions between Polymer Micelles, Drugs, and Cyclodextrins: The Importance of Drug Solubilization Locus. Langmuir. 2016;32: 13174–13186. doi:10.1021/acs.langmuir.6b03367

68. Puig-Rigall J, Obregon-Gomez I, Monreal-Pérez P, Radulescu A, Blanco-Prieto MJ, Dreiss CA, et al. Phase behaviour, micellar structure and linear rheology of tetrablock copolymer Tetronic 908. J Colloid Interface Sci. 2018;524: 42–51. doi:10.1016/j.jcis.2018.03.096

69. Wanka G, Hoffmann H, Ulbricht W. Phase Diagrams and Aggregation Behavior of Poly(oxyethylene)-Poly(oxypropylene)-Poly(oxyethylene) Triblock Copolymers in Aqueous Solutions. Macromolecules. 1994;27: 4145–4159. doi:10.1021/ma00093a016

70. Alexandridis P, Alan Hatton T. Poly(ethylene oxide)ltl poly(propylene oxide)ltl poly(ethylene oxide) block copolymer surfactants in aqueous solutions and at interfaces: thermodynamics, structure, dynamics, and modeling. Colloids Surfaces A Physicochem Eng Asp. 1995;96: 1–46. doi:10.1016/0927-7757(94)03028-X

71. Alexandridis P, Holzwarth JF, Hatton TA. Micellization of Poly(ethylene oxide)-Poly(propylene oxide)-Poly(ethylene oxide) Triblock Copolymers in Aqueous Solutions: Thermodynamics of Copolymer Association. Macromolecules. 1994;27: 2414–2425. doi:10.1021/ma00087a009

72. Alexandridis P, Athanassiou V, Fukuda S, Hatton TA. Surface Activity of Poly(ethylene oxide)-block-Poly(propylene oxide)-block-Poly(ethylene oxide) Copolymers. Langmuir. 1994;10: 2604–2612. doi:10.1021/la00020a019

73. Lopes JR, Loh W. Investigation of Self-Assembly and Micelle Polarity for a Wide Range of Ethylene Oxide−Propylene Oxide−Ethylene Oxide Block Copolymers in Water. Langmuir. 1998;14: 750–756. doi:10.1021/la9709655

74. Booth C, Attwood D. Effects of block architecture and composition on the association properties of poly(oxyalkylene) copolymers in aqueous solution. Macromol Rapid Commun. 2000;21: 501–527. doi:10.1002/1521-3927(20000601)21:9<501::AID-MARC501>3.0.CO;2-R

75. Pepić I, Lovrić J, Hafner A, Filipović-Grčić J. Powder form and stability of Pluronic mixed micelle dispersions for drug delivery applications. Drug Dev Ind Pharm. 2014;40: 944–951. doi:10.3109/03639045.2013.791831

76. Wanka G, Hoffmann H, Ulbricht W. The aggregation behavior of poly-(oxyethylene)-poly-(oxypropylene)-poly-(oxyethylene)-block-copolymers in aqueous solution. Colloid Polym Sci. 1990;268: 101–117. doi:10.1007/BF01513189

77. Kositza MJ, Bohne C, Alexandridis P, Hatton TA, Holzwarth JF. Micellization Dynamics and Impurity Solubilization of the Block-Copolymer L64 in an Aqueous Solution. Langmuir. 1999;15: 322–325. doi:10.1021/la9812368

78. Loh W. Block and Copolymer Micelles. In: Somasundaram, P., Dekker M, editor. Encyclopedia of Surface and Colloid Science. 2006. pp. 1014–1025.

79. Prasad KN, Luong TT, FlorenceJoelle Paris AT, Vaution C, Seiller M, Puisieux F. Surface activity and association of ABA polyoxyethylene— polyoxypropylene block copolymers in aqueous solution. J Colloid Interface Sci. 1979;69: 225–232. doi:10.1016/0021-9797(79)90151-6

80. Sahoo L, Sarangi J, Misra PK. Organization of Amphiphiles, Part 1: Evidence in Favor of Pre-micellar Aggregates through Fluorescence Spectroscopy. Bull Chem Soc Jpn. 2002;75: 859–865. doi:10.1246/bcsj.75.859

81. Kabanov A V., Nazarova IR, Astafieva I V., Batrakova E V., Alakhov VY, Yaroslavov AA, et al. Micelle Formation and Solubilization of Fluorescent Probes in Poly(oxyethylene-b-oxypropylene-b-oxyethylene) Solutions. Macromolecules. American Chemical Society; 1995;28: 2303–2314. doi:10.1021/ma00111a026

82. Lowe KC, Furmidge BA, Thomas S. Haemolytic Properties of Pluronic Surfactants and Effects of Purification. Artif Cells, Blood Substitutes, Biotechnol. 1995;23: 135–139. doi:10.3109/10731199509117673

83. Magnusson G, Olsson T, Nyberg JA. Toxicity of Pluronic F-68. Toxicol Lett. 1986;30: 203–7. Available: http://www.ncbi.nlm.nih.gov/pubmed/3705104

84. Morgan N, Chan S. Imidazoline Binding Sites in the Endocrine Pancreas: Can They Fulfil Their Potential as Targets for the Development of New Insulin Secretagogues? Curr Pharm Des. 2001;7: 1413–1431. doi:10.2174/1381612013397366

85. Morgan NG. Imidazoline receptors: new targets for antihyperglycaemic drugs. Expert Opin Investig Drugs. 1999;8: 575–84. doi:10.1517/13543784.8.5.575

86. Gao H, Mourtada M, Morgan NG. Effects of the imidazoline binding site ligands, idazoxan and efaroxan, on the viability of insulin-secreting BRIN-BD11 cells. JOP. 2003;4: 117–24. Available: http://www.ncbi.nlm.nih.gov/pubmed/12743417

87. Yi X, Yuan D, Farr SA, Banks WA, Poon C-D, Kabanov A V. Pluronic modified leptin with increased systemic circulation, brain uptake and efficacy for treatment of obesity. J Control Release. 2014;191: 34–46. doi:10.1016/j.jconrel.2014.05.044

88. Batrakova E V, Miller DW, Li S, Alakhov VY, Kabanov A V, Elmquist WF. Pluronic P85 enhances the delivery of digoxin to the brain: in vitro and in vivo studies. J Pharmacol Exp Ther. 2001;296: 551–7. Available: http://www.ncbi.nlm.nih.gov/pubmed/11160643

89. Saeed M, Hartmann A, Bing RJ. Inhibition of vasoactive agents by perfluorochemical emulsion. Life Sci. 1987;40: 1971–9. Available: http://www.ncbi.nlm.nih.gov/pubmed/2437426

90. Watanabe M, Okada T. Lysophosphatidylcholine-induced myocardial damage is inhibited by pretreatment with poloxamer 188 in isolated rat heart. Mol Cell Biochem. 2003;248: 209–15. Available: http://www.ncbi.nlm.nih.gov/pubmed/12870676

91. Bronner U, Doua F, Ericsson Ö, Gustafsson LL, Miézan T, Rais M, et al. Pentamidine concentrations in plasma, whole blood and cerebrospinal fluid during treatment of Trypanosoma gambiense infection in Côte d’Ivoire. Trans R Soc Trop Med Hyg. 1991;85: 608–611. doi:10.1016/0035-9203(91)90364-5

