## Supplementary file for "Drug reformulation for a neglected disease. The NANOHAT project to develop a safer more effective sleeping sickness drug"

**Full title:**

Drug reformulation for a neglected disease.

**Short title:**

Pentamidine reformulation to improve safety and efficacy against Human African Trypanosomiasis

**AUTHORS:** Lisa Sanderson^1†^, Marcelo da Silva^1†^, Gayathri N. Sekhar^1^, Rachel C. Brown^1^, Hollie Burrell-Saward^2^, Mehmet Fidanboylu^1^, Bo Liu^3^, Lea Ann Dailey^1^, Cécile A. Dreiss^1^, Chris Lorenz^4^, Mark Christie^1^, Shanta J. Persaud^3^, Vanessa Yardley^2^, Simon L Croft^2^, Margarita Valero^5^ and Sarah A. Thomas^1^*

^1^King’s College London, Institute of Pharmaceutical Science, Stamford Street, London, UK

^2^Faculty of Infectious and Tropical Diseases, London School of Hygiene and Tropical Medicine, London, UK.

^3^King's College London, Department of Diabetes, School of Life Course Sciences, Faculty of Life Sciences & Medicine, 2.9N Hodgkin Building, Guy's campus, London UK.

^4^ King's College London, Theory & Simulation of Condensed Matter Group, Department of Physics, Strand, London WC2R 2LS, UK

^5^Physical Chemistry Department, Faculty of Pharmacy, University of Salamanca, Spain.

†joint first authors

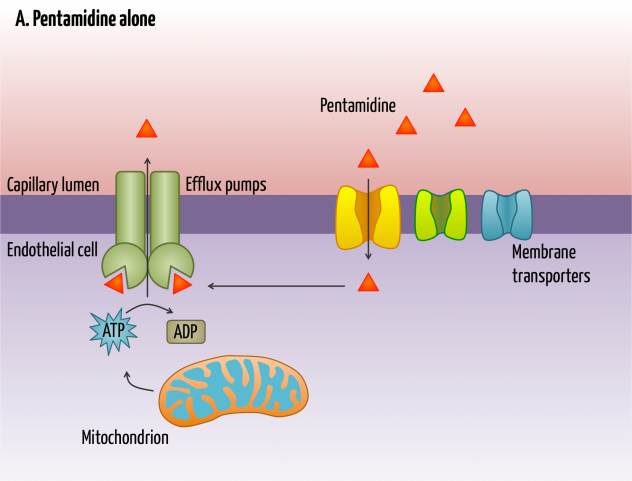

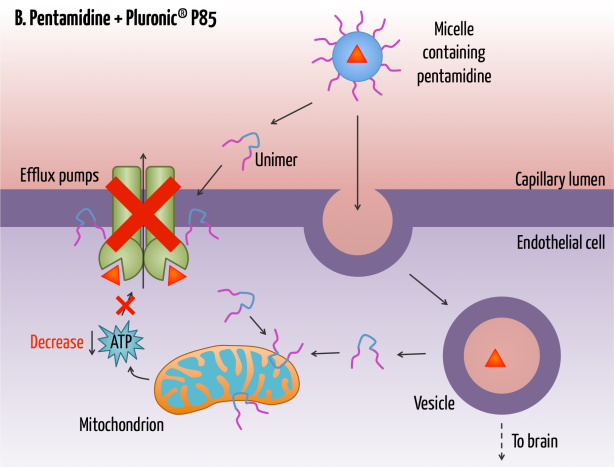

**Fig S1.** **Pentamidine is returned to the blood from the capillary endothelial cell by P-gp and MRP. Pluronic^®^P85 inhibits-mediated efflux (e.g. P-gp and MRP transport) by two mechanisms; the first through membrane fluidisation and the second through transient ATP depletion**. These effects are believed to be mediated by unimers (single polymer chains) (Shaik *et al.* 2009)(Batrakova *et al.* 1998). Inhibition of efflux should facilitate the accumulation of pentamidine in the human cerebral capillary endothelium and the murine choroid plexus epithelium, leading to higher concentrations of pentamidine.

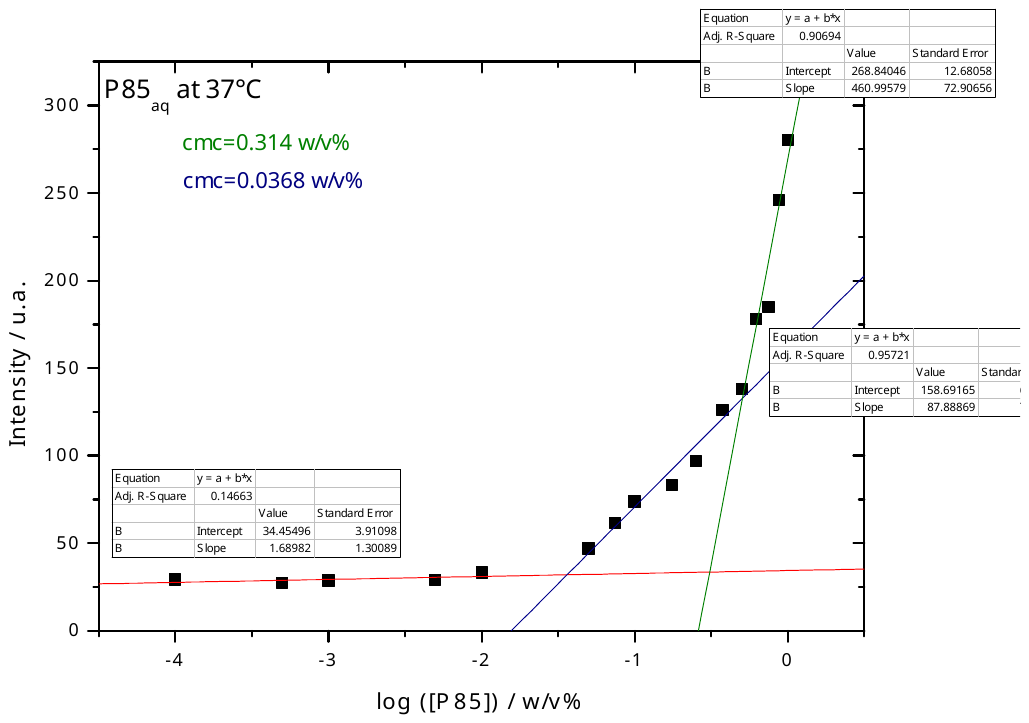

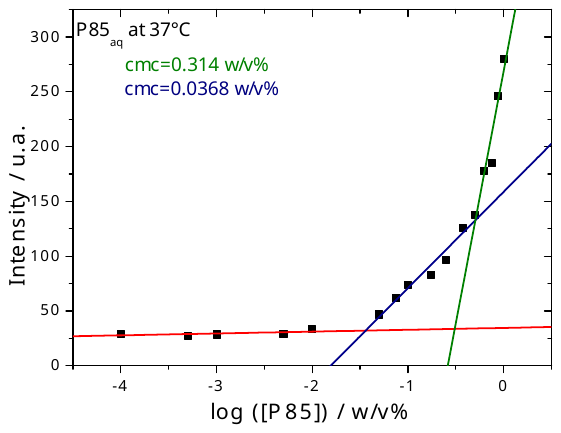

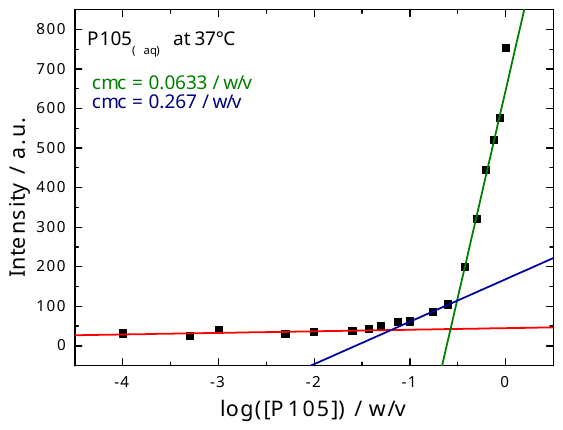

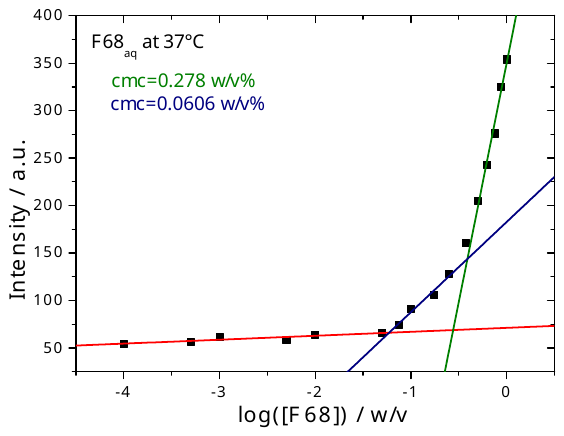

**Fig S2**. **Pyrene fluorescence intensity dependence on pluronic concentration for F68, P85 and P105**. The CMC was determined using 18 different concentrations (range 0.0001 to 1 w/v%) of pure P85, P105 and F68. The value at each concentration is the mean of two samples, each prepared from a separate preparation of the stock solution.  As expected the curves show two inflection points. The first was taken as the CMC.

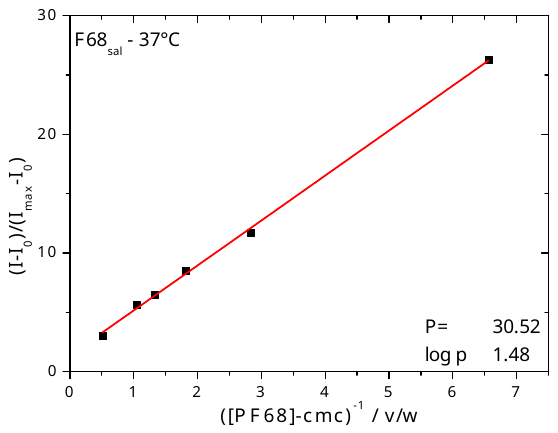

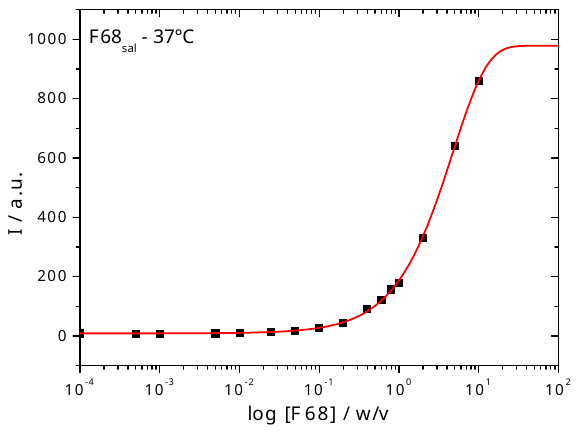

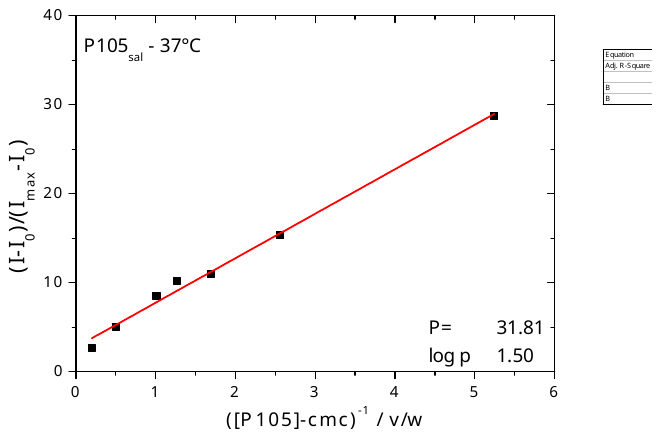

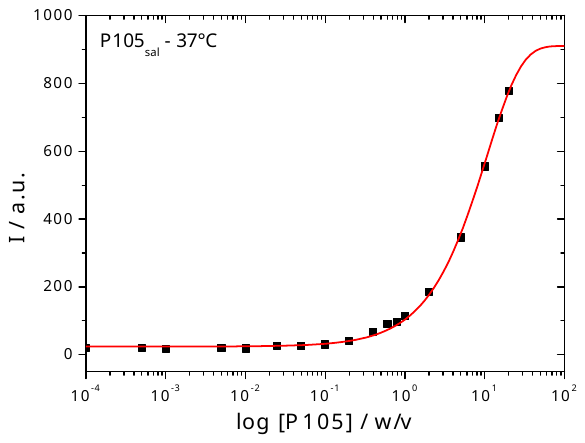

**Fig S3.** **Typical partition data for PTI fluorescence as a function of F68 and P105 concentration.**

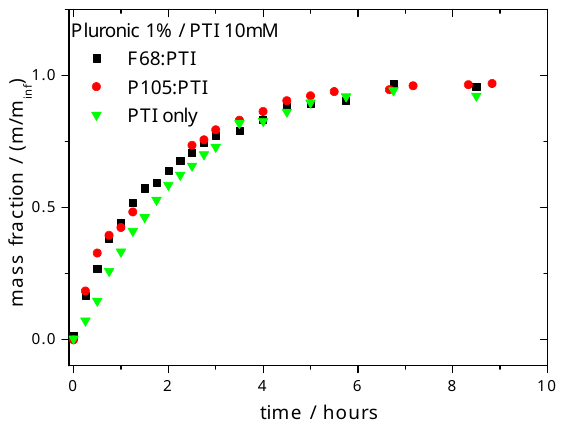

**Fig S4. Drug release from dialysis cells measured over time. The experiments were conducted in water at 37°C for concentrations as close as possible to *in vitro* conditions, within experimental limitations, namely, 1% w/v of Pluronics and 10mM PTI**. No significant differences between the Pluronics were observed and drug release is diffusion controlled (Fickian diffusion) under the experimental conditions. Pluronics micelles are not a barrier to drug release.

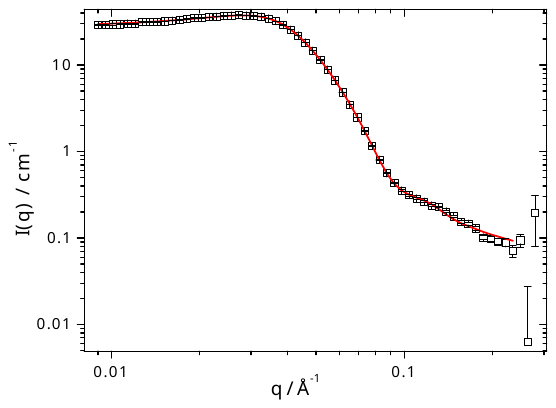

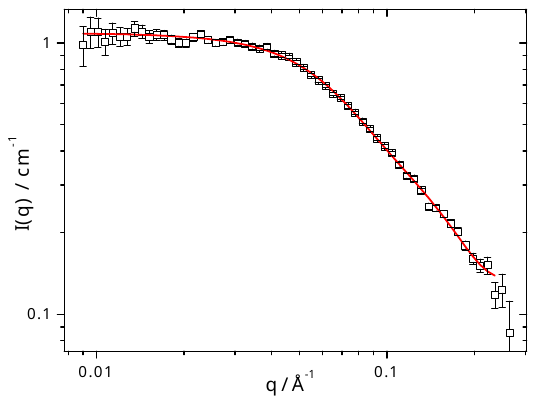

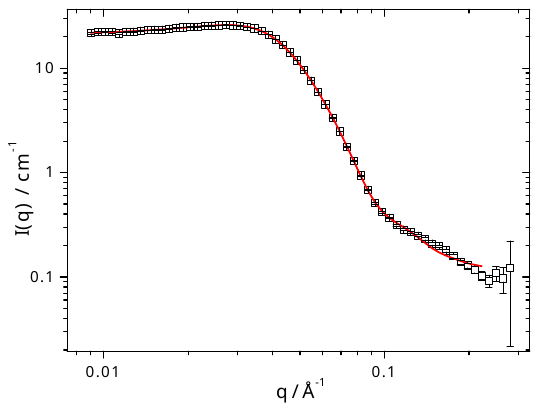

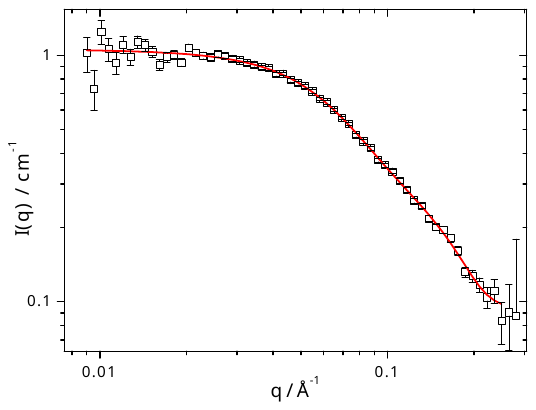

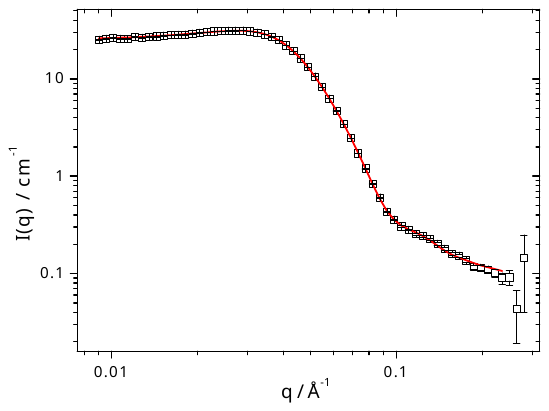

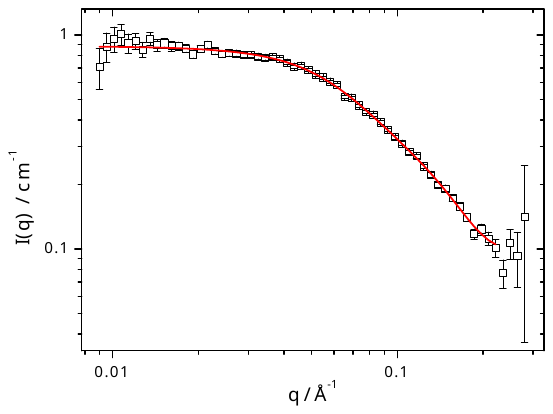
**Fig S5.** **SANS Pluronic data at 37°C.** A) P85 5% B) F68 5% C) P85 5% / PTI 1 % D) F68 5% / PTI 1 % E) P85 5% / PTI 3 % F) F68 5% / PTI 3 %.

D

E

F

B

A

C

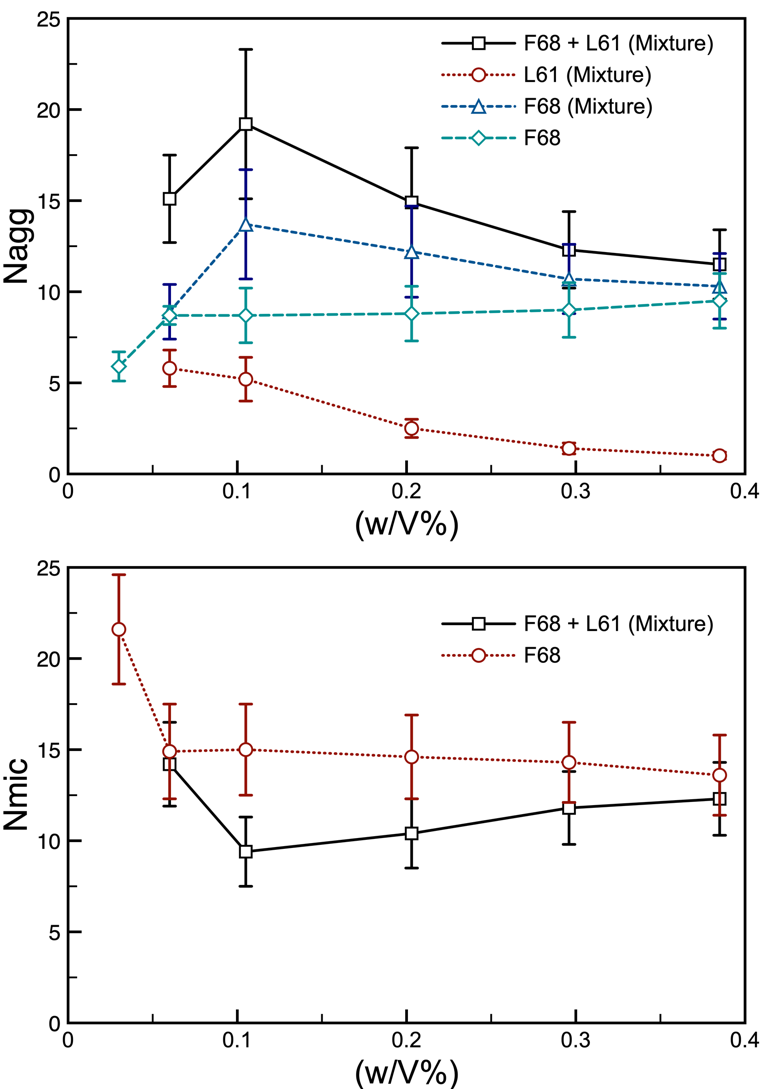

**Fig S6.** **The average number of Pluronic molecules found in a micelle (N_agg_) and the number of micelles in our system (after they have equilibrated) (N_mic_) as a function of the concentration of the F68 Pluronic in a system that contains F68 and 0.01 w/v% of L61 Pluronic**. In the both plots, the black curve represents the results when considering both the L61 and F68 polymers in the mixture, and the blue dashed curve represents the data from the pure F68 simulated systems. In the top curve, the red curve represents the number of F68 in a micelle which contains both F68 and L61, and the green curve represents the number of L61 in a micelle. The results show that as we increase the concentration of F68, and therefore make the system more and more like the pure F68 system, the number of polymer molecules in a micelle and the number of micelles converge to that observed in the pure F68 system, as expected. Interestingly, it seems that from our simulations that L61 causes the aggregation of F68 to become slightly enhanced as the number of F68 in the average micelle is always larger than that found in the pure F68 micelles, which naturally results in their being fewer micelles.

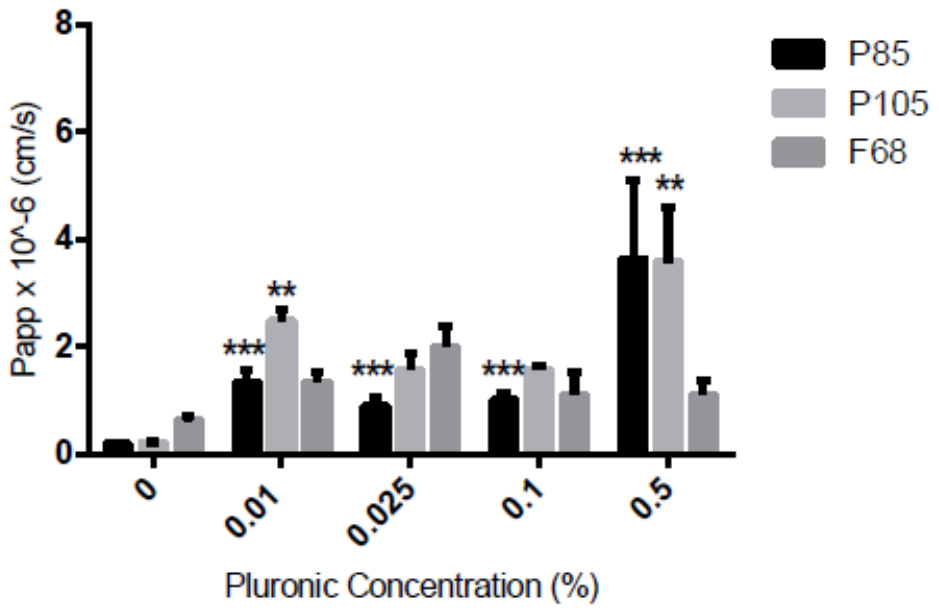

**Fig S7.** **Apical to basolateral permeability of [^14^C]sucrose in the presence of P85, P105, and F68 concentrations measured over 60 minutes**. Significant differences compared to control (no pluronic) was observed in the presence of P85 and P105 (***p<0.001, **p<0.01). All data are expressed as mean ± S.E.M, n= 3 wells. Data were analysed using one-way ANOVA with SigmaPlot 13.0.

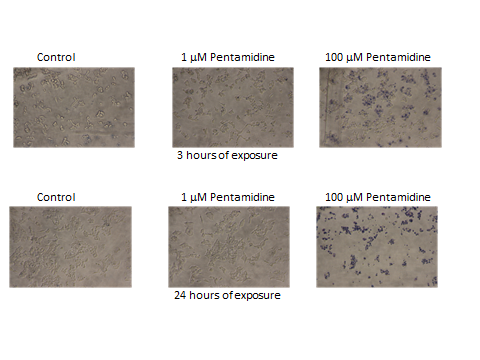

**Fig S8.** **Effects of exposure of MIN6 β-cells to 0 (control), 1 or 100 μM pentamidine for 3 and 24 hours. Trypan blue uptake.** Blue staining demonstrates cells of compromised viability, highlighting the toxicity of 100 μM pentamidine to these cells after 3 hours exposure.

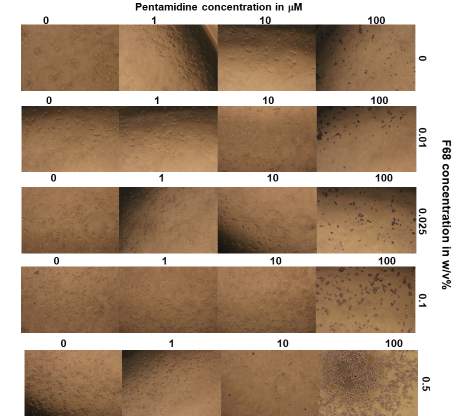

**Fig S9.** **Effects of exposure of MIN6 β-cells to 0, 1, 10 or 100 μM pentamidine and 0, 0.01, 0.025, 0.1 or 0.5% w/v% F68 for 24 hours**. Trypan blue uptake. Blue staining demonstrates cells of compromised viability, highlighting the toxicity of 100 μM pentamidine and 0.5% F68 to these cells.

**Table S1. Single point CNS screening of pentamidine at a concentration of 1.0E^-5^ M (PERKIN ELMER study no. 13-9625).** Values are expressed as the percent inhibition of specific binding and represent the average of replicate tubes. Bolder values represent inhibition of 50% or greater.

| **RECEPTOR** | **Percent Inhibition (%)**  **Mean**  **n = 2** | **RECEPTOR** | **Percent Inhibition (%)**  **Mean**  **n = 2** |
| --- | --- | --- | --- |
| **NEUROTRANSMITTER RELATED** |  | **Opioid Non-selective** | **76.00** |
| Adenosine Non-selective | 1.28 | Opioid, Orphanin, ORL1 (h) | 36.62 |
| **Adrenergic, Alpha 1, Non-selective** | **100.77** | **Serotonin Transporter** | **61.05** |
| **Adrenergic, Alpha 2, Non-selective** | **56.14** | Serotonin, Non-selective | 0.09 |
| **Adrenergic, Beta, Non-selective** | **55.37** | Sigma, Non-selective | 6.50 |
| **Dopamine Transporter** | **75.89** | **ION CHANNELS** |  |
| Dopamine, D1 | 27.48 | Potassium Channel, ATP-sensitive | 6.07 |
| Dopamine, D2s | 20.12 | Potassium Channel,  Ca ^2+^ Act., VI | 2.07 |
| GABA A, Agonist Site | 31.10 | Sodium, Site 2 | **46.94** |
| GABA A, BDZ, alpha 1 site | 20.54 | **SECOND MESSENGERS** |  |
| GABA-B | 10.74 | Nitric Oxide, NOS (Neuronal binding) | 12.98 |
| Glutamate, AMPA Site (Ionotropic) | 1.34 | **GROWTH FACTORS/HORMONES** |  |
| Glutamate, Kainate Site (Ionotropic | -13.63 | Corticotropin Releasing Factor, Non-selective | 17.05 |
| Glutamate, MK-801 site (Ionotropic) | 6.25 | **BRAIN/GUT PEPTIDES** |  |
| Glutamate, NMDA Agonist Site (Ionotropic) | 5.99 | Angiotensin II, AT2 | 2.96 |
| Glutamate, NMDA, Glycine (Strychnine-insensitive Site) (Ionotropic) | -17.52 | Cholecystokinin, CCK2 (CCKB) | 0.09 |
| Glycine (Strychnine-sensitive Site) | -0.22 | Endothelin, ET-B (h) | 4.50 |
| Histamine, H1 | -1.9 | **ENZYMES** |  |
| **Histamine, H2** | **71.07** | Decarboxylase,  Glutamic Acid | -1.41 |
| **Imidazoline, I2 , Central** | **101.03** | Esterase, Acetylcholine | 23.21 |
| Melatonin, Non-selective | 23.45 | **Oxidase, MAO-A-Peripheral** | **89.42** |
| **Muscarinic, Non-selective, Central** | **90.90** | **Oxidase MAO-B Peripheral** | **99.99** |
| Nicotonic, Neuronal (a-BnTx insensitive) | 10.87 |  |  |
| Norepinephrine Transporter | 21.39 |  |  |

**Table S2. Inhibition of hKir2.1 potassium channel activity with pentamidine isethionate.**

**Evaluated by the QPatch HT an automatic parallel patch clamp system.** The duration of exposure to each test concentration was 3 minutes.

| Concentration  (μM) | % hKir2.1 Inhibition  Mean (± SEM) % | Number of cells  n |
| --- | --- | --- |
| 0.001 | -2.9 (± 1.7) | 5 |
| 0.01 | 1.2 (± 1.4) | 5 |
| 0.1 | 1.4 (± 1.3) | 5 |
| 1 | 3.0 (± 2.7) | 5 |
| 10 | 12.3 (± 1.3) | 4 |

**Table S3.** **A visual evaluation of the phase separation of Pluronics dispersions in pure water. Transparent is fully transparent. Opaque completely blocks light.** Slight indicates for slightly translucent (faintly white tint in the solution), and medium indicates obvious translucence.

| **P105**  **Wt%** | **F68**  **Wt%** | **L61**  **Wt%** | **Temperature**  **25°C** | **Temperature**  **30°C** | **Temperature**  **37°C** | **Temperature**  **40°C** | **Temperature**  **45°C** | **Temperature**  **50°C** |
| --- | --- | --- | --- | --- | --- | --- | --- | --- |
| 5 | 0 | 0.5 | Transparent | Transparent | Transparent | Transparent | Transparent | Transparent |
| 0 | 5 | 0.5 | Transparent | medium | opaque | opaque | opaque | opaque |
| 5 | 5 | 0.5 | Transparent | Transparent | Transparent | Transparent | Transparent | Transparent |
| 5 | 0 | 0.1 | Transparent | Transparent | Transparent | Transparent | Transparent | Transparent |
| 0 | 5 | 0.1 | Transparent | Transparent | slight | slight | slight | slight |
| 5 | 5 | 0.1 | Transparent | Transparent | Transparent | Transparent | Transparent | Transparent |
| 2.5 | 0 | 0.5 | Transparent | Transparent | Transparent | Transparent | Transparent | Transparent |
| 0 | 2.5 | 0.5 | Transparent | Medium | Opaque | Opaque | Opaque | Opaque |
| 2.5 | 2.5 | 0.5 | Transparent | Transparent | Transparent | Transparent | Transparent | Transparent |
| 2.5 | 0 | 0.1 | Transparent | Transparent | Transparent | Transparent | Transparent | Transparent |
| 0 | 2.5 | 0.1 | Transparent | Transparent | Slight | Medium | Medium | Medium |
| 2.5 | 2.5 | 0.1 | Transparent | Transparent | Transparent | Transparent | Transparent | Transparent |
| 1 | 0 | 0.1 | Transparent | Transparent | Transparent | Transparent | Transparent | Transparent |
| 0 | 1 | 0.1 | Transparent | Transparent | Medium | Opaque | Opaque | Opaque |
| 1 | 1 | 0.1 | Transparent | Transparent | Transparent | Transparent | Transparent | Transparent |
| 0.5 | 0 | 0.1 | Transparent | Transparent | Transparent | Transparent | Transparent | Transparent |
| 0 | 0.5 | 0.1 | Transparent | Transparent | Medium | Medium | Medium | Medium |
| 0.5 | 0.5 | 0.1 | Transparent | Transparent | Transparent | Transparent | Transparent | Transparent |
| 0.2 | 0 | 0.1 | Transparent | Transparent | Medium | Medium | Medium | Medium |
| 0 | 0.2 | 0.1 | Transparent | Transparent | Medium | Medium | Medium | Medium |
| 0.2 | 0.2 | 1 | Transparent | Transparent | Medium | Medium | Medium | Medium |
| 0 | 0 | 0.5 | Transparent | Opaque | Opaque | Opaque | Opaque | Opaque |
| 0 | 0 | 0.1 | Transparent | Slight | Opaque | Opaque | Opaque | Opaque |

**Table S4.** **A visual evaluation of the phase separation of pluronics dispersions in saline. Transparent is fully transparent**. Opaque completely blocks light. Slight indicates for slightly translucent (faintly white tint in the solution), and medium indicates obvious translucence.

| **P105**  **Wt%** | **F68**  **Wt%** | **L61**  **Wt%** | **Temperature**  **25°C** | **Temperature**  **30°C** | **Temperature**  **37°C** | **Temperature**  **40°C** | **Temperature**  **45°C** | **Temperature**  **50°C** |
| --- | --- | --- | --- | --- | --- | --- | --- | --- |
| 5 | 0 | 0.5 | Transparent | Transparent | Transparent | Transparent | Transparent | Transparent |
| 0 | 5 | 0.5 | Transparent | Opaque | Opaque | Opaque | Opaque | Opaque |
| 5 | 5 | 0.5 | Transparent | Transparent | Transparent | Transparent | Transparent | Transparent |
| 5 | 0 | 0.1 | Transparent | Transparent | Transparent | Transparent | Transparent | Transparent |
| 0 | 5 | 0.1 | Transparent | Transparent | Slight | Slight | Slight | Slight |
| 5 | 5 | 0.1 | Transparent | Transparent | Transparent | Transparent | Transparent | Transparent |
| 2.5 | 0 | 0.5 | Transparent | Transparent | Transparent | Transparent | Transparent | Transparent |
| 0 | 2.5 | 0.5 | Transparent | Opaque | Opaque | Opaque | Opaque | Opaque |
| 2.5 | 2.5 | 0.5 | Transparent | Transparent | Transparent | Transparent | Transparent | Transparent |
| 2.5 | 0 | 0.1 | Transparent | Transparent | Transparent | Transparent | Transparent | Transparent |
| 0 | 2.5 | 0.1 | Transparent | Transparent | Medium | Medium | Medium | Medium |
| 2.5 | 2.5 | 0.1 | Transparent | Transparent | Transparent | Transparent | Transparent | Transparent |
| 1 | 0 | 0.1 | Transparent | Transparent | Transparent | Transparent | Transparent | Transparent |
| 0 | 1 | 0.1 | Transparent | Slight | Medium | Opaque | Opaque | Opaque |
| 1 | 1 | 0.1 | Transparent | Transparent | Transparent | Transparent | Transparent | Transparent |
| 0.5 | 0 | 0.1 | Transparent | Transparent | Transparent | Transparent | Transparent | Transparent |
| 0 | 0.5 | 0.1 | Transparent | Slight | Medium | Opaque | Opaque | Opaque |
| 0.5 | 0.5 | 0.1 | Transparent | Transparent | Transparent | Transparent | Transparent | Transparent |
| 0.2 | 0 | 0.1 | Transparent | Transparent | Medium | Medium | Medium | Medium |
| 0 | 0.2 | 0.1 | Transparent | Slight | Opaque | Opaque | Opaque | Opaque |
| 0.2 | 0.2 | 1 | Transparent | Transparent | Medium | Medium | Medium | Medium |
| 0 | 0 | 0.5 | Medium | Opaque | Opaque | Opaque | Opaque | Opaque |
| 0 | 0 | 0.1 | Transparent | Opaque | Opaque | Opaque | Opaque | Opaque |

**Table S5**. **The effect of P85, F68 and P105 on the apparent permeability of pentamidine isethionate across MDR1-MDCK cell monolayers in the basolateral to apical direction.** The apical to basolateral movement of pentamidine isethionate was below the limits of detection. The percentage recovery of pentamidine isethionate is also shown. Lucifer yellow permeation was below 0.5 x 10^-6^ cm/s in all experiments confirming the integrity of the monolayer. Transcellular marker (propranolol) and Pgp and BCRP substrate (prazosin) apparent permeability values are also shown.

| Drug | Concentration (%) | P_app_ A2B (10^-6^ cm/s) | | P_app_ B2A (10^-6^ cm/s) | | Mass balance A2B (%) | Mass balance B2A (%) |
| --- | --- | --- | --- | --- | --- | --- | --- |
|  |  | Replicate 1 | Replicate 2 | Replicate 1 | Replicate 2 |  |  |
| 10 μM propranolol^1^ | - | 34.1 | 28.0 | 41.4 | 41.6 | 81.5 | 84.4 |
| 10 μM prazosin^2^ | - | 2.48 | 1.96 | 69.5 | 70.7 | 92.0 | 92.0 |
| 10 μM pentamidine | - | - | - | 0.336 | 0.357 | 74.5 | 72.7 |
| 10 μM pentamidine | 0.01% P85 | - | - | 0.237 | 0.206 | 83.9 | 86.0 |
| 10 μM pentamidine | 0.1% P85 | - | - | 0.344 | 0.343 | 82.4 | 106.0 |
| 10 μM pentamidine | 0.5% P85 | - | - | 0.443 | 0.433 | 88.5 | 96.9 |
| 10 μM pentamidine | 0.01% P105 | - | - | 0.265 | 0.272 | 84.8 | 91.4 |
| 10 μM pentamidine | 0.1% P105 | - | - | 0.221 | 0.237 | 80.6 | 82.5 |
| 10 μM pentamidine | 0.5% P105 | - | - | 0.337 | 0.391 | 87.0 | 89.8 |
| 10 μM pentamidine | 0.01% F68 | - | - | 0.211 | 0.233 | 86.6 | 87.9 |
| 10 μM pentamidine | 0.1% F68 | - | - | 0.314 | 0.410 | 89.4 | 89.4 |
| 10 μM pentamidine | 0.5% F68 | - | - | 0.233 | 0.186 | 89.4 | 93.9 |

^1^ Transcellular maker that diffuses through cell membranes (Artursson 1990).
^2^ Pgp and BCRP substrate (Cisternino 2004)(Enokizono *et al.* 2008).

**Table S6. The effect of Pluronic P85 on the accumulation of [^3^H(G)]pentamidine (15.7 nM) into brain tissues after 10 minutes of *in situ* perfusion.** All values have been corrected for vascular space by subtraction of the R_TISSUE_% for [^14^C(U)]sucrose from the R_TISSUE_% for [^3^H(G)]pentamidine. All values mean ± SEM.

| **Region** | **R_CORR TISSUE_%** | | | |
| --- | --- | --- | --- | --- |
|  | **[^3^H]pentamidine (n=8)** | **[^3^H]pentamidine**  **+ 0.01% P85 (n=8)** | **[^3^H]pentamidine**  **+ 0.1% P85 (n=5)** | **[^3^H]pentamidine**  **+ 0.5% P85  (n=8)** |
| **frontal cortex** | 4.29 (± 0.93) | 1.99 (± 0.18) | 2.31 (± 0.92) | 4.91 (± 0.98) |
| **caudate putamen** | 4.02 (± 1.03) | 1.58 (± 0.19) | 0.94 (± 0.92) | 2.81 (± 0.58) |
| **occipital cortex** | 4.03 (± 0.98) | 1.80 (± 0.34) | 1.93 (± 0.92) | 4.02 (± 1.03) |
| **hippocampus** | 4.37 (± 0.97) | 1.25 (± 0.33) | 0.94 (± 0.92) | 3.28 (± 1.06) |
| **corpus callosum** | 6.73 (± 2.65) | 4.77 (± 1.90) | 3.40 (± 1.51) | 3.71 (± 1.30) |
| **hypothalamus** | 5.56 (± 1.58) | 4.54 (± 0.87) | 1.82 (± 0.92) | 8.71 (± 2.49) |
| **thalamus** | 4.76 (± 0.91) | 1.42 (± 0.33) | 1.38 (± 0.92) | 5.13 (± 1.12) |
| **pons** | 7.16 (± 1.75) | 3.15 (± 0.64) | 2.36 (± 0.92) | 8.16 (± 2.11) |
| **cerebellum** | 3.82 (± 0.64) | 1.59 (± 0.14) | 1.66 (± 0.92) | 4.69 (± 1.02) |
| **homogenate** | 4.61 (± 0.94) | 2.19 (± 0.40) | 1.41 (± 0.20) | 4.59 (± 1.06) |
| **supernatant** | 1.99 (± 0.45) | 0.58 (± 0.10) | 0.63 (± 0.09) | 1.64 (± 0.30) |
| **pellet** | 12.54 (± 1.74) | 8.43 (± 0.92) | 9.63 (± 3.22) | 14.13 (± 2.71) |
| **pineal gland** | 284.7 (± 81.4) | 285.8 (± 56.4) | 166.2 (± 19.9) | 372.6 (± 41.0) |
| **choroid plexus** | 490.9 (± 103.1) | 432.7 (± 84.3) | 836.3 (± 150.6) | 703.80 (± 150.6) |
| **pituitary gland** | 182.75(± 29.6) | 450.74(± 88.6) | 569.21(± 51.3) | 517.48(± 114.0) |

**Table S7.** **The effect of Pluronic P105 on the accumulation of [^3^H(G)]pentamidine (15.7 nM) into brain parenchyma after 10 minutes of *in situ* perfusion.** All values have been corrected for vascular space by subtraction of the R_TISSUE_% for [^14^C(U)] sucrose from the R_TISSUE_% for [^3^H(G)]pentamidine.

| **Region** | **R_CORR TISSUE_%** | | | |
| --- | --- | --- | --- | --- |
|  | **[^3^H]pentamidine (n=8)** | **[^3^H]pentamidine+**  **0.01% P105 (n=4)** | **[^3^H]pentamidine+**  **0.1% P105 (n=8)** | **[^3^H]pentamidine+**  **0.5% P105 (n=4)** |
| **frontal cortex** | 4.29 (± 0.93) | 3.70 (± 0.36) | 3.36 (± 0.78) | 1.96 (± 1.44) |
| **caudate putamen** | 4.02 (± 1.03) | 3.47 (± 0.61) | 2.36 (± 0.48) | 2.46 (± 0.87) |
| **occipital cortex** | 4.03 (± 0.98) | 3.95 (± 0.99) | 2.83 (± 0.49) | 1.90 (± 0.58) |
| **hippocampus** | 4.37 (± 0.97) | 3.22 (±0.92) | 3.20 (± 0.69) | 0.68 (± 0.36) |
| **corpus callosum** | 6.73 (± 2.65) | 10.79 (± 4.21) | 6.46 (± 1.10) | 3.62 (± 1.37) |
| **hypothalamus** | 5.56 (± 1.58) | 6.36 (± 2.16) | 3.95 (± 0.59) | 2.77 (± 1.04) |
| **thalamus** | 4.76 (± 0.91) | 4.98 (± 1.56) | 2.44 (± 0.35) | 4.82 (± 1.05) |
| **pons** | 7.16 (± 1.75) | 5.52 (± 2.87) | 5.83 (± 1.54) | 1.51 (± 1.43) |
| **cerebellum** | 3.82 (± 0.64) | 2.42 (± 0.44) | 3.23 (± 0.56) | 4.03 (± 0.56) |
| **homogenate** | 4.61 (± 0.94) | 2.75 (± 0.59) | 3.22 (± 0.60) | 3.33 (± 0.36) |
| **supernatant** | 1.99 (± 0.45) | 1.12 (± 0.28) | 1.25 (± 0.25) | 1.31 (± 0.20) |
| **pellet** | 12.54 (± 1.74) | 9.49 (± 0.72) | 16.73 (± 5.17) | 6.59 (± 0.74) |
| **pineal gland** | 284.7 (± 81.4) | 326.2 (± 10.5) | 266.7 (± 28.3) | 315.9 (± 17.4) |
| **choroid plexus** | 490.9 (± 103.1) | 289.1 (± 74.3) | 474.9 (± 74.6) | 402.6 (± 166.1) |
| **pituitary gland** | 182.75(± 37.5) | 425.16(± 63.9) | 437.78(± 105.4) | 458.99(± 166.1) |

**Table S8.** **Accumulation of ^3^H-pentamidine (15.7 nM) after 10 minutes perfusion with or without pluronic F68 (not corrected for vascular space; Control A and 0.01% and 0.1% F68 experiments were carried out using MP Biomedicals dextran.** Control B and 0.5% F68 experiments were carried out using VWR dextran).

| ^3^H-pentamidine | **R_TISSUE/PLASMA_%** | | | **(mean±SEM)** |  |
| --- | --- | --- | --- | --- | --- |
| **Region** | **Control A [^3^H]pentamidine (n=5)** | **[^3^H]pentamidine**  **+ 0.01% F68 (n=5)** | **[^3^H]pentamidine**  **+ 0.1% F68 (n=5)** | **Control B [^3^H]pentamidine**  **(n=5)** | **[^3^H]pentamidine**  **+ 0.5% F68 (n=5)** |
| **Frontal cortex** | 4.88 (± 1.18) | 4.48 (± 0.37) | 3.57 (± 0.63) | 6.23 (± 0.92) | 5.73 (± 0.73) |
| **Caudate putamen** | 4.08 (± 0.84) | 2.94 (± 0.28) | 2.38 (± 0.28) | 5.40 (± 1.00) | 3.19 (± 0.42) |
| **Occipital cortex** | 4.56 (±0.87) | 3.89 (± 0.54) | 3.95 (± 0.42) | 5.67 (± 0.99) | 4.18 (± 0.77) |
| **Hippocampus** | 3.48 (± 0.94) | 2.91 (± 0.26) | 2.51 (± 0.10) | 5.60 (± 1.43) | 3.50 (± 0.48) |
| **Corpus callosum** | 5.32 (± 1.47) | 4.45 (±0.48) | 3.26 (± 0.49) | 9.17 (± 2.67) | 6.21 (± 1.42) |
| **Hypothalamus** | 5.96 (± 1.23) | 4.95 (± 1.17) | 3.90 (± 0.60) | 8.14 (± 1.75) | 6.50 (± 1.14) |
| **Thalamus** | 4.34 (± 0.83) | 3.16 (± 0.38) | 3.24 (± 0.35) | 5.95 (± 0.92) | 3.96 (± 0.36) |
| **Pons** | 7.81 (± 0.90) | 8.41 (± 1.26) | 3.60 (± 0.64) | 10.72 (± 1.82) | 8.25 (± 2.14) |
| **Cerebellum** | 5.28 (± 0.56) | 4.77 (± 0.59) | 4.59 (± 0.63) | 6.54 (± 0.70) | 5.95 (± 0.74) |
| **Homogenate** | 4.03 (± 0.78) | 4.21 (± 0.34) | 4.42 (± 0.76) | 6.20 (± 0.97) | 6.03 (± 0.57) |
| **Supernatant** | 2.54 (± 0.39) | 2.39 (± 0.29) | 2.13 (± 0.40) | 3.00 (± 0.43) | 2.97 (± 0.28) |
| **Pellet** | 4.11 (± 1.24) | 7.50 (± 0.99) | 8.28 (± 2.47) | 13.19 (± 1.79) | 15.14 (± 3.19) |
| **Pineal gland** | 367.59 (± 39.97) | 210.80 (± 33.30) | 210.41 (± 48.84) | 361.76 (± 79.03) | 350.80 (± 58.07) |
| **Choroid plexus** | 385.74 (± 24.35) | 357.71 (± 58.50) | 376.66 (± 74.84) | 535.95 (± 107.86) | 526.78 (± 197.05) |
| **Pituitary gland** | 251.21 (± 66.80) | 167.83 (± 21.49) | 223.56 (± 19.43) | 203.27(± 32.85) | 497.44 (± 133.74) |

**Table S9. Accumulation of [^14^C]sucrose after 10 minutes perfusion with or without Pluronic F68; Control A and 0.01% and 0.1% F68 experiments were carried out using MP Biomedicals dextran.** Control B and 0.5% F68 experiments were carried out using VWR dextran).

| ^14^C-sucrose | **R_TISSUE/PLASMA_%** | | | **(mean±SEM)** |  |
| --- | --- | --- | --- | --- | --- |
| **Region** | **Control A [^14^C]sucrose (n=5)** | **[^14^C]sucrose**  **+0.01% F68 (n=5)** | **[^14^C]sucrose**  **+ 0.1% F68 (n=5)** | Control B **[^14^C]sucrose** (n=5) | **[^14^C]sucrose**  **+ 0.5% F68** (n=5) |
| **Frontal cortex** | 2.25 (± 0.45) | 2.10 (± 0.24) | 1.72 (± 0.33) | 1.94 (± 0.23) | 2.28 (± 0.43) |
| **Caudate putamen** | 2.01 (± 0.37) | 1.21 (± 0.21) | 1.05 (± 0.18) | 1.38 (± 0.12) | 1.17 (± 0.33) |
| **Occipital cortex** | 1.85 (± 0.38) | 1.64 (± 0.34) | 1.64 (± 0.49) | 1.64 (± 0.11) | 1.57 (± 0.41) |
| **Hippocampus** | 1.52 (± 0.37) | 1.22 (± 0.21) | 0.99 (± 0.12) | 1.23 (± 0.09) | 1.25 (± 0.29) |
| **Corpus callosum** | 2.88 (± 0.65) | 1.88 (± 0.34) | 1.38 (± 0.49) | 1.30 (± 0.32) | 1.80 (± 0.42) |
| **Hypothalamus** | 3.72 (± 0.88) | 2.04 (± 0.39) | 2.45 (± 0.53) | 2.58 (± 0.34) | 2.23 (± 0.22) |
| **Thalamus** | 1.66 (± 0.27) | 1.06 (± 0.22) | 1.12 (± 0.17) | 1.19 (± 0.14) | 1.18 (± 0.16) |
| **Pons** | 4.02 (± 0.49) | 4.23 (± 0.68) | 3.04 (± 0.91) | 3.56 (± 0.34) | 3.41 (± 0.53) |
| **Cerebellum** | 2.82 (± 0.41) | 2.50 (± 0.45) | 2.81 (± 0.83) | 2.72 (± 0.32) | 2.53 (± 0.38) |
| **Homogenate** | 1.80 (±0.34) | 1.63 (±0.31) | 1.76 (±0.54) | 1.59 (±0.15) | 1.77 (± 0.19) |
| **Supernatant** | 1.35 (±0.17) | 1.18 (±0.20) | 1.38 (±0.50) | 1.01 (±0.09) | 1.39 (± 0.26) |
| **Pellet** | 0.41 (±0.14) | 0.48 (±0.07) | 0.55 (±0.15) | 0.65 (±0.07) | 0.79 (± 0.09) |
| **Pineal gland** | 78.85 (±11.23) | 43.97 (±11.10) | 34.64 (±7.88) | 77.04 (±16.38) | 62.65 (± 6.00) |
| **Choroid plexus** | 54.64 (±15.07) | 42.36 (±10.69) | 37.01 (±8.46) | 45.06 (±7.07) | 47.70 (± 12.87) |
| **Pituitary gland** | 48.66 (±10.98) | 16.85 (±3.19) | 17.94 (±1.57) | 20.52 (±3.86) | 35.09 (± 4.82) |

**Table S10A. Accumulation of [^3^H]pentamidine after 30 minutes perfusion with or without pluronic F68**. (Not corrected for vascular space).

| **A [^3^H]pentamidine** | **R_TISSUE/PLASMA_% (mean ± SEM)** | | | |
| --- | --- | --- | --- | --- |
| **Region** | **Control  (15.7 nM pent) (n=5)** | **0.01% F68 +  (15.7 nM pent) (n=5)** | **0.1% F68 + (15.7 nM pent) (n=4)** | **0.5% F68 + (15.7 nM pent) (n=5)** |
| **Frontal cortex** | 10.35 (± 0.84) | 9.00 (± 1.47) | 11.91 (± 1.35) | 17.54 (± 3.58) |
| **Caudate putamen** | 8.65 (± 0.28) | 6.90 (± 1.58) | 10.41 (± 1.18) | 13.07 (± 3.45) |
| **Occipital cortex** | 9.52 (± 1.08) | 8.80 (± 1.47) | 12.14 (± 1.62) | 15.85 (± 2.71) |
| **Hippocampus** | 8.19 (± 0.30) | 6.05 (± 1.20) | 7.93 (± 1.20) | 10.65 (± 2.55) |
| **Corpus callosum** | 8.19 (± 1.97) | 2.02 (± 1.22) | 9.47 (± 0.94) | 15.72 (± 6.78) |
| **Hypothalamus** | 18.49 (± 4.17) | 8.90 (± 1.42) | 11.17 (± 1.42) | 18.73 (± 4.67) |
| **Thalamus** | 9.94 (± 1.95) | 7.99 (± 1.06) | 11.08 (± 1.06) | 13.52 (± 3.10) |
| **Pons** | 15.16 (± 1.48) | 17.61 (± 3.70) | 20.36 (± 3.70) | 20.43 (± 5.67) |
| **Cerebellum** | 12.92 (± 1.10) | 9.38 (± 2.47) | 14.44 (± 2.47) | 17.28 (± 3.54) |
| **Homogenate** | 10.18 (± 0.92) | 9.63 (± 0.93) | 11.19 (± 1.62) | 14.45 (± 3.93) |
| **Supernatant** | 5.54 (± 0.37) | 3.77 (± 0.94) | 6.63 (± 0.52) | 6.45 (± 1.35) |
| **Pellet** | 16.45 (± 2.76) | 18.16 (± 2.79) | 22.15 (± 4.44) | 18.42 (± 2.97) |
| **Pineal gland** | 664.25 (± 82.42) | 341.30 (± 39.57) | 342.95 (± 92.02) | 525.94 (± 122.85) |
| **Choroid plexus** | 1241.60 (± 92.86) | 835.26 (± 260.20) | 944.85 (± 235.90) | 1844.80 (± 493.51) |
| **Pituitary gland** | 394.51 (± 52.42) | 324.42 (± 44.06) | 434.10 (± 59.56) | 881.90 (± 170.25) |

**Table S10B. Accumulation of [^14^C]sucrose (B) after 30 minutes perfusion with or without pluronic F68.** (Not corrected for vascular space).

| **B ^14^C-sucrose** | **R_TISSUE/PLASMA_% (mean ± SEM)** | | | |
| --- | --- | --- | --- | --- |
| **Region** | **Control  (15.7 nM pent) (n=5)** | **0.01% F68 +  (15.7 nM pent) (n=5)** | **0.1% F68 + (15.7 nM pent) (n=4)** | **0.5% F68 + (15.7 nM pent) (n=5)** |
| **Frontal cortex** | 2.24 (± 0.29) | 5.02 (± 2.36) | 3.26 (± 0.38) | 5.42 (± 1.59) |
| **Caudate putamen** | 1.48 (± 0.18) | 4.12 (± 2.18) | 3.57 (± 0.44) | 4.02 (± 0.90) |
| **Occipital cortex** | 1.72 (± 0.12) | 3.70 (± 1.59) | 3.15 (± 0.33) | 4.26 (± 1.12) |
| **Hippocampus** | 1.32 (± 0.17) | 2.29 (± 1.22) | 2.79 (± 0.36) | 3.29 (± 1.12) |
| **Corpus callosum** | 1.67 (± 0.35) | 1.12 (± 2.52) | 4.21 (± 0.48) | 4.39 (± 2.08) |
| **Hypothalamus** | 3.50 (± 0.35) | 1.12 (± 0.99) | 4.21 (± 0.48) | 4.39 (± 2.08) |
| **Thalamus** | 1.68 (± 0.68) | 1.81 (± 1.58) | 4.21 (± 0.21) | 5.88 (± 1.04) |
| **Pons** | 4.29 (± 0.22) | 3.20 (± 2.06) | 3.09 (± 1.18) | 3.67 (± 1.49) |
| **Cerebellum** | 3.10 (± 0.26) | 5.57 (± 1.42) | 6.13 (± 0.99) | 7.21 (± 1.96) |
| **Homogenate** | 2.14 (± 0.23) | 2.60 (± 0.87) | 3.02 (± 0.52) | 3.05 (± 1.09) |
| **Supernatant** | 1.21 (± 0.17) | 2.11 (± 0.84) | 2.31 (± 0.26) | 2.53 (± 0.64) |
| **Pellet** | 1.09 (± 0.33) | 1.19 (± 0.30) | 1.43 (± 0.06) | 1.13 (± 0.18) |
| **Pineal gland** | 65.85 (± 8.08) | 30.21 (± 8.14) | 52.42 (± 4.23) | 74.63 (± 24.42) |
| **Choroid plexus** | 99.54 (± 15.32) | 66.37 (± 7.52) | 76.84 (± 13.07) | 116.86 (± 31.26) |
| **Pituitary gland** | 39.02 (± 5.27) | 18.41 (± 5.34) | 30.35 (± 5.23) | 65.21 (± 13.10) |
